# Parallel Trajectories in the Evolution of C-class MADS-box Genes in Flowering Plants and Conifers

**DOI:** 10.64898/2026.11.06.764951

**Authors:** Austin T. Nguyen, Lena C. Hileman, Kelly K. S. Matsunaga

## Abstract

Gene duplication events frequently underpin phenotypic diversity due to the increase in genetic material that can be shaped by evolution. *AGAMOUS* (*AG*) genes are members of the MADS-box gene family that show conserved function in specifying reproductive organ identity in seed plants. While the function, expression, and evolution of these genes have been extensively studied in flowering plants, they remain understudied in gymnosperms. Consequently, little is known about how *AG* has contributed to reproductive diversity and evolution outside of the flowering plants. Here, we report that *AG* genes underwent two duplication events within conifers prior to the split between Cupressaceae and its sister family, Taxaceae. Further, we characterize the *in situ* expression pattern for each resulting paralog (*AG1*, *AG2*, and *AG3*) in seed cones of three Cupressaceae species, *Metasequoia glyptostroboides*, *Taxodium distichum*, and *Juniperus virginiana*. We discover gene expression patterns consistent with subfunctionalization: *AG1* specialized in seed and sporangial development, *AG2* in seed cone and seed coat development, and *AG3* in ovuliferous scale development. This study provides an evolutionary mechanism for the development of unique seed cone reproductive traits in Cupressaceae. More broadly, our results uncover convergent patterns in the evolution of *AG* gene lineages across conifers and angiosperms.

**Significance Statement:** Gene duplications underly evolutionary innovations across the tree of life. While their role in plant evolution is widely recognized, how specific gene duplications have contributed to the evolution of important traits is poorly understood among gymnosperms, which pose significant challenges for developmental biology. We present new data from conifers on the evolution and gene expression patterns of a key regulator of reproductive development. Results help explain unique reproductive traits in conifers and provide evidence that two distantly related plant groups have taken similar paths in the evolution of these developmental regulatory genes.

## Introduction

Gene duplication can be a source of evolutionary novelty by doubling genetic material on which evolution can act and has been associated with increased morphological diversity, phenotypic innovation, and the independent evolution of complex traits in plants (1–3). After a duplication event, gene copies (paralogs) can remain functionally redundant or undergo a number of possible functional shifts: nonfunctionalization where one paralog loses functionality; subfunctionalization where ancestral functions are partitioned among paralogs; and neofunctionalization where one paralog takes on a new role (4). Sub- and neofunctionalization, in particular, can result in novelty by decoupling pleiotropic traits, allowing for the removal of trade-offs and exploration of different morphologies, or by causing new genetic programs to arise (4, 5). However, relatively little is known about how duplications of specific genes have contributed to phenotypic evolution, particularly outside of flowering plants (angiosperms), as well as whether independent duplications of the same genes share similar evolutionary fates.

The expansion of the MADS-box gene family via gene duplication is hypothesized to have driven the evolution of key reproductive innovations in plants, such as the seed and flower, and played a key role in the diversification of angiosperms (6–9). MADS-box genes encode transcription factors with diverse and critical functions in plant development and greatly increased in number as land plants diversified (10, 11). In angiosperms, the *AGAMOUS* (*AG*) subfamily of MADS-box genes performs the C function in the ABC model of flower development. The ABC model describes how the expression of different classes of genes (A, B, and C) overlap to confer floral organ identity: A alone for sepal identity, A and B for petal identity, B and C for stamen identity, and C alone for carpel identity (12). Expansions to the ABC model in angiosperms include D- and E-classes, which are involved in determining the identity of ovules and all flower organs, respectively (13, 14). Phylogenetic analyses have found that genes with D-class function are close paralogs of the angiosperm C-class genes (i.e., they are *AG* subfamily members), indicating that D function is a case of subfunctionalization from the ancestral C function (15).

In gymnosperms (non-flowering seed plants), members of the *AG* subfamily are expressed in pollen- and ovule/seed-producing structures, which suggests that *AG* genes have a conserved role in specifying reproductive organ identity across seed plants (16–26). Among gymnosperm groups, *AG* genes are best characterized in conifers. Conifers are the most diverse gymnosperm clade and, unlike many angiosperms, they produce separate pollen- and seed-bearing reproductive structures (cones). Conifer seed cones are compact shoots consisting of a stem (cone axis) and leaves (bracts), which subtend the seed-bearing organ (ovuliferous scale). Conifers exhibit significant diversity in cone structure, ranging from large and woody to single-seeded and fleshy (27), and this diversity is particularly pronounced in the cypress (Cupressaceae) and yew (Taxaceae) families, which are sister lineages to one another (2, 28). In seed cones of families Pinaceae and Podocarpaceae, the single *AG* gene is expressed in the ovuliferous scale and ovules (16, 17, 22, 24). However, in the Cupressaceae, previous work showed that expression of a single *AG* gene is restricted to the ovules and only a small portion of the subtending organ (21). These findings suggest that *AG* expression and function have diverged among major conifer lineages. However, the Cupressaceae study reported finding only one *AG* copy in the focal Cupressaceae species (21), and more recent genomic studies of some conifer taxa have uncovered multiple *AG* paralogs (29, 30). Little attention has been paid to the discovery of *AG* gene duplications in conifers, and conifer *AG* paralogs have not been placed into broader phylogenetic or developmental contexts.

In this study, we used genomic datasets to identify *AG* genes in representatives of all conifer families and analyzed the phylogenetic relationships among them to understand the evolutionary history of *AG*-lineage gene duplication events across conifer diversity. We then investigated whether *AG* genes underwent regulatory divergence in Cupressaceae and the potential molecular mechanisms underlying evolutionary changes. We report that *AG* genes underwent two duplication events prior to the divergence between Cupressaceae and its sister family, Taxaceae, and that these paralogs in Cupressaceae show expression patterns consistent with subfunctionalization within developing reproductive structures. These duplications and subsequent regulatory divergence may explain key differences between the reproductive organs of Cupressaceae and other conifers and mirror the evolutionary patterns of *AG* duplications in angiosperms.

## Results

### Multiple AG paralogs in Cupressaceae and Taxaceae

Using searches of transcriptomes generated as part of this study, as well as publicly available genomes and transcriptomes, we identified one *AG* gene in species of Pinaceae, Araucariaceae, Podocarpaceae, and Sciadopityaceae, and three *AG* genes in species of Cupressaceae and Taxaceae (*AG1*-*3* herein). In the genus *Taxus* (Taxaceae), a total of four copies were identified (*AG1*, *AG2A*, *AG2B*, and *AG3*). Our phylogenetic analysis of seed plant MADS-box genes (SI Appendix, Fig. S1) demonstrates that the three *AG* genes found in each Cupressaceae and Taxaceae species are paralogs resulting from two gene duplication events that occurred after the ancestor of Cupressaceae and Taxaceae diverged from other conifer lineages, but before the divergence of Cupressaceae from Taxaceae (Fig. 1).

**Fig. 1.**
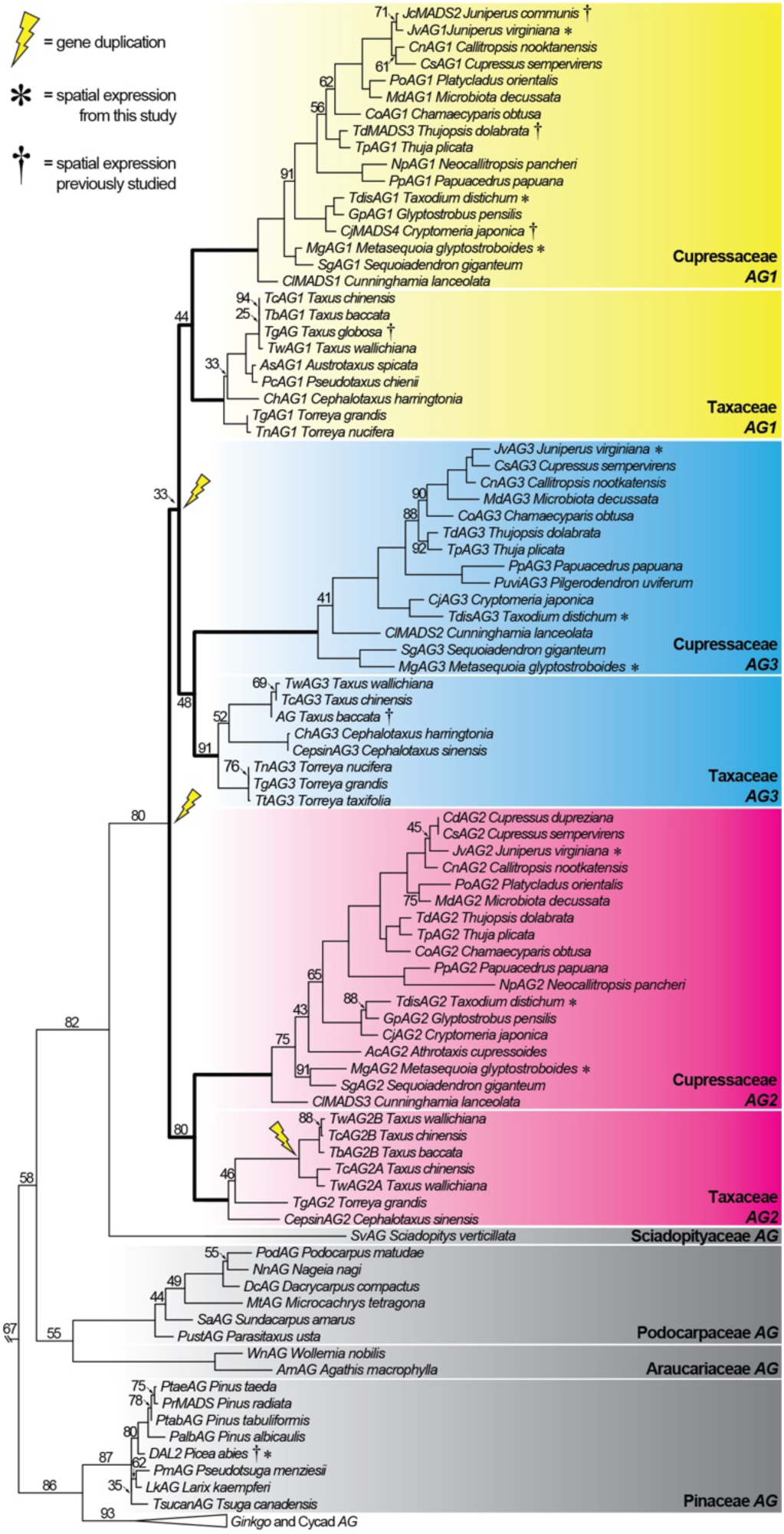
Phylogeny of *AGAMOUS* genes in conifers pruned from a larger MADS-box gene tree (marked by \\; see SI Appendix, Fig. S1 for the full MADS-box tree). Conifer families highlighted with each *AG* paralog in a different color. Numbers at nodes indicate bootstrap support. Bootstrap values ≥ 95 not shown. Lightning bolts indicate inferred gene duplication events. Bold branches indicate ones tested for positive selection. Asterisks (*) indicate genes whose spatial expression were studied in this study. Daggers (†) indicate genes whose spatial expression were previously described.

### *Patterns of* AG *expression in developing cones*

To determine in which parts of the seed cones the Cupressaceae *AG* paralogs are expressed, we conducted mRNA *in situ* hybridization (ISH) on developing cones of three Cupressaceae species: *Metasequoia glyptostroboides*, *Taxodium distichum*, and *Juniperus virginiana*. The seed cones of Cupressaceae exhibit variation in the degree of fusion between the bract and ovuliferous scale. In *T. distichum* the bract and scale are partially fused at the base, whereas in *M. glyptostroboides* and *J. virginiana*, the two structures are completely congenitally fused. In all three taxa the ovuliferous scale (seed-bearing organ) does not undergo significant growth until after pollination, so in the earliest developmental stages sampled, the ovuliferous scale is primordial in *T. distichum* and *J. virginiana*. *Metasequoia glyptostroboides* cones were at a slightly later developmental stage at the time of sampling, so the ovuliferous scale had already begun to develop.

In seed cones near pollination stage (earlier in development), the three paralogs display the following general expression patterns: *AG1* is expressed throughout the ovules and, to a lesser extent, in adjacent and subtending tissues (Fig. 2A-C; SI Appendix, Figs. S8, S12); *AG2* is expressed in almost all tissues except the nucellus (megasporangium and site of meiosis), where there is minimal to no expression (Fig. 2D–F; SI Appendix, Figs. S9, S13); *AG3* is expressed in the cone axis and the ovule-supporting organs—tissues corresponding to ovuliferous scales, or their primordia, and proximal regions of the bract where they are fused to the ovuliferous scale (Fig. 2G–I; SI Appendix, Figs. S10, S14). Based on additional assays in *M. glyptostroboides* at the stage of ovule development, the expression patterns for the three paralogs are established prior to pollination (SI Appendix, Figs. S16-19). In later stages of development, expression patterns remained consistent for *AG1* and *AG3* in all taxa, except for some *AG3* expression in the innermost layer of the integument (seed coat) in *T. distichum* and in the distal region of the nucellus in *J. virginiana* (Fig. 3; SI Appendix, Figs. S20-23). We observed some differences at later developmental stages for *AG2*. In *M. glyptostroboides* and *J. virginiana*, *AG2* expression underwent little change as seed cones developed. However, in more mature *T. distichum* seed cones, *AG2* expression became restricted primarily to the innermost region of the integument (Fig. 3D).

**Fig. 2.**
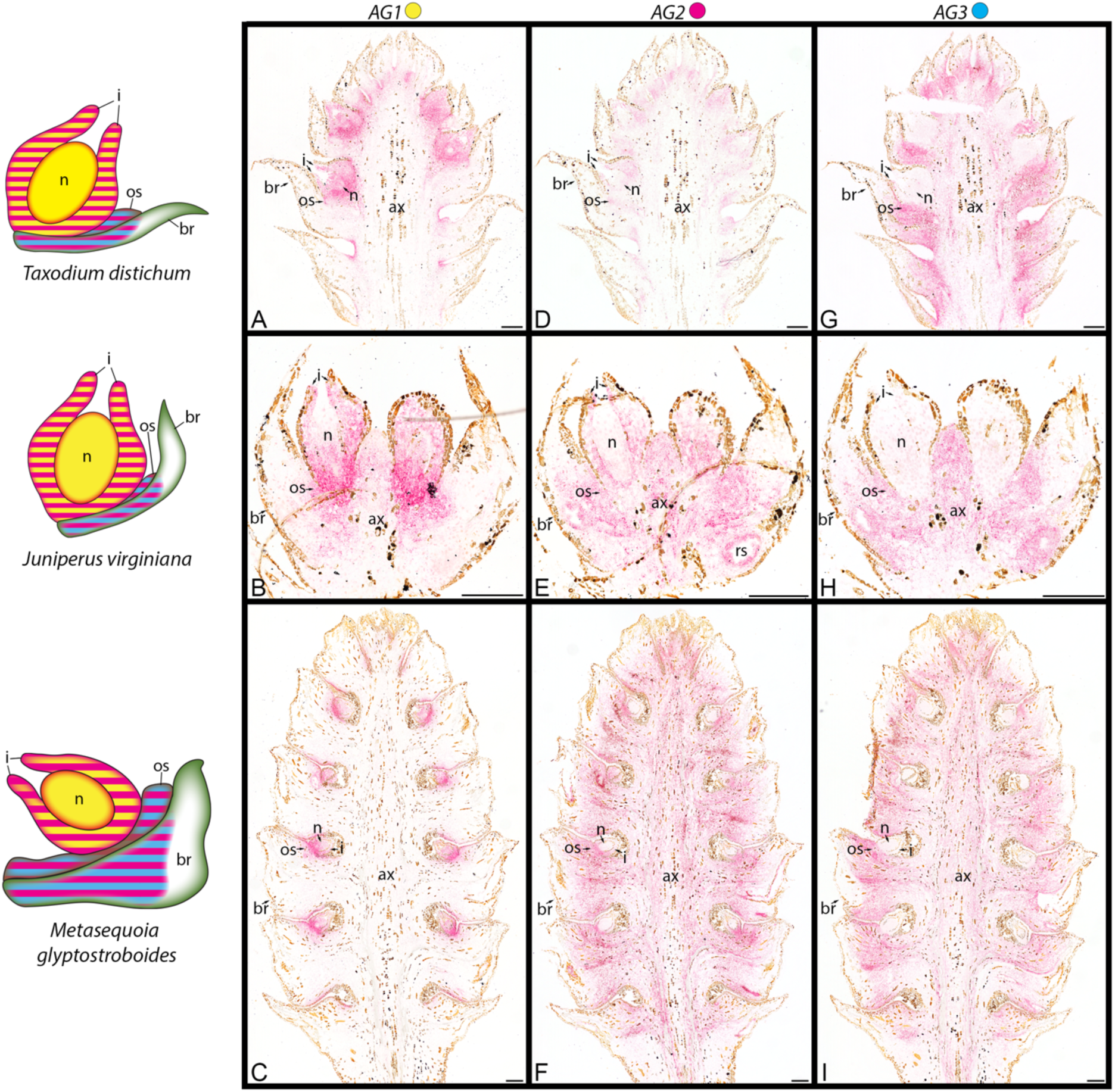
*In situ* hybridization assays of *AGAMOUS* paralogs (**[A-C]** *AG1*, **[D-F]** *AG2*, and **[G-I]** *AG3*) in seed cones near pollination stage of **(A, D, G)** *Taxodium distichum*, **(B, E, H)** *Juniperus virginiana*, and **(C, F, I)** *Metasequoia glyptostroboides*. Magenta stain indicates areas of gene expression. Note that only one ovule, ovuliferous scale, and bract are labeled for each section due to the repetitive structure of seed cones. Corresponding negative controls are found in SI Appendix, Figs. S2-4. Models at left summarize expression patterns of *AG1* (yellow), *AG2* (fuchsia), and *AG3* (cyan) throughout the ovule, ovuliferous scale, and bract in cones of each species. Regions of overlapping expression among paralogs indicated by stripes. Note that at this stage of development, the ovuliferous scale is primordial in *T. distichum* and *J. virginiana* but more developed in *M. glyptostroboides*. ax = cone axis; br = bract; os = ovuliferous scale; n = nucellus; i = integument; rs = resin canal.

**Fig. 3.**
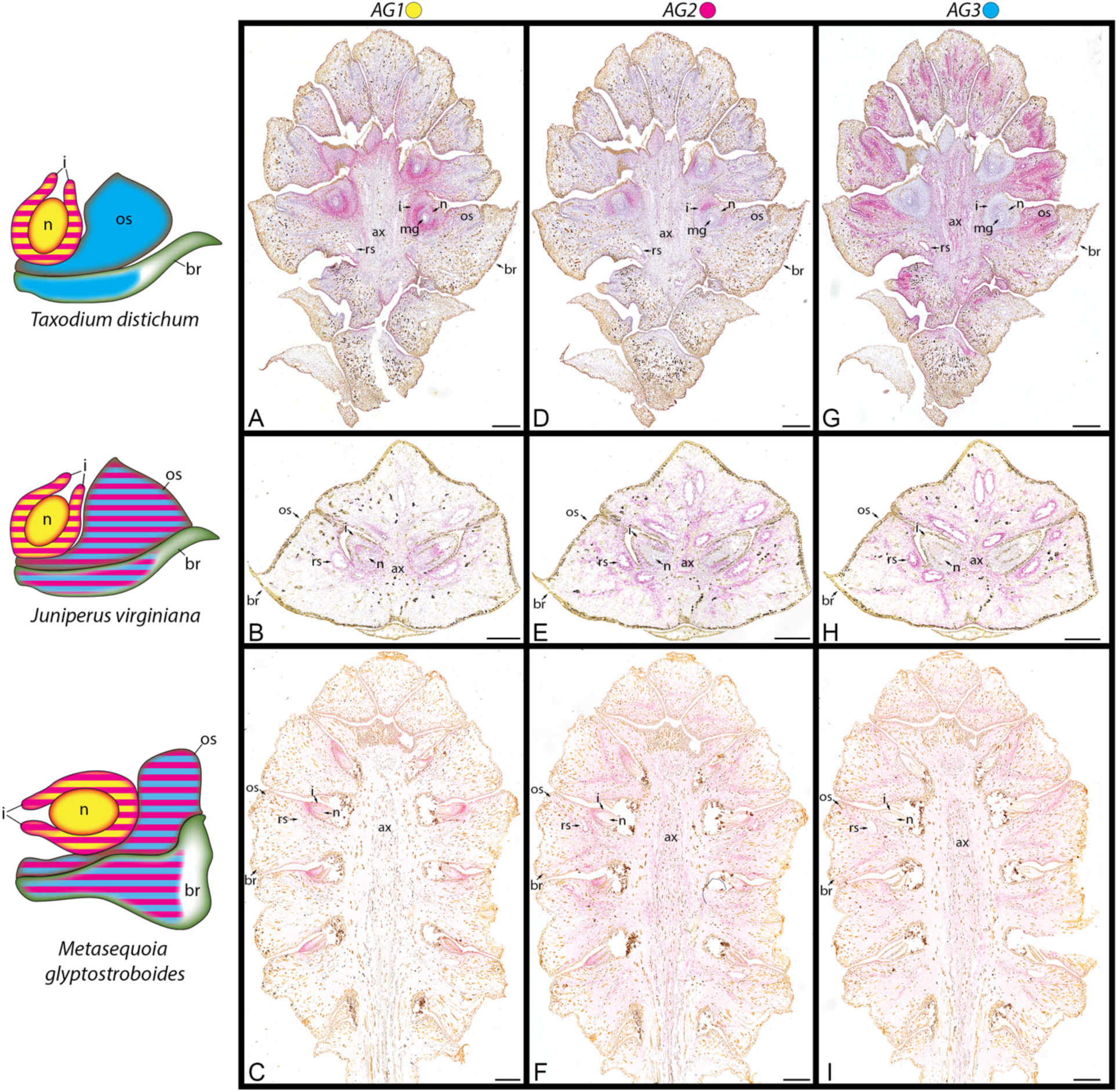
*In situ* hybridization assays of *AGAMOUS* paralogs (**[A-C]** *AG1*, **[D-F]** *AG2*, and **[G-I]** *AG3*) in more mature seed cones of **(A, D, G)** *Taxodium distichum*, **(B, E, H)** *Juniperus virginiana*, and **(C, F, I)** *Metasequoia glyptostroboides*. Magenta stain indicates areas of gene expression. *Taxodium distichum* and *J. virginiana* sections were counterstained with Gill’s Hematoxylin, giving cells a purple color. Note that only one ovule, ovuliferous scale, and bract are labeled for each section due to the repetitive structure of seed cones. Corresponding negative controls are found in SI Appendix, Figs. S5-7. Models at left summarize expression patterns of *AG1* (yellow), *AG2* (fuchsia), and *AG3* (cyan) throughout the ovule, ovuliferous scale, and bract in cones of each species. Note that the ovuliferous scale and bract are completely congenitally fused in *J. virginiana* and *M. glyptostroboides*. ax = cone axis; br = bract; os = ovuliferous scale; n = nucellus; i = integument; rs = resin canal; mg = megagametophyte.

In addition to these overall patterns, we observed several other features of *AG* expression consistent with previous observations in conifers (21, 24, 29). Expression of all three *AG* paralogs was detected to varying degrees in the procambium and epithelial cells of developing resin canals (e.g., Fig. 3G & 3H; SI Appendix, Figs. S8-15, S20-23). In pollen cones, we detected *AG1* expression in the microsporangia (pollen sac) walls and adjacent regions of the microsporophylls in *T. distichum* and *J. virginiana*. However, no comparable *AG2* or *AG3* expression was detected in pollen cones of either of these species (SI Appendix, Figs. S24-31). None of the paralogs, including *AG1*, are expressed within the developing sporocytes or resulting gametophytes (Fig. 3A, D, G; SI Appendix, S8-12, S20-31). Along with the ISH assays on Cupressaceae cones, we also analyzed the expression of the *AG* ortholog in *Picea abies* (Pinaceae), *DAL2*. *DAL2* expression was detected in the ovules and ovuliferous scale, as reported in previous studies (16, 22, 24, 31), but was also present in the epidermis of and diffusely throughout the bract (SI Appendix, Figs. S32, S33), which was previously unreported.

### Cis*-regulatory and protein-level divergence*

Our analyses of intron length and putative transcription factor binding sites (TFBSs) among Cupressaceae *AG* paralogs consistently point toward more *cis*-regulatory differences in *AG2*. In angiosperm *AG* genes, the large first intron (second in some core eudicots) contains TFBSs that are critical for the activation and maintenance of *AG* expression (32–37). Disruptions to these introns or the regulatory sequences within them alter *AG* expression (32–36). With this in mind, to determine if non-conservation of intronic *cis*-regulation is associated with expression divergence among Cupressaceae *AG* paralogs, we explored differences in intron length and the number of TFBSs among *AG* paralogs in eight Cupressaceae species. We found that in Cupressaceae, *AG2* has shorter first introns and longer second introns than those of *AG1* and *AG3* (SI Appendix, Fig. S34). Likewise, *AG2* has fewer TFBSs in the first intron and more in the second intron (SI Appendix, Fig. S35). *AG1* and *AG3*, however, do not differ significantly in the lengths of their first two introns nor the number of TFBS in them (SI Appendix, Figs. S34, S35).

Because positive selection is often indicative of functional divergence at the protein-level, we tested for signatures of positive selection that may have occurred after the *AG* genes duplicated or after the Cupressaceae-Taxaceae divergence. Positive selection was not detected on the branches after each duplication nor on the branches leading to each corresponding paralog for Cupressaceae or Taxaceae. However, 7 sites were identified to be under episodic positive selection along these same branches (SI Appendix, Table. S5).

## Discussion

### *Two duplications occurred in the* AG *subfamily prior to the split between Cupressaceae and Taxaceae*

The *AG* subfamily experienced at least two duplication events before the divergence of Cupressaceae and Taxaceae (Fig. 1). The divergence of these two families is estimated to have occurred ∼180–200 million years ago (2, 28, 38). These are therefore ancient gene duplication events that occurred on a roughly similar timescale to an analogous *AG* duplication in the common ancestor of angiosperms (15). These conifer *AG* paralogs occur sequentially in genomes (SI Appendix, Table S3), indicating these duplications likely occurred via a process of tandem duplication. A fourth *AG* paralog in *Taxus* indicates that a duplication of *AG2* occurred in Taxaceae after the divergence of *Taxus* and *Torreya*.

### AG *expression in Cupressaceae shows a pattern of subfunctionalization*

In the families Pinaceae and Podocarpaceae, which have a single *AG* gene, *AG* is expressed throughout the ovules and ovuliferous scales (16, 17, 22, 24), suggesting that this is the ancestral expression pattern among conifers. In Cupressaceae, the expression of *AG1* in the ovules, *AG2* in almost all tissues but the nucellus, and *AG3* in the ovule-supporting organs suggest that the three paralogs have diverged in their seed cone patterning roles. While *AG2* may have largely retained the ancestral expression pattern, at least in the seed cone, *AG1* and *AG3* exhibit expression patterns consistent with subfunctionalization. *AG1* expression suggests a function in ovule development, and *AG3* expression suggests a role in the development of the ovuliferous scale. The similar expression patterns of these paralogs across the three focal species imply that subfunctionalization occurred before the three species diverged from each other, potentially in the Cupressaceae common ancestor. In *T. distichum*, *AG2* expression appears to have diverged even further, becoming mainly associated with integument development at later developmental stages.

We also observed expression of *AG2* and *AG3* in tissues of the bract, particularly in regions where the bract and ovuliferous scale are congenitally fused, a pattern most evident at early developmental stages when the ovuliferous scale is still primordial (Fig. 2D-I). Previous studies of *AG* orthologs in species of *Picea* (Pinaceae) have not documented expression in the bract (16, 17, 22, 24, 31). In *Picea*, the bract and ovuliferous scale are partially fused at the base and the fusion is most apparent early in development. We detected low levels of *DAL2* (the *AG* ortholog in *P. abies*) in the bracts (SI Appendix, Figs. S32, S33), at least in the region where bract and ovuliferous scale are fused, indicating some expression in the bract may be part of the ancestral expression pattern for *AG* in conifers. However, given that *AG* expression corresponds to regions of fusion between bract and ovuliferous scale in Cupressaceae and occurs at low levels in a similar region in Pinaceae, it is possible that the strong expression in Cupressaceae bract tissue reflects the high degree of synorganization of the bract and ovuliferous scale in Cupressaceae, as well as neofunctionalization for bract development. It may also reflect a “fading borders” mode of gene expression, in which organ identity genes are expressed broadly but weakly at the outer limits of their expression zone, as has been shown in early-diverging angiosperms (39, 40).

In seed plants, *AG* is also involved in the development of pollen-producing organs (microsporangia and microsporophylls) (12, 17, 19–21, 23–26), but not all Cupressaceae *AG* paralogs appear to be involved in pollen cone development. Our ISH assays of *T. distichum* and *J. virginiana* pollen cones show that while *AG1* is expressed in microsporangia and microsporophylls, *AG2* and *AG3* are not expressed at all, at least at the developmental stage sampled (SI Appendix, Figs. S24-31). Consistent with our ISH results, a transcriptome analysis in *Cunninghamia lanceolata* found that while *AG1* is expressed at high levels in pollen cones, *AG2* and *AG3* are expressed only at low levels (29). Together this indicates that *AG1* is likely associated with sporangium development more generally, whereas *AG2* and *AG3* appear to be involved exclusively in seed cone development. More specifically, *AG2* and *AG3* appear to be involved in the development of the “vegetative” components of the seed cone: the ovuliferous scale and bract in the case of *AG3* and all structures other than the nucellus for *AG2*, including the ovule integuments. *AG1* is notably the only paralog expressed within the nucellus, which is the megasporangium and a site of meiosis in the life cycle.

While not the explicit focus of this paper, this study also reveals interesting patterns of *AG* expression in Taxaceae reproductive structures. Two different studies have investigated *AG* expression spatially in *Taxus* (22, 23). Our gene tree indicates these studies assayed different paralogs: *TgAG* from *T. globosa* (*AG1*) and *AG* from *T. baccata* (*AG3*) (Fig. 1). Like Cupressaceae, *AG1* was found to be expressed in the nucellus and aril (a possible ovuliferous scale homolog) of *T. globosa*. However, *AG1* expression is absent in the integument at later stages of cone development (22), unlike that of Cupressaceae *AG1*. In *T. baccata*, *AG3* is expressed strongly in the ovule as well as in the aril (23), which differs from Cupressaceae *AG3* where there is little to no expression in the ovules. These expression patterns suggest that each paralog has different functions in Cupressaceae and Taxaceae and that these functional divergences occurred independently in the two families. Further research in Taxaceae is needed to understand *AG* expression through development and the role of *AG* paralogs in the reproductive biology of Taxaceae.

### *Evidence of functional divergence on both a* cis*-regulatory level and a protein level*

Studies in angiosperms have identified *cis*-regulatory elements in the first and second introns (or second and third in some core eudicots) of members of the *AG* subfamily. Disruptions to these elements, particularly in the first intron, have led to changes in *AG* expression, including loss of and shifts in timing of expression (32–37). The truncation of the first intron and expansion of the second intron (and the resulting decrease and increase of the number of TFBSs) for Cupressaceae *AG2* may underpin its seed-cone-specific expression. Notably, the second intron of *AG1* and *AG3* are short and hold few putative TFBSs; thus, *AG2* seems to have shifted much of its *cis*-regulation to its second intron from its first. We could not identify a potential mechanism by which *AG1* and *AG3* expression differ. *AG3* first introns are longer and contain more TFBSs than *AG1* first introns, on average, but these differences were not statistically significant (SI Appendix, Figs. S34, S35). Other mechanisms that we did not explicitly study, such as changes to promoters, the sequences of the TFBSs, or the distance between TFBSs, may underpin the difference in expression among the paralogs and should be examined in future studies. Interestingly, the first intron for *AG2* in *Taxodium* and *Cryptomeria*, both members of the subfamily Taxodioideae, is even shorter than the first *AG2* intron of other genera (except *Thuja*) (SI Appendix, Dataset S1). This reduction in intron size may explain why *T. distichum AG2* expression differs from that of *M. glyptostroboides* and *J. virginiana* (Fig. 3D-F).

Although we did not detect positive selection on the branches post-duplication or the branches leading to each Cupressaceae and Taxaceae paralog, seven sites (i.e., codons) were found to be under episodic positive selection along these branches. Five of these sites are in the K domain, which is responsible for protein-protein interactions (41). This suggests that in addition to divergence in expression patterns among paralogs, there may also be divergence in protein function, particularly in protein-protein interactions. This is further exemplified by the observation that in some Cupressaceae species, a portion of the K3 subdomain has been duplicated in their *AG2* genes (SI Appendix, Fig. S36). This subdomain encodes an α-helix critical for the formation of tetramers with other MADS proteins to modulate gene expression (41, 42). Similar instances of positive selection in the K3 subdomain of *AG* paralogs has been documented in the monocot order the Zingiberales (43). In snapdragon, an amino acid insertion in the K3 subdomain of its *AG* ortholog, *FARINELLI* (*FAR*), limits its ability to interact with certain MADS proteins (some of the E-class genes), resulting in *FAR*’s stamen-specific function (44). Overall, molecular evolution in the K domain points to functional changes among conifer *AG* paralogs driven by shifts in protein-protein interactions.

### *Subfunctionalization of* AG *may have decoupled ovule and ovuliferous scale development*

The seed cones of Cupressaceae are morphologically diverse and differ from those of most other conifers in the relative timing of ovule and ovuliferous scale development, as well as the position and orientation of ovules throughout development (45). Our results provide a mechanism to explain these unique traits of Cupressaceae seed cones. Gene duplication can reduce pleiotropy and allow for the evolution of different morphologies without constraints (5). In this vein, duplication and subsequent subfunctionalization of *AG* genes for ovule and ovuliferous scale development may have allowed these two organs to become developmentally decoupled, increasing the developmental modularity of the seed cone (5, 46) and allowing for shifts in the developmental timing (heterochrony) and position (heterotopy) of the ovules relative to the ovuliferous scale. Heterochrony and heterotopy have been previously invoked to explain the ontogenetic diversity of Cupressaceae seed cones (21, 27), but an underlying mechanism has been lacking. Interestingly, a study that deleted a TFBS for LEAFY, an *AG* activator, in the C-function gene of snapdragon, *PLENA* (*PLE*), found that this delayed *PLE* expression until fourth-whorl development such that stamens transformed into petals, but carpels remained unaffected (34). Similar changes to the *cis*-regulatory elements in the introns of the *AG* paralogs in Cupressaceae may have facilitated changes in the timing of the development of the ovuliferous scale or the ovule.

These developmental changes have important functional consequences for reproductive biology. The ovuliferous scale has multiple roles in conifer reproductive systems, and these roles change throughout cone ontogeny (47). At pollination stage the ovuliferous scale is responsible for orienting the ovule’s micropyle (opening of the integument) toward the cone axis, which is ancestral in conifers and necessary for some modes of pollination (38, 45). The ovuliferous scale may also help to direct wind-borne pollen towards the micropyle (48–50). At later developmental stages, the ovuliferous scale is involved in protecting developing seeds and, in some cases, forms wings for wind dispersal (Pinaceae) or fleshy tissues that attract vertebrate dispersers (Podocarpaceae). Notably, all members of Cupressaceae have cones with ovules that are erect (micropyle facing away from the cone axis) and exposed at pollination, and the cones become heavily armored or even fleshy at maturity (38, 45). Decoupling ovule and ovuliferous scale development would enable pollination systems that involve erect ovules by delaying or preventing ovule inversion by the ovuliferous scale, while simultaneously retaining the later roles of the ovuliferous scale in seed protection and dispersal. In this way, duplication and functional divergence of *AG* genes, which are key developmental regulators of seed plant reproductive structures, may have allowed for the evolution of new pollination systems and seed cone phenotypes.

### *Parallel evolution of* AG *in conifers and angiosperms*

Angiosperms are the only other group in which major duplications in the *AG* subfamily are known to have occurred (Fig. 4; SI Appendix, Fig. S1) (15, 18, 19, 22, 24, 25, 51, 52). Flowering plants share an ancestral duplication of *AG*, which gave rise to the C and D lineages (15). Additional duplications nested within angiosperms have occurred as well, such as in the C lineage of core eudicots, forming the *euAG* and *PLE* lineages (15), in the Ranunculales (53), and in several monocot lineages (43, 54). The patterns of divergent expression observed among the Cupressaceae paralogs indicate similar patterns of gene duplication and subfunctionalization have occurred independently in angiosperms and conifers.

**Fig. 4.**
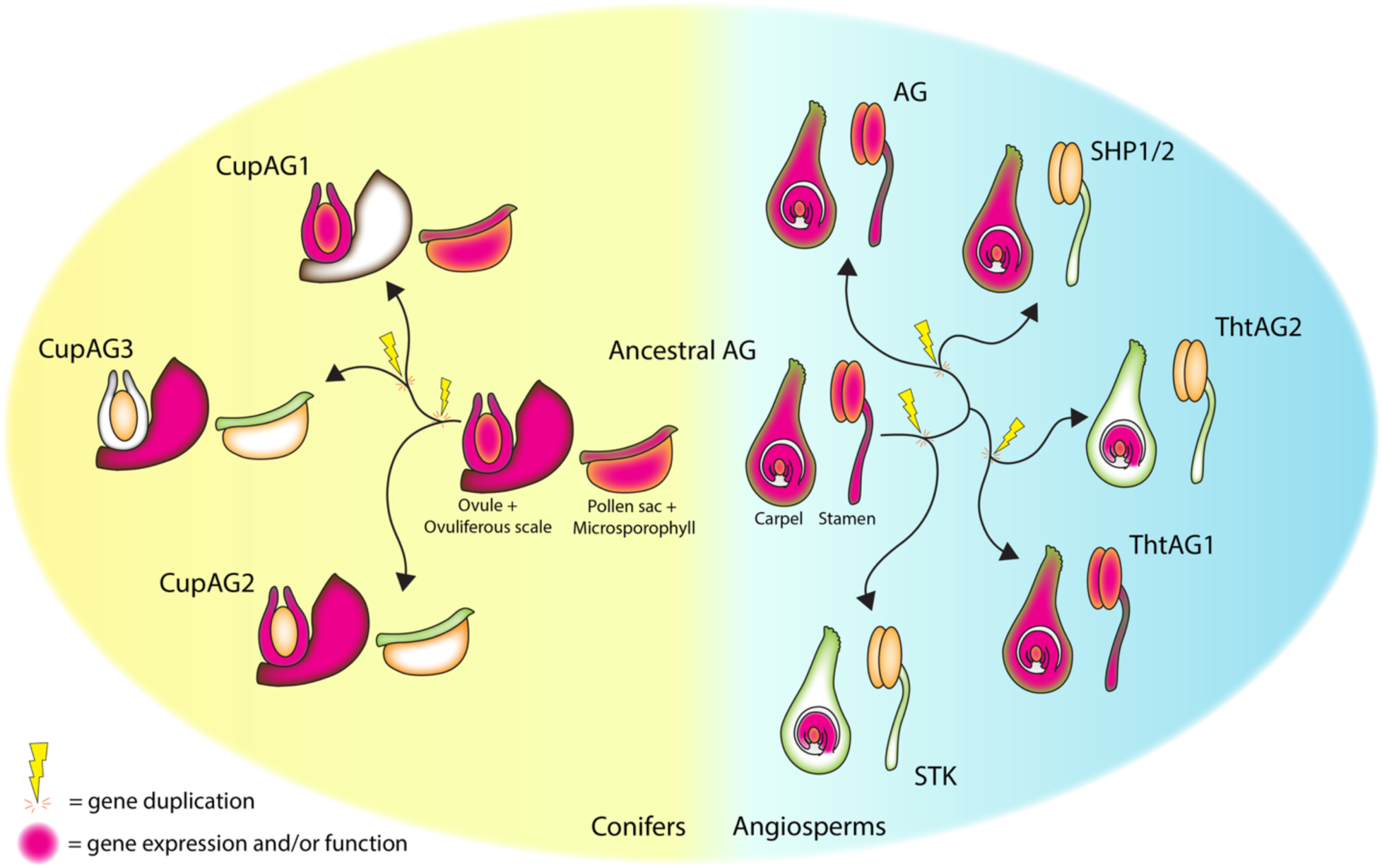
Parallel trajectories in the evolution and expression of *AG* in angiosperms and conifers. Hypothesized ancestral expression patterns for seed and pollen-producing organs are shown at center. In angiosperms, *AG* functions in the patterning of carpels, ovules, and stamens (pollen organ) and is expressed throughout these structures during development. Comparable expression patterns and functions have been hypothesized for conifers, with *AG* expressed in ovules, ovuliferous scales, pollen sacs (microsporangia), and microsporophylls. Note that the bract is not shown in these diagrams. Gene duplications in each group are indicated by arrows and lightning bolts. Within angiosperms, key duplications leading to subfunctionalization of *AG* are shown (15, 53, 55, 56). D-lineage genes such as *STK* in *Arabidopsis* are specialized in ovule development. An independent duplication in *Thalictrum* (Ranunculales) resulted in the ovule-specific *ThtAG2.* Within core eudicots, duplications leading to *Arabidopsis SHP1* and *SHP2* resulted in carpel- and ovule-specific functions. Note the loss of expression in stamens for *STK*, *ThtAG2*, and *SHP1/2*. Within conifers, Cupressaceae *AG2* (*CupAG2*) is expressed in the ovule integument and ovuliferous scale and absent in pollen organs, similar to *SHP1/2*. In seed cones, *CupAG1* is expressed exclusively in the ovules, similar to *STK* and *ThtAG2*, but has retained expression in pollen organs. *CupAG3* is exclusively expressed in the ovuliferous scale and absent in pollen organs. Note that *CupAG1* is the only paralog expressed in the nucellus (megasporangium) of the ovules and in the pollen organs (including the microsporangia), indicating a broader role in sporangium development.

Subfunctionalization to roles in ovule development or ovule-specific expression is a consistent pattern seen in several angiosperm *AG* paralogs, such as the D lineage, the *PLE* genes in *Arabidopsis* (*SHATTERPROOF1* [*SHP1*] and *SHP2*), and *ThtAG2* of the ranunculid genus *Thalictrum* (Fig. 4) (15, 53, 55, 56). The occurrence of similar patterns of functional divergence of an *AG* paralog to ovule expression, observed in Cupressaceae *AG1*, suggests that this is a common pathway for *AG* genes after duplication. Ovule-specific expression in angiosperms often involves a loss of expression in stamens (21, 29), but Cupressaceae *AG1* genes are also expressed in pollen cones (SI Appendix, Figs. S24, S28) (21, 29), indicating a broader role in sporangium development that results in ovule-specific expression in seed cones. Unlike the carpels and stamens of bisexual angiosperm flowers, which develop in synchrony from the same floral meristems, the development of pollen and seed cones of Cupressaceae are spatially and temporally segregated. As a result, the sporangium-specific *AG1* may not be under the same selective pressures that resulted in loss of expression in the stamens of their ovule-specific angiosperm analogs.

Loss of expression in pollen-producing organs was instead observed for *AG2* and *AG3*. The expression of Cupressaceae *AG2* throughout the ovule-supporting organs (ovuliferous scale and fused parts of the bract) and integument and its loss in pollen cones is similar to *SHP1* and *SHP2* in *Arabidopsis*, which are involved in carpel and ovule development but not stamen development (Fig. 4) (56). Like *AG2* in Cupressaceae, *SHP1* and *SHP2* lack elements that contain various TFBSs and have shorter second introns, which are positionally homologous to the first intron in conifers (33). It is therefore possible that the similar patterns of Cupressaceae *AG2* expression and *Arabidopsis SHP1* and *SHP2* expression arose independently through similar mechanisms. In contrast, *AG3* appears to be exclusively involved in the development of the ovule-supporting organs in Cupressaceae but not the ovules. The equivalent in angiosperms would be expression in the carpel but not the ovule, but to our knowledge this has not been observed. While it is possible this occurs but has not been documented, it may also reflect differing functional constraints of ovuliferous scales and carpels. Unlike ovuliferous scales, the carpel encloses the ovules throughout their development and so the two structures must develop in synchrony to some extent.

The duplication of *AG* in angiosperms to form the C and D lineages and their subsequent subfunctionalization has been proposed as a mechanism that has facilitated evolutionary modifications of both carpels and ovules (15). Our data on the evolution and expression of conifer *AG* genes suggests a similar process has occurred in conifers, potentially on a similar time scale given that angiosperms and Cupressaceae are estimated to be roughly the same age (28, 57). Angiosperms and conifers therefore appear to have undertaken parallel evolutionary trajectories in the diversification of the *AG* subfamily, which may have shaped their reproductive biology through time and led to the evolution of phenotypic novelties.

## Materials and Methods

### Plant material

Developing seed cones were collected from individuals of *Metasequoia glyptostroboides*, *Taxodium distichum*, and *Juniperus virginiana* (Cupressaceae; subfamilies Sequoioideae, Taxodioideae, and Cupressoideae, respectively) and *Picea abies* (Pinaceae) on the University of Kansas (KU) campus from March-May in 2023-2025. Cones were collected near pollination stage, as well as after pollination, when the ovuliferous scale had grown to enclose the ovules. An additional stage was collected for *M. glyptostroboides* in early November of 2025 during ovule development. Pollen cones of *T. distichum* and *J. virginiana* were collected in August and September of 2025 and 2026; the individuals of *M. glyptostroboides* from which we collected seed cones did not produce pollen cones. Vouchers for each species have been deposited in the KU Ronald L. McGregor Herbarium (KANU) under catalog numbers 442980-442985, 447090, 447094.

### *Identification of* AG *genes*

To identify the sequence of *AG* genes in our study species and obtain transcriptomic resources for *in situ* hybridization probe design (see In situ *mRNA hybridization [ISH]* below), we sequenced, assembled, and functionally annotated transcriptomes from the seed cones of *T. distichum* and *J. virginiana* (see SI Appendix, Supporting Text and Table S2 for more information). *AG* gene sequences from *Picea abies* (Pinaceae), *Podocarpus matudae* (Podocarpaceae), *Cunninghamia lanceolata* (Cupressaceae), *Taxus globosa*, and *T. baccata* (Taxaceae) (22, 23, 58) were used as queries to search for *AG* subfamily members in the genomes and transcriptomes of conifers and other gymnosperms (SI Appendix, Table S1) using tBLASTX (59). For species without genome annotations for their respective *AG* genes, Exonerate (v. 2.4.0) (60) was used to predict the coding sequence of identified *AG* subfamily members using translations of the query sequences listed above, and we corrected predictions by eye using alignments with MAFFT (61) in AliView (62).

### Gene tree reconstruction

To confirm the membership of identified genes to the *AG* subfamily and infer the position of duplications in conifer phylogeny, we conducted a phylogenetic analysis of MADS-box genes sampled from all major seed plant groups (angiosperms, Gnetophytes, conifers, *Ginkgo*, and cycads), which included the putative conifer *AG* genes identified above. The *Arabidopsis* Type I MADS-box genes *DIA*, *AGL23*, and *AGL62* served as the outgroup since the *AG* subfamily are Type II MADS-box genes (SI Appendix, Fig. S1, Table S3). We aligned coding sequences as translated amino acids in AliView (v. 1.28) (62) with MAFFT (v. 7.526) (61) under default parameters. We removed sites consisting of >75% gaps with trimAl (v. 1.5.0) (63) as well as the less conserved I and C domains. This resulted in a matrix of 362 sequences and 456 characters. 100 tree searches were conducted with RAxML (v. 8.2.12) (64) under the GTR + Γ model as identified by ModelFinder (65) in IQ-TREE (v. 2.3.6) (66) according to corrected Akaike information criterion (AICc) with 500 nonparametric bootstrap replicates to assess support.

### In situ *mRNA hybridization (ISH)*

To analyze spatial expression of *AG* genes within seed cones, we performed in situ hybridization (ISH) assays at each developmental stage collected for the focal species using RNAscope from Advanced Cell Diagnostics (ACD) (67). Probes targeting the mRNA sequence of *DAL2* in *P. abies* and the coding sequences for all *AG* paralogs identified in *M. glyptostroboides*, *T. distichum*, and *J. virginiana* were designed by the ACD Probe Design Team (see SI Appendix, Table S4 for more information on probes). The genome of *M. glyptostroboides* (68) and transcriptomes of *P. abies* (69, 70), *T. distichum*, and *J. virginiana* were used to assess potential cross-reaction with non-target transcripts. Tissues were processed, sectioned, and assayed for gene expression by ISH as described in Anaya et al. (71), with a 12-16 hour fixation time, 10-15 minute protease treatment, and no Hematoxylin counterstain for most assays. The full protocol from Anaya et al. (71) can also be found on protocols.io (72). Images were taken using a Nikon Eclipse LV100ND upright brightfield microscope with a DS-Fi3 camera, using Manual Large Image capture in NIS-Elements D software (v. 5.20.02). Images were minimally processed for brightness and contrast in Adobe Photoshop 2026.

### *Identification of differences in* cis*-regulation among paralogs*

Known regulatory sequences for angiosperm *AG* genes are located in the large first intron (second in some species) (32–37). This intron contains transcription factor binding sites (TFBSs) for LEAFY (LFY); WUSCHEL (WUS); other MADS-box proteins; CCAAT-box binding proteins; PERIANTHIA (PAN), a member of the bZIP family; and ULT1 INTERACTING FACTOR (UIF1, also known as HHO5), a GARP-domain protein (32–37). In particular, the CCAAT-box binding proteins bind to two CCAATCA motifs flanking ∼37 bp (henceforth, the CCAATCA box), and the TFBSs for PAN and UIF1 are located in the 19-bp aAGAAT box with UIF1 acting as a possible cofactor for PAN (33, 35, 37). These TFBSs are crucial for the activation and maintenance of *AG* expression in angiosperms. Mutations to these sequences or the introns that carry them cause changes in *AG* expression, and some (the aAGAAT and CCAATCA boxes) are notably absent in some angiosperm *AG* paralogs (32–36). Outside of the first intron, some TFBSs for LFY, WUS, and MADS-box proteins have also been found in the second intron of some angiosperms (36). Thus, to identify differences in *cis*-regulation that may underpin the divergence in expression among Cupressaceae *AG* paralogs, we compared the length of the first two introns among each *AG* paralog in Cupressaceae and scanned the two introns for sequence motifs of the TFBSs with the FIMO (73) program in the MEME suite (74). Ideally, we would identify these regulatory sequences by aligning the introns of the conifer *AG* paralogs and angiosperm *AG* genes. However, this was not possible due to the length and poor sequence conservation of the conifer introns. Previous studies in angiosperms also have had difficulty with intron alignment among paralogs and have relied on predictive software for identification (33, 36). Thus, all identified regulatory sequences here are putative in the sense that they were identified using predictive software rather than multiple sequence alignments or experimental approaches.

Eight species with assembled genomes were included in the analysis (SI Appendix, Table S1) (68, 75–80). Intron sequences and their lengths were based on Exonerate outputs or the associated genome annotation file. The binding motifs for LFY (matrix ID: MA0590.1) (81) and UIF1 (matrix ID: MA1164.2) (82) were downloaded from the JASPAR database (83). Motifs for the other TFBSs were pulled from previous publications (32, 33, 35, 36) and converted into MEME-formatted motifs. Default parameters were used, apart from WUS and PAN, for which we used a more lenient p-value threshold (0.0005 instead of 0.0001) since the default threshold was too strict and yielded few or no hits. We used Welch’s ANOVA on log10-transformed intron lengths and Kruskal-Wallis tests on the number of identified TFBSs to assess differences among each Cupressaceae paralog for each intron. Post-hoc pairwise comparisons between each paralog were performed by Games-Howell and Dunn’s tests. Statistical analyses were performed using the rstatix package (v. 1.1.0) (84) in R (v. 4.6.0) (85).

### Test for selection

To test for potential positive selection (ω [dN/dS] ratios > 1) on paralogs, we used the aBSREL (86) and MEME (87) programs in HyPhy (v. 2.5.96) (88, 89). To generate the input alignment, we subsampled our matrix to include only *AG* genes from conifers and realigned and trimmed the matrix as specified above for the gene tree reconstruction, but without removing the I and C domains. For the input tree, we pruned the MADS-box gene tree with the ape package (v. 5.8.1) (90) in R (85) to include only conifer *AG* genes. Because functional divergence can happen after duplication and Cupressaceae and Taxaceae *AG* genes appear to have functionally diverged based on the ISH results from this study and previous studies (22, 23), the branches after each duplication and the branches of *AG1*, *AG2*, and *AG3* for Cupressaceae and Taxaceae were specified as foreground branches (Fig. 1).

## Data, Materials, and Software Availability

Raw sequencing reads and transcriptome assemblies generated for this study will be deposited on NCBI under BioProject PRJNA1556916. Other data generated from this study, including multi-sequence alignments, tree files, unedited images, and associated files for the transcription factor binding site and selection analyses, will be deposited on Zenodo.

## Acknowledgements

The authors thank Dr. Brian Sanderson for bioinformatics support, as well as Drs. Mark Holder, Paulyn Cartwright, Jae Choi, Josie Chandler, and the KU EEB Genetics community for feedback on this project. We thank Dr. Jesús Martínez-Gómez for his suggestions to look at transcription factor binding sites and Dr. Elena Kramer for sharing unpublished data that led us to look more closely at pollen cones. We thank Haylee Coffman, Frances Szaraz, Aspen Avelar, Cyrus Shaikh, Dr. Ana Andruchow-Colombo, and Dr. Kelly Pfeiler for assistance with collection, tissue processing, and histology. This work was performed at the HPC facilities operated by the Center for Research Computing at the University of Kansas supported in part through the National Science Foundation MRI Award #2117449. Research reported in this publication was made possible in part by the services of the KU Genome Sequencing Core, which is supported by the National Institute of General Medical Sciences (NIGMS) of the National Institutes of Health under award number P30GM145499, and K-INBRE, which is supported by the IDeA Program of the National Institute of General Medical Science award number P20 GM103418, as well as the University of Kansas Center for Genomics. A. T. N. is supported in part by the National Science Foundation Graduate Research Fellowship Program (Award No. 1940699). This research was funded by the following awards and institutions: KU Biodiversity Institute Leaman Harris and Panorama research awards to A.T.N.; Botanical Society of America Graduate Student Research Award to A.T.N.; Society for the Study of Evolution Lewontin Early Award to A.T.N.; National Science Foundation CAREER Award (DEB 2338756) to K.K.S.M..

## Author Contributions

K.K.S.M. conceptualized the research; A.T.N, L.C.H., and K.K.S.M. designed research; A.T.N. gathered and analyzed the data; A.T.N, L.C.H., and K.K.S.M. interpreted the data; A.T.N. and K.K.S.M. wrote the paper and created the figures; A.T.N, L.C.H., and K.K.S.M. revised the paper.

