## Supplementary material for "Parallel Trajectories in the Evolution of C-class MADS-box Genes in Flowering Plants and Conifers": SI Appendix

### Supporting Information

#### Supporting Text

#### Transcriptome sequencing and assembly for *Taxodium distichum* and *Juniperus virginiana*

Post-pollination stage seed cones of *Taxodium distichum* and *Juniperus virginiana* were collected for RNA sequencing from individuals located on KU campus. Seed cones were flash frozen in liquid nitrogen and stored at -80ºC. Each replicate of *T. distichum* contained one seed cone, and each replicate of *J. virginiana* pooled 5 seed cones. RNA extractions were performed using the Spectrum Plant Total RNA Kit (Sigma-Aldrich) per manufacturer's protocol with on-column DNase digestion, and RNA concentration and purity were checked with a NanoDrop. Samples were then given to the KU Genome Sequencing Core for RNA quality check, library construction, and sequencing. RNA concentrations and integrity were analyzed using a Qubit and an Agilent TapeStation 4150, respectively. All samples had a RIN above 8. The NEBNext Ultra II Directional RNA Library Prep Kit for Illumina with the NEBNext Poly(A) mRNA Magnetic Isolation Module was used to prepare cDNA libraries with Unique Dual Indices for each sample. Two rounds of AMPure Bead size selection and one round of BluePippin Gel size selection were used to remove off-target peaks (primer and adapter dimers). 150 bp paired-end sequencing was done on one lane of Illumina's NextSeq2000. This resulted in an average of approximately 20.9 million read pairs for *T. distichum* samples and 17.9 million for *J. virginiana* samples.

We used fastp (v. 0.23.4) (1) to remove reads less than 85 nucleotides long, reads that had more than 20% of their bases with Phred scores less than 30, and adapter sequences. FastQC (v. 0.11.8) (2) was used to check for proper read trimming. *De novo* transcriptome assembly for *T. distichum* and *J. virginiana* was performed with Trinity (v. 2.15.1) (3), specifying reverse-forward strand-specific RNA-seq read orientation and all other parameters default. We ensured proper transcriptome assembly by using E90N50 values, read representation to ensure >80% of trimmed reads mapped back to the transcriptomes with Bowtie 2 (4) built into the Trinity toolkit (5), and the BUSCO (v. 5.4.7) (6, 7) gymnosperm dataset (8) (see Table S2 for assembly statistics). To calculate the E90N50 statistics, RSEM (9) in the Trinity toolkit (5) was used for quantification. Open reading frames were predicted using TransDecoder (v. 5.7.1) (5), and functional annotation was done with the Trinotate annotation suite (v 4.0.2) (10). For the transcriptome of *Picea abies*, previously published sequencing reads (11, 12) were downloaded from NCBI's Sequencing Read Archive (ERR10020120-ERR10020129; ERR2632798-ERR2632807; ERR6099827-ERR6099846), and assembly was done as described prior except specifying a 70-nucleotide length requirement and 15% unqualified percent limit for fastp.

### Figures

A

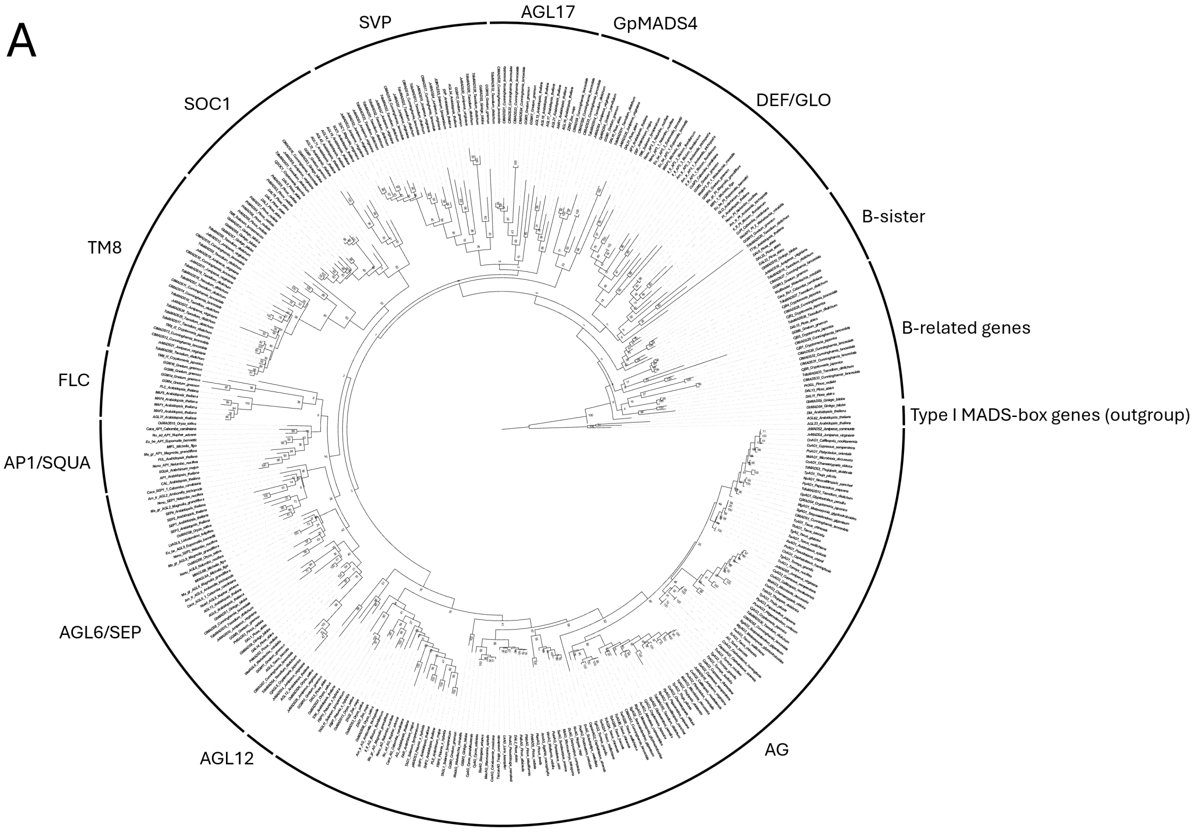
SVP
AGL17
GpMADS4
DEF/GLO
SOC1
B-sister
TM8
B-related genes
FLC
Type I MADS-box genes (outgroup)
AP1/SQUA
AGL6/SEP
AGL12
AG

B

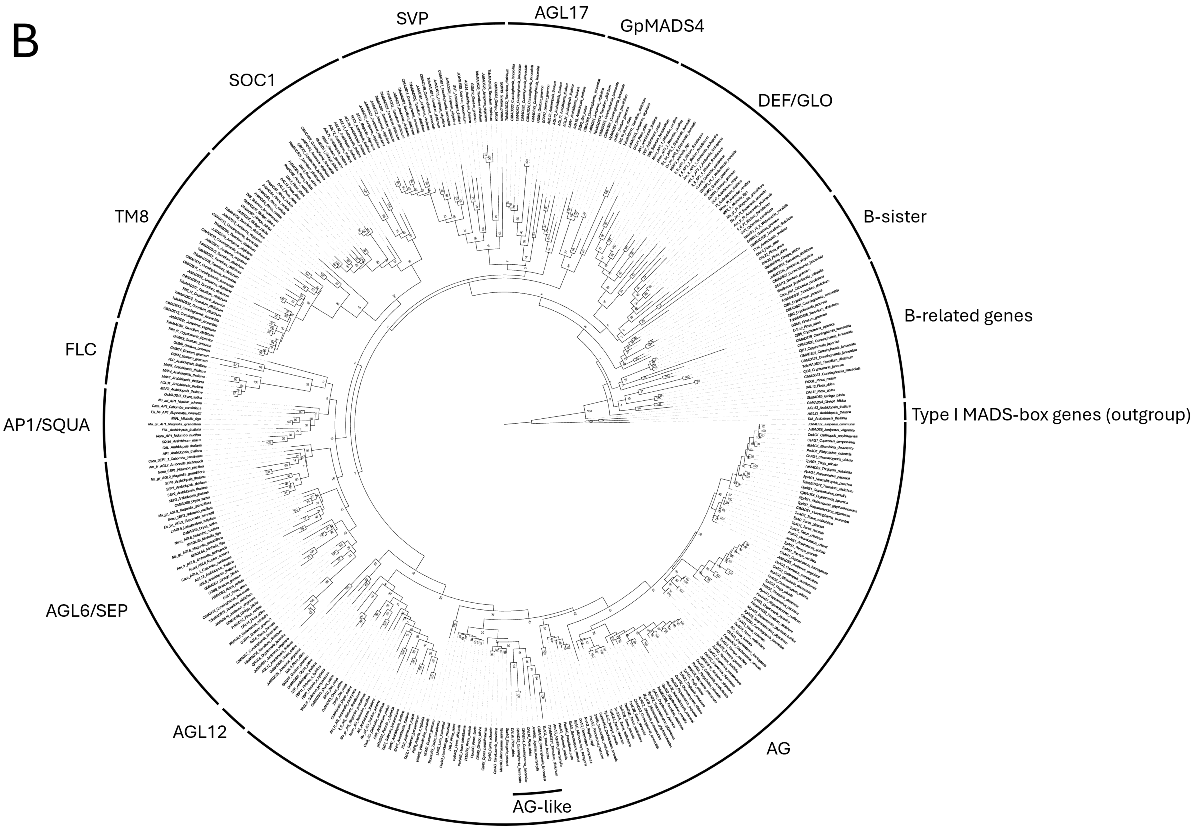
SVP
AGL17
GpMADS4
SOC1
DEF/GLO
TM8
B-sister
B-related genes
FLC
Type I MADS-box genes (outgroup)
AP1/SQUA
AGL6/SEP
AGL12
AG
AG-like

**Fig. S1.** MADS-box gene trees **(A)** without and **(B)** with long-branch *AG*-like clade. Fig. 1 in main text is pruned from (A). Relationships among the major Type II MADS-box genes are largely consistent with previous studies (13, 14) except for the B-class (DEF/GLO), B-sister, and B-related genes, which do not get resolved as monophyletic. In addition to *AG* duplications within the Cupressales, we also found evidence for an additional *AG*-like clade in conifers, shown in (B). This clade of *AG*-like genes was placed sister to cycad *AG* with low bootstrap support and included sequences from members of Pinaceae, Araucariaceae, Taxaceae, and Cupressaceae. Although the phylogenetic placement indicates that members of this clade are *AG* genes, we believe they are not for the following reasons. The clade exists on a long branch and its relationship to the other *AG* genes have low bootstrap support, suggesting an unstable relationship to the other *AG* genes. When these genes are removed from the analysis, shown in (A), bootstrap values within and for the overall *AG* clade increase, further exemplifying this group's unstable relationship and indicating that this group occasionally gets reconstructed not as members of the *AG* clade. Despite having an inferred closer relationship to Pinaceae *AG* than the other conifer *AG* genes in this study, a BLAST search using Pinaceae *AG* as queries finds the other conifer *AG* genes first rather than members of this *AG*-like clade. Some of these *AG*-like genes lack motifs in the C domain characteristic to that of the *AG* subfamily across seed plants (15, 16). For many of the *AG*-like genes isolated from reproductive tissues (17, 18), the sequences are fragmented or incomplete, suggesting low expression levels in reproductive tissues and divergence from the canonical *AG* expression pattern and function. Thus, further investigation into the relationships, expression patterns, and functions of this clade is needed.

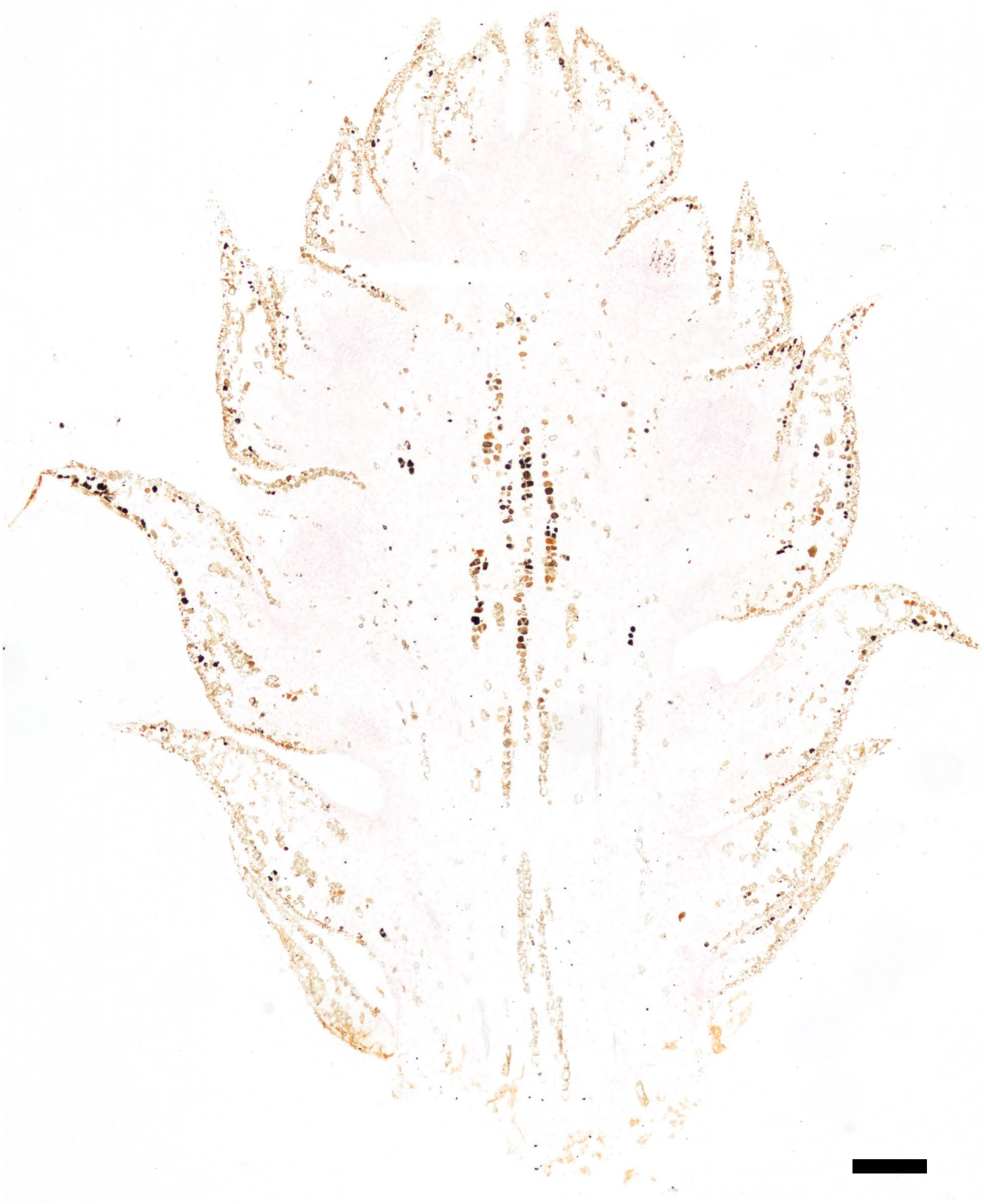

**Fig. S2.** Negative control for *Taxodium distichum* images from Fig. 2 in main text. Scale bar = 250 µm.

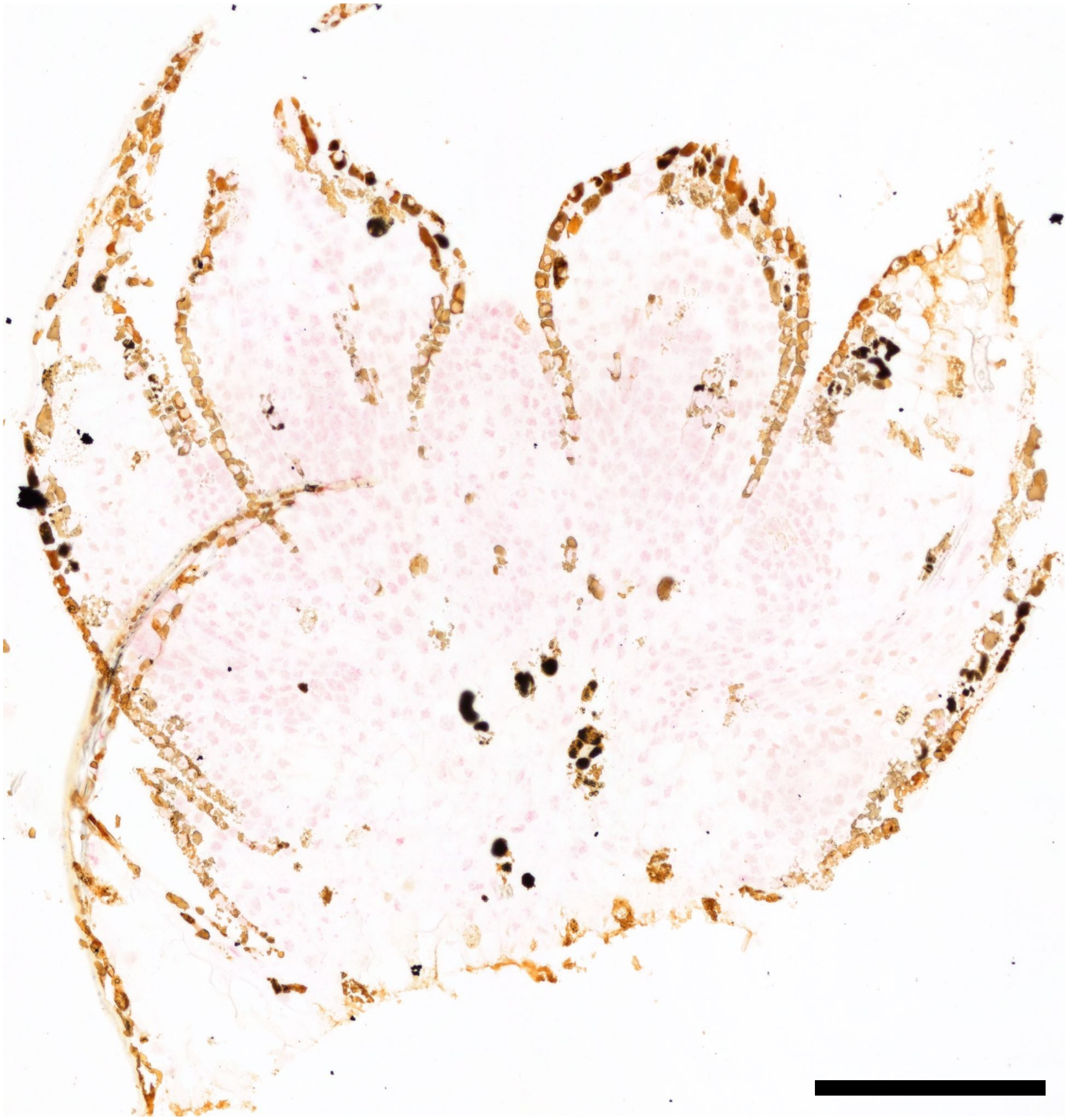

**Fig. S3.** Negative control for *Juniperus virginiana* images from Fig. 2 in main text. Scale bar = 250 μm.

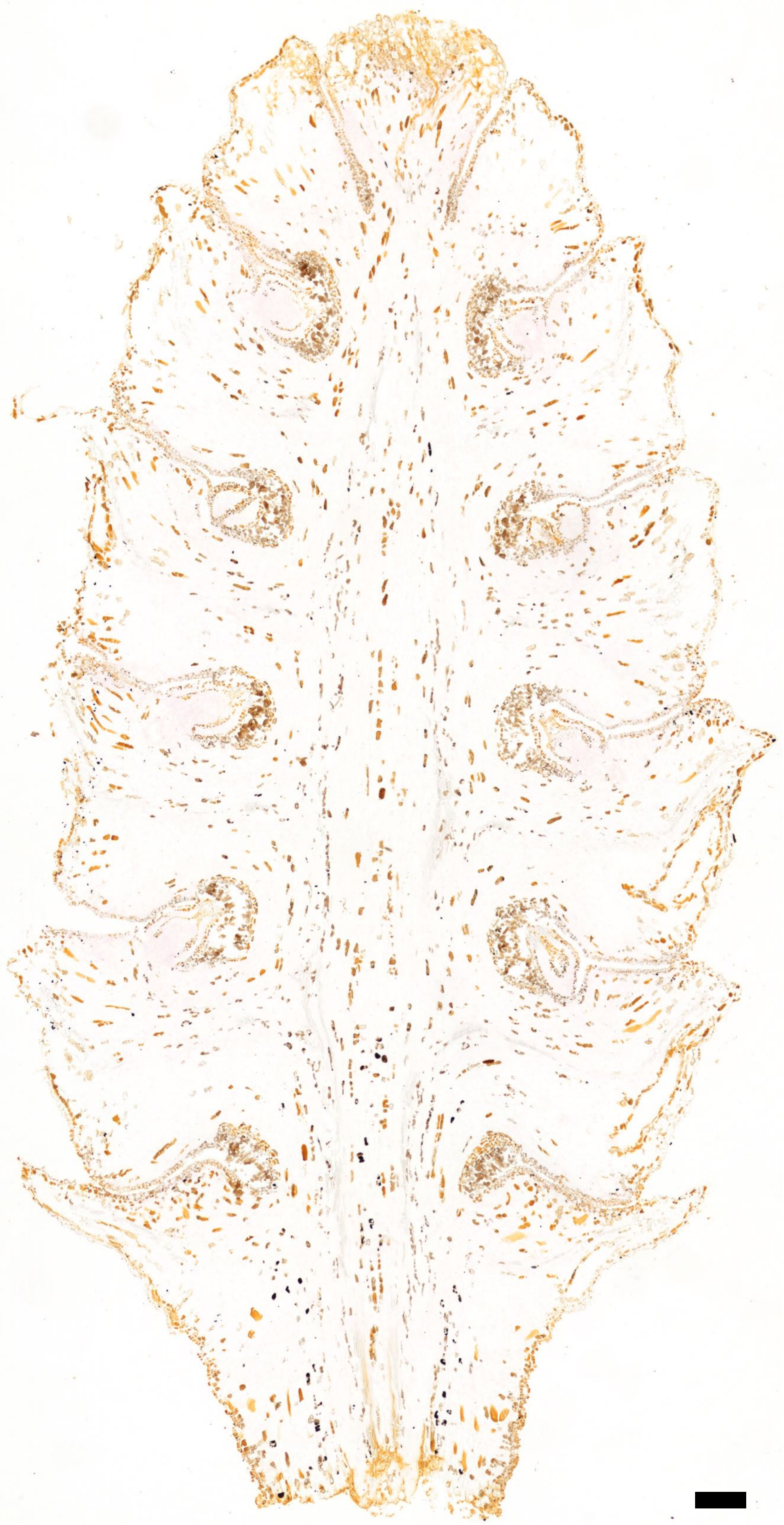

**Fig. S4.** Negative control for *Metasequoia glyptostroboides* images from Fig. 2 in main text. Scale bar = 250 µm.

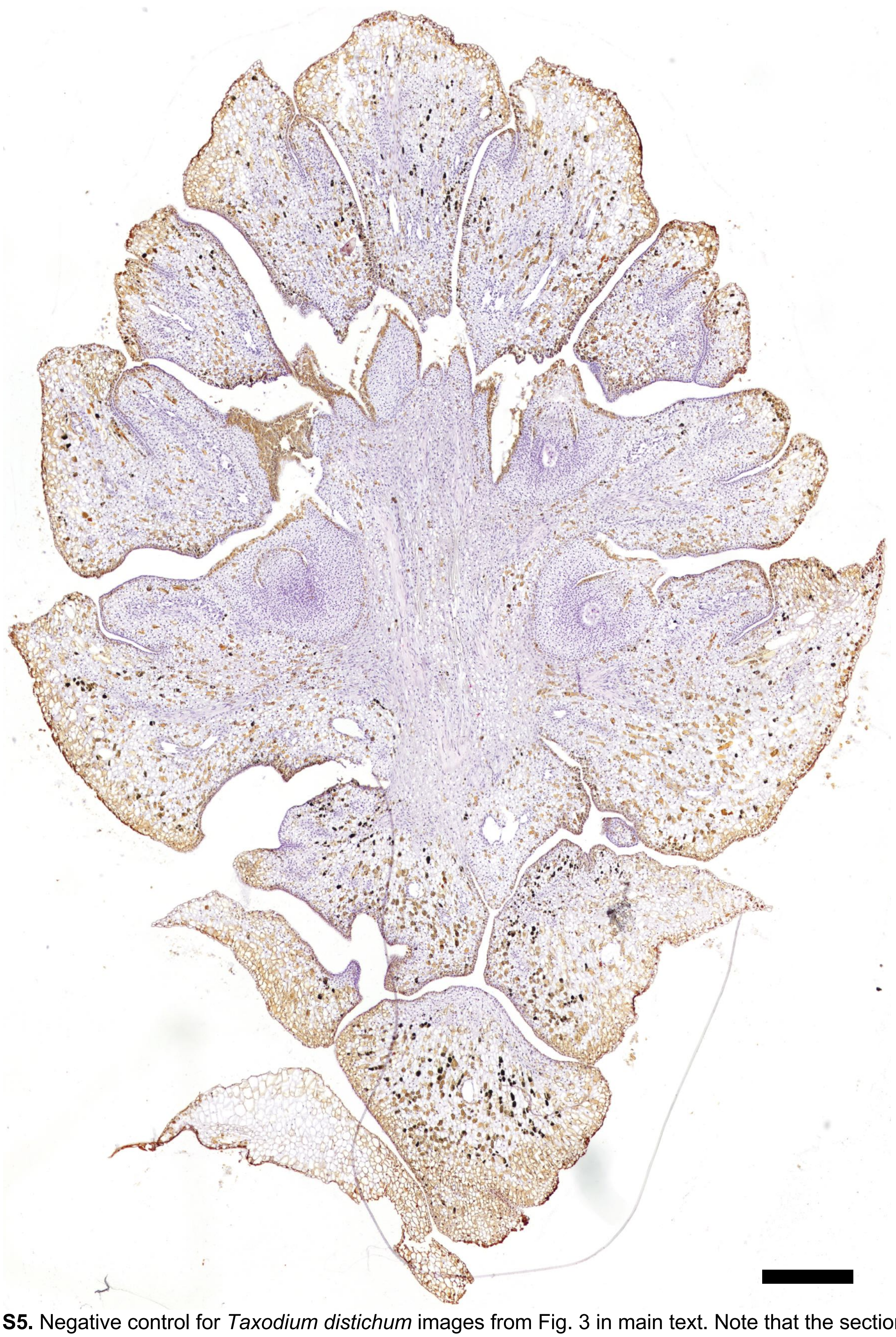

**Fig. S5.** Negative control for *Taxodium distichum* images from Fig. 3 in main text. Note that the section was counterstained with Gill's Hematoxylin. Scale bar = 500 µm.

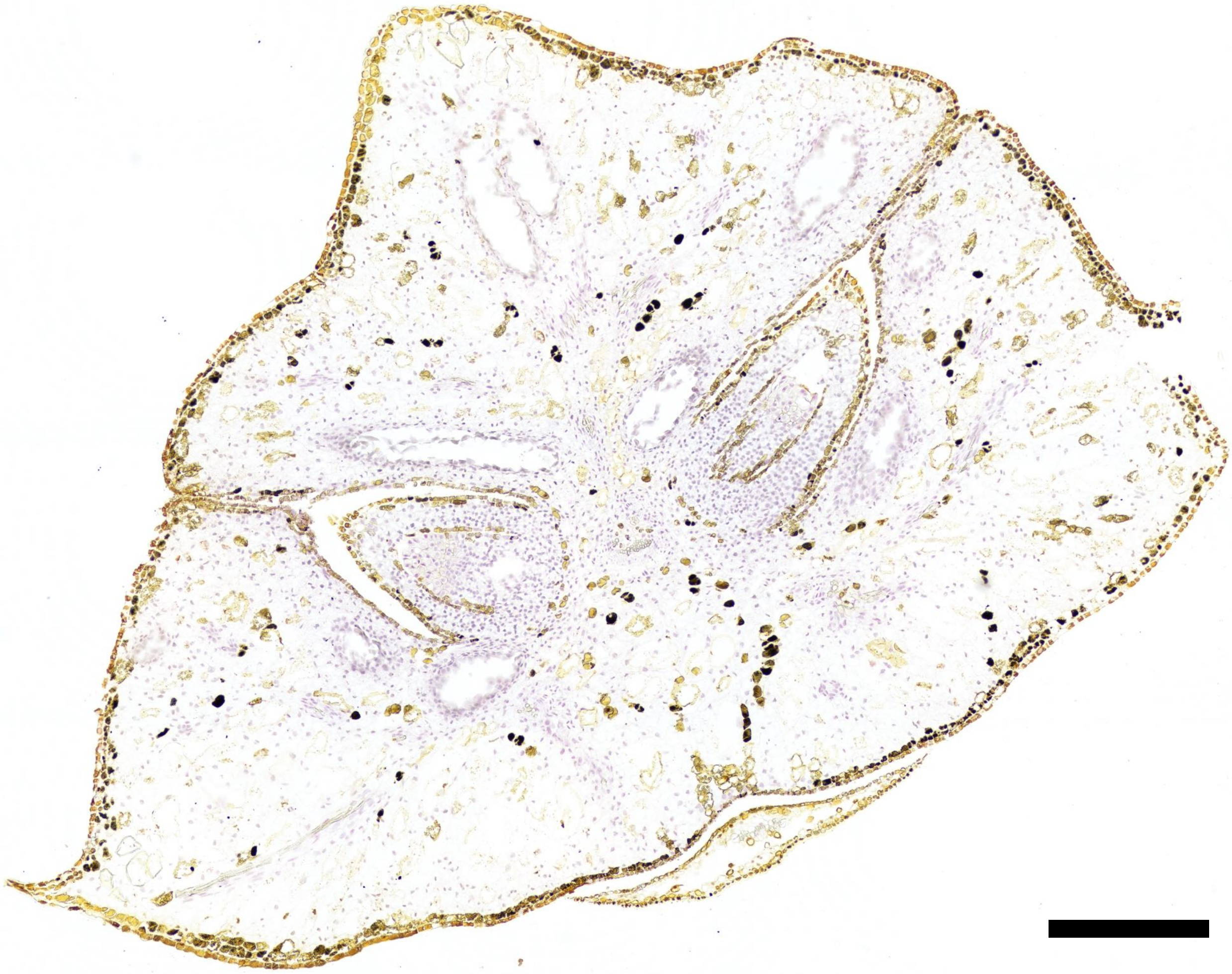

**Fig. S6.** Negative control for *Juniperus virginiana* images from Fig. 3 in main text. Note that the section was counterstained with Gill's Hematoxylin. Scale bar = 500 μm.

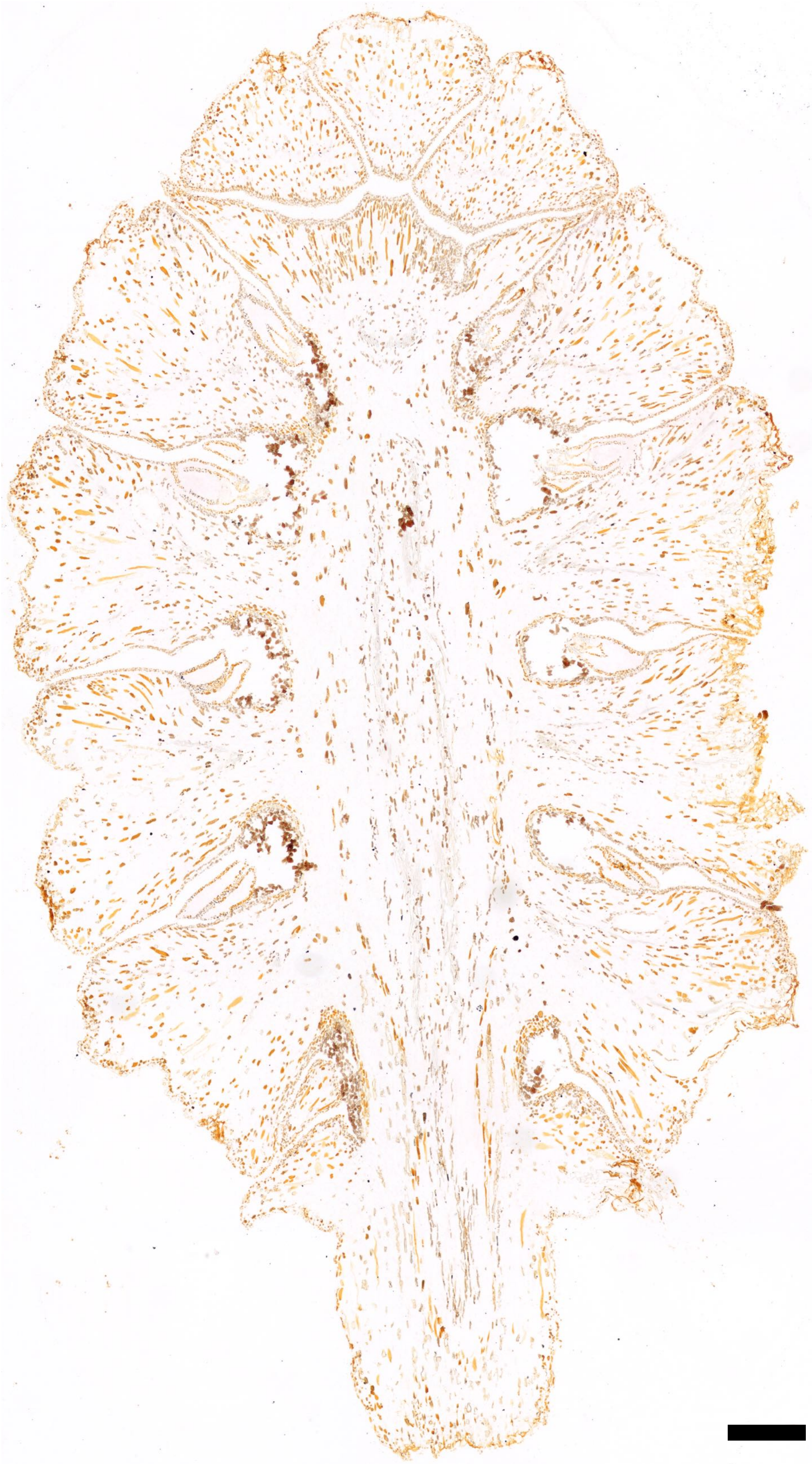

**Fig. S7.** Negative control for *Metasequoia glyptostroboides* images from Fig. 3 in main text. Scale bar = 500 µm.

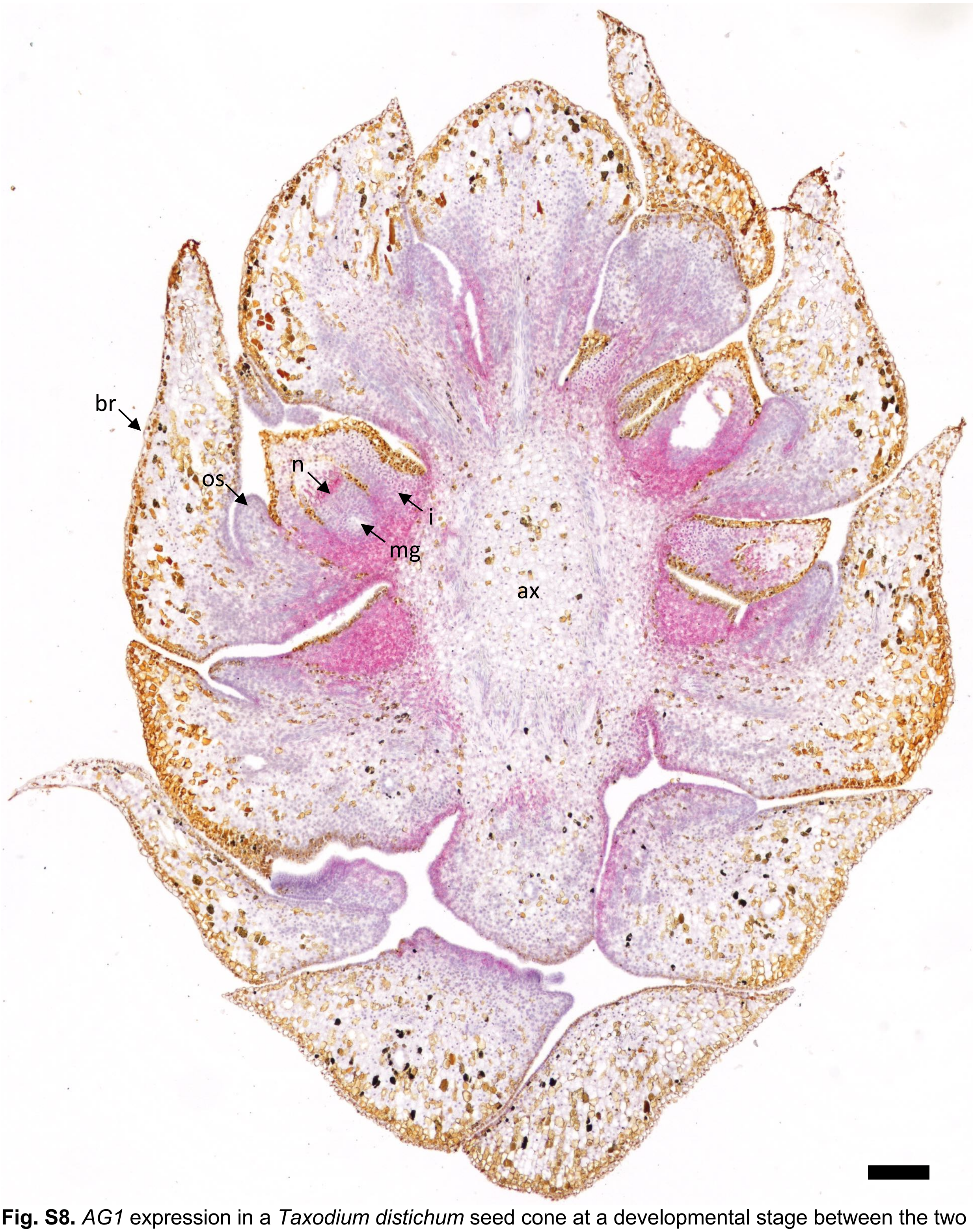


**Fig. S8.** *AG1* expression in a *Taxodium distichum* seed cone at a developmental stage between the two stages shown in Figs. 2 and 3 when the ovuliferous scale has begun to grow. Note that the section was counterstained with Gill's Hematoxylin and cut at 10 μm. br = bract; os = ovuliferous scale; n = nucellus; i = integument; mg = megagametophyte; ax = cone axis. Scale bar = 250 μm.

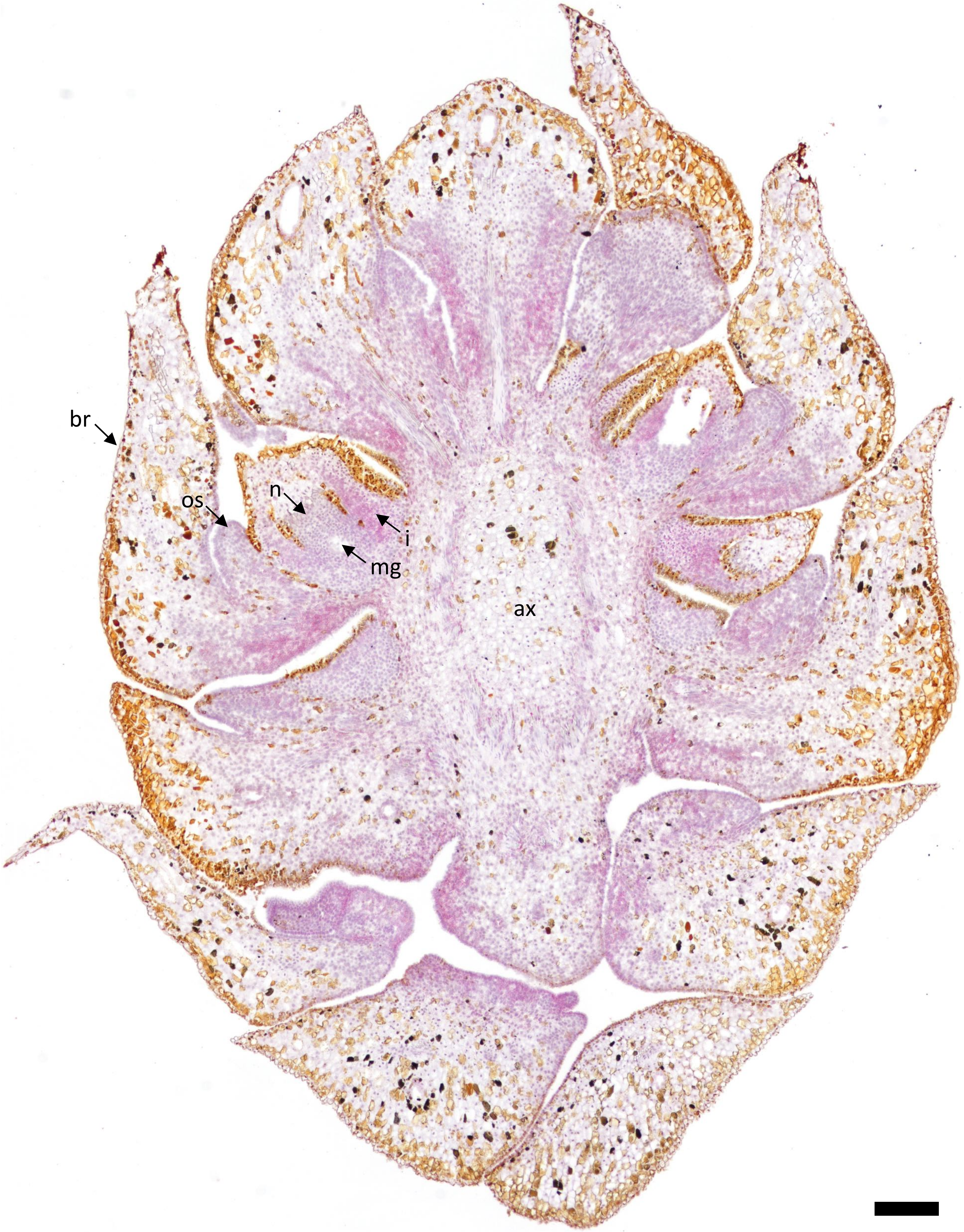


**Fig. S9.** *AG2* expression in a *Taxodium distichum* seed cone at a developmental stage between the two stages shown in Figs. 2 and 3 when the ovuliferous scale has begun to grow. Note that the section was counterstained with Gill's Hematoxylin and cut at 10 µm. br = bract; os = ovuliferous scale; n = nucellus; i = integument; mg = megagametophyte; ax = cone axis. Scale bar = 250 µm.

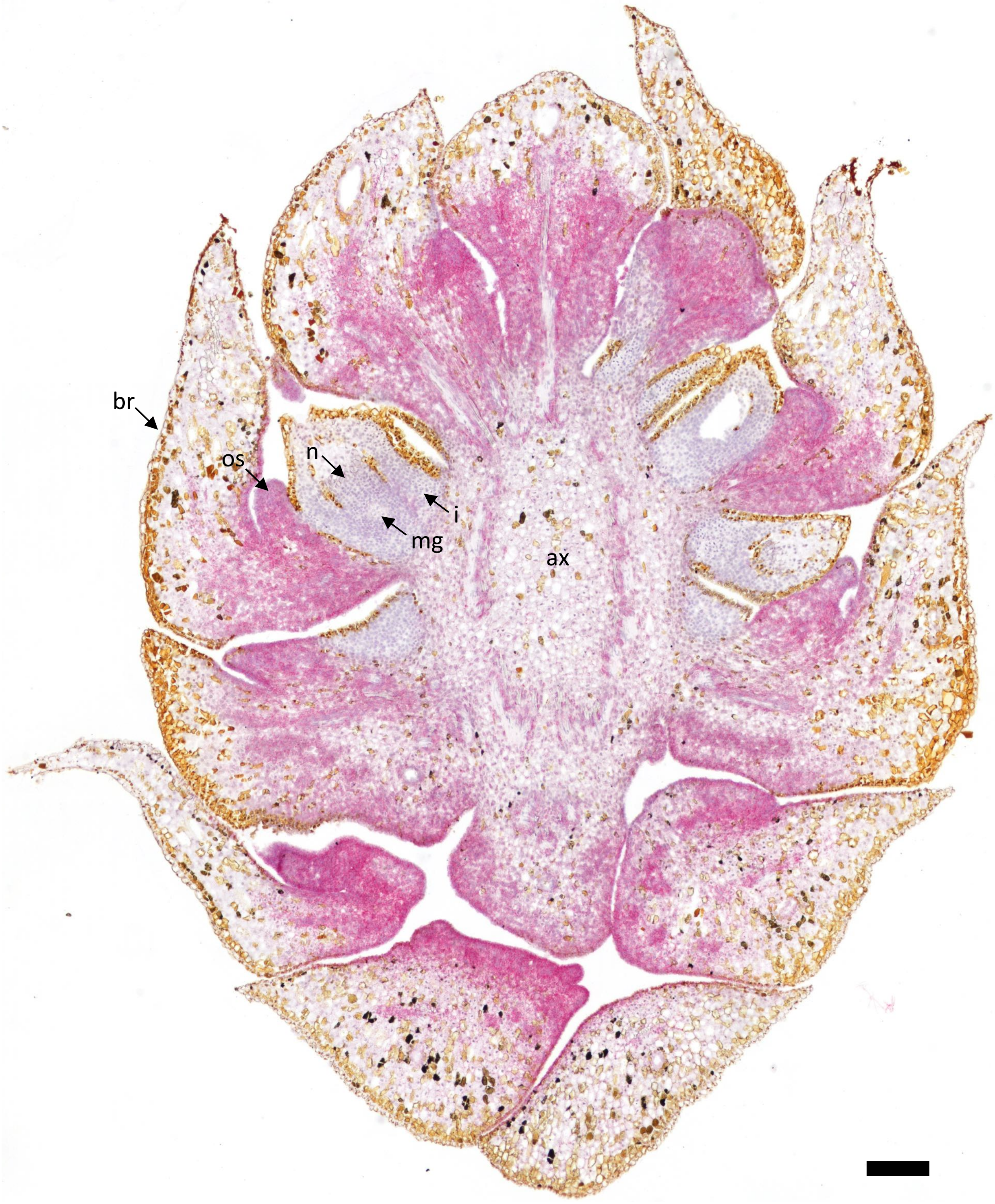


**Fig. S10.** *AG3* expression in a *Taxodium distichum* seed cone at a developmental stage between the two stages shown in Figs. 2 and 3 when the ovuliferous scale has begun to grow. Note that the section was counterstained with Gill's Hematoxylin and cut at 10 µm. br = bract; os = ovuliferous scale; n = nucellus; i = integument; mg = megagametophyte; ax = cone axis. Scale bar = 250 µm.

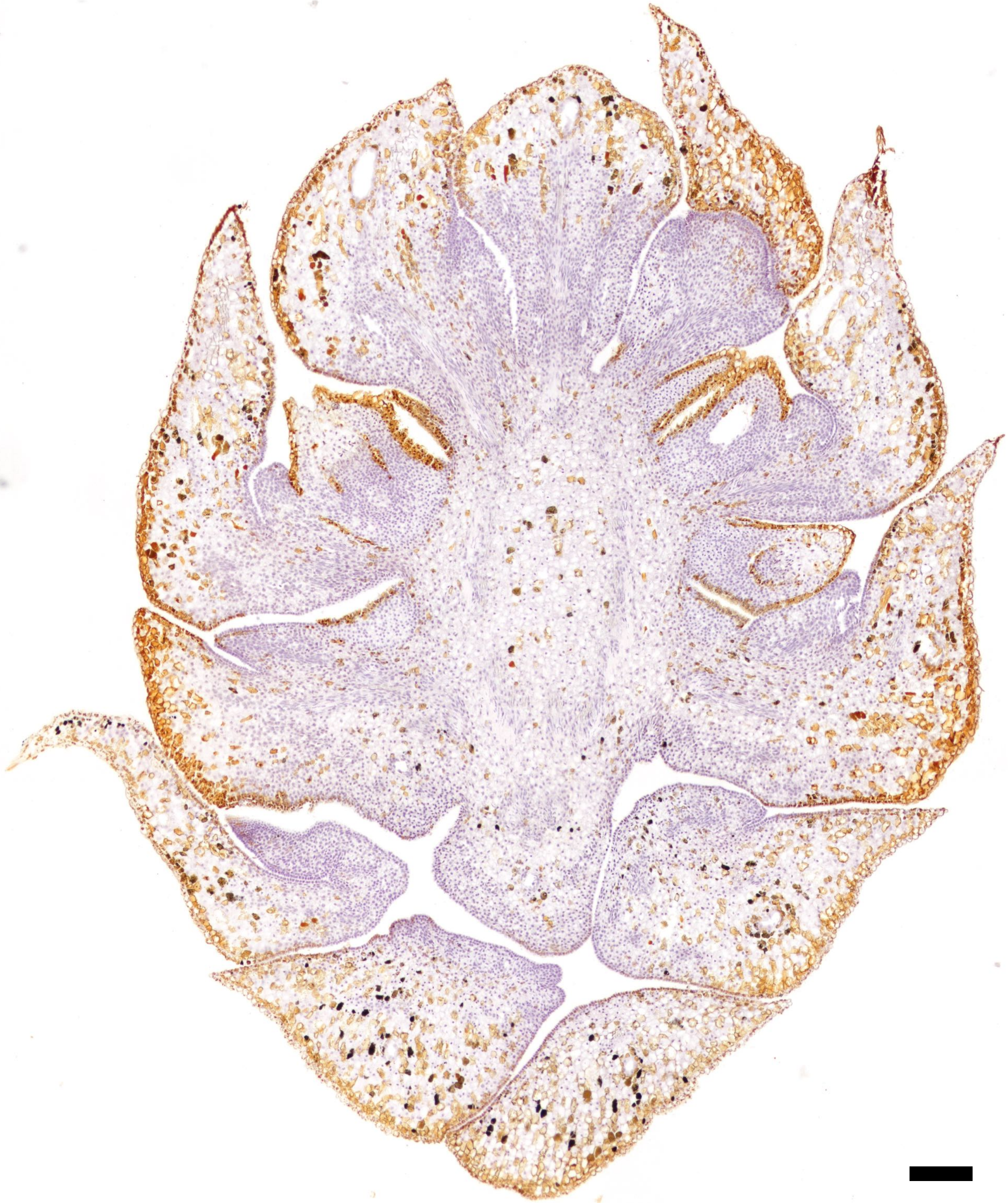

**Fig. S11.** Negative control for *Taxodium distichum* seed cone sections at a developmental stage between the two stages shown in Figs. 2 and 3 when the ovuliferous scale has begun to grow. Note that the section was counterstained with Gill's Hematoxylin and cut at 10 μm. Scale bar = 250 μm.

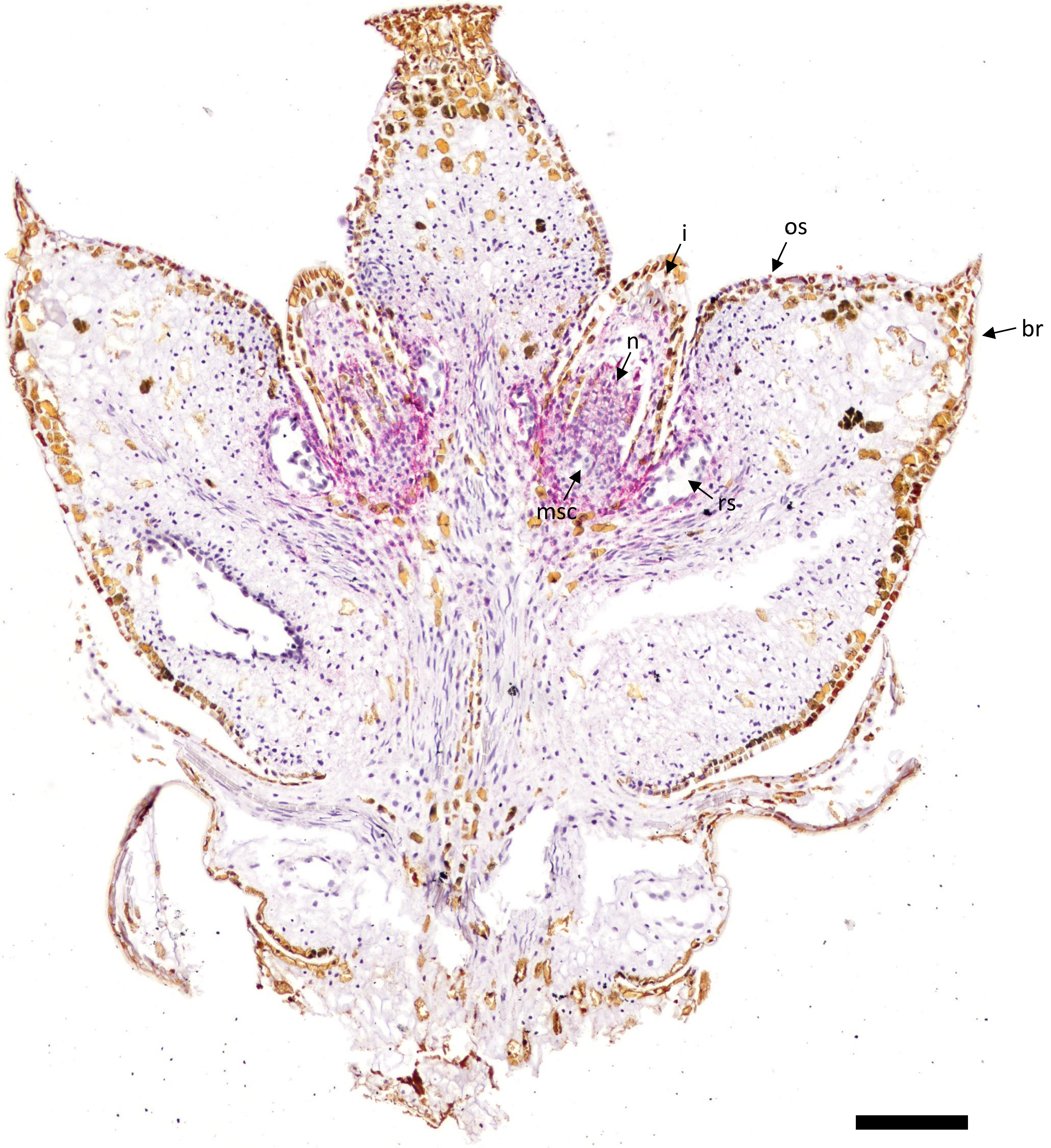


**Fig. S12.** *AG1* expression in a *Juniperus virginiana* seed cone at a developmental stage between the two stages shown in Figs. 2 and 3 when the ovuliferous scale has begun to grow. Note that the section was counterstained with Gill's Hematoxylin. br = bract; os = ovuliferous scale; n = nucellus; i = integument; msc = megasporocyte; rs = resin canal. Scale bar = 250 μm.

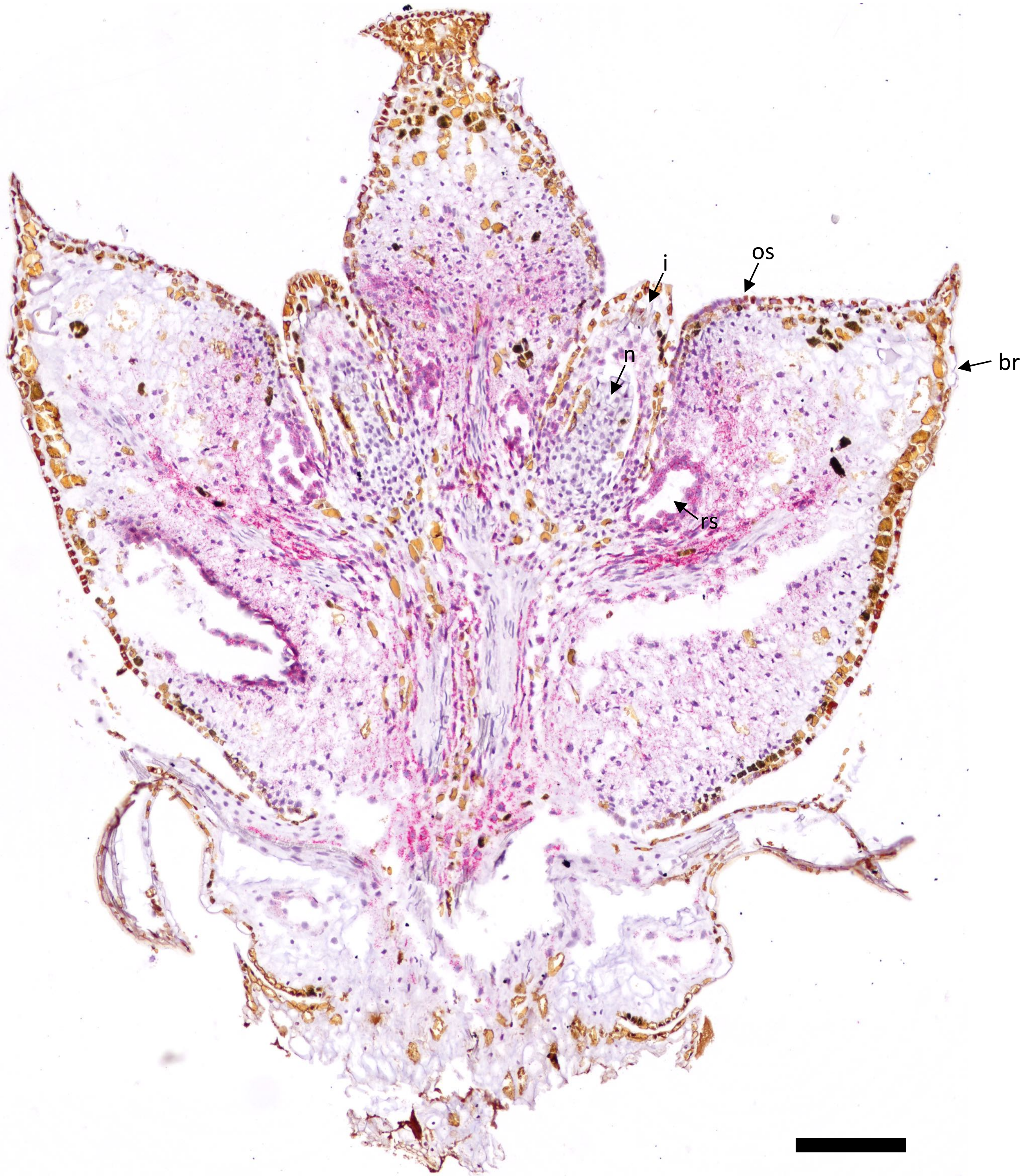


**Fig. S13.** *AG2* expression in a *Juniperus virginiana* seed cone at a developmental stage between the two stages shown in Figs. 2 and 3 when the ovuliferous scale has begun to grow. Note that this section does not contain the megasporocyte and was counterstained with Gill's Hematoxylin. br = bract; os = ovuliferous scale; n = nucellus; i = integument; rs = resin canal. Scale bar = 250 μm.

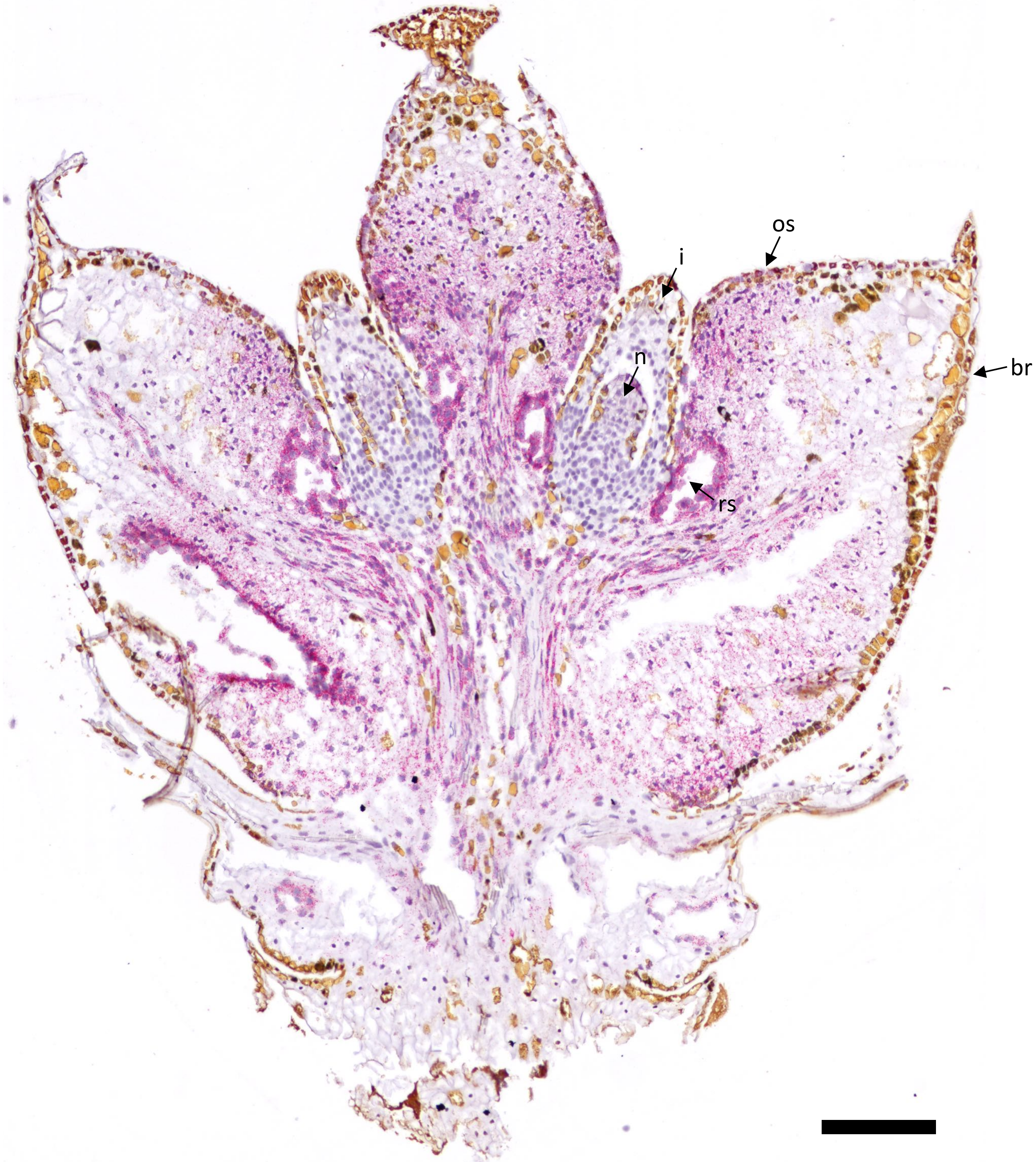


**Fig. S14.** *AG3* expression in a *Juniperus virginiana* seed cone at a developmental stage between the two stages shown in Figs. 2 and 3 when the ovuliferous scale has begun to grow. Note that this section does not contain the megasporocyte and was counterstained with Gill's Hematoxylin. br = bract; os = ovuliferous scale; n = nucellus; i = integument; rs = resin canal. Scale bar = 250 µm.

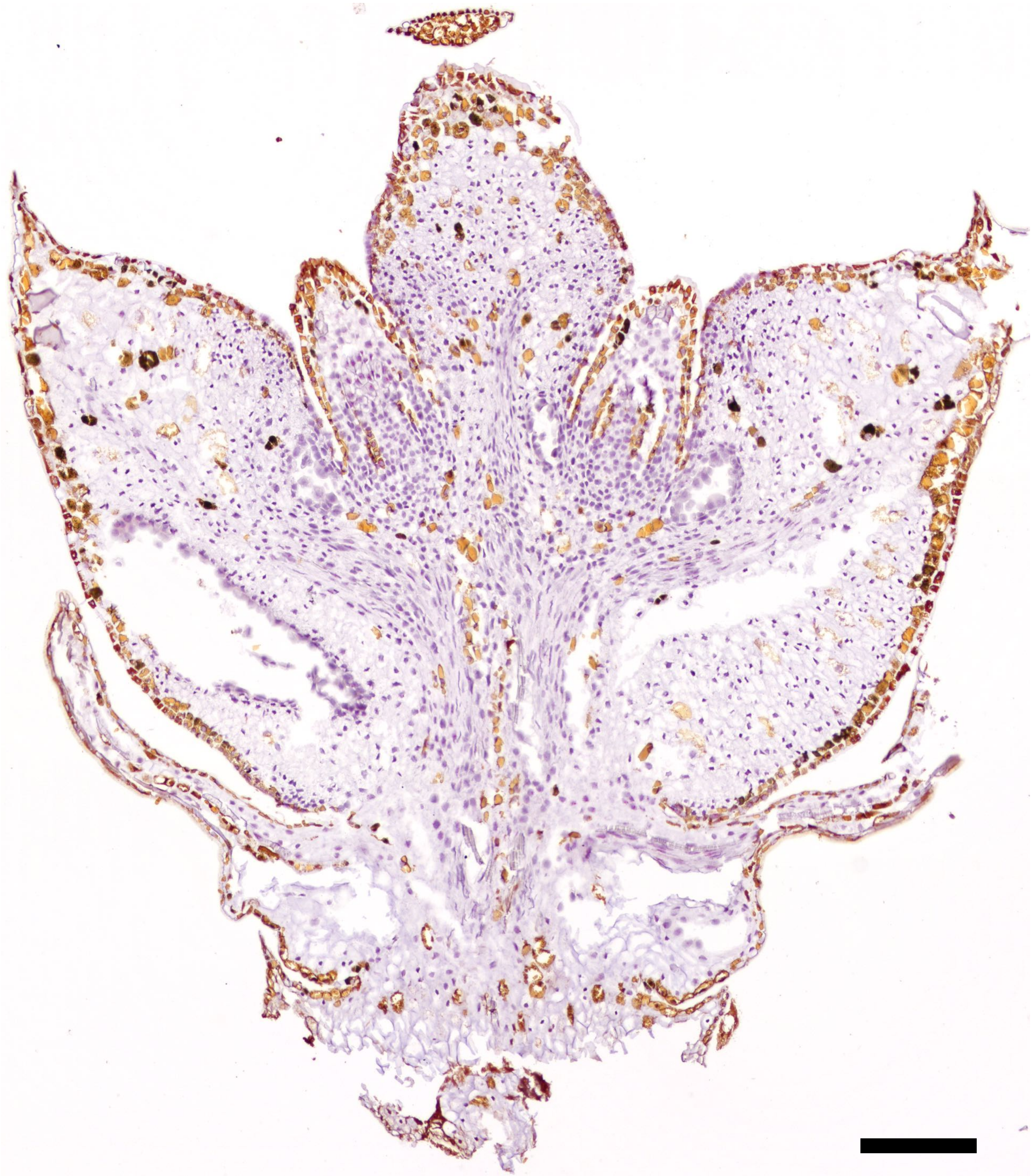

**Fig. S15.** Negative control for *Juniperus virginiana* seed cone sections at a developmental stage between the two stages shown in Figs. 2 and 3 when the ovuliferous scale has begun to grow. Note that this section does not contain the megasporocyte and was counterstained with Gill's Hematoxylin. Scale bar = 250 μm.

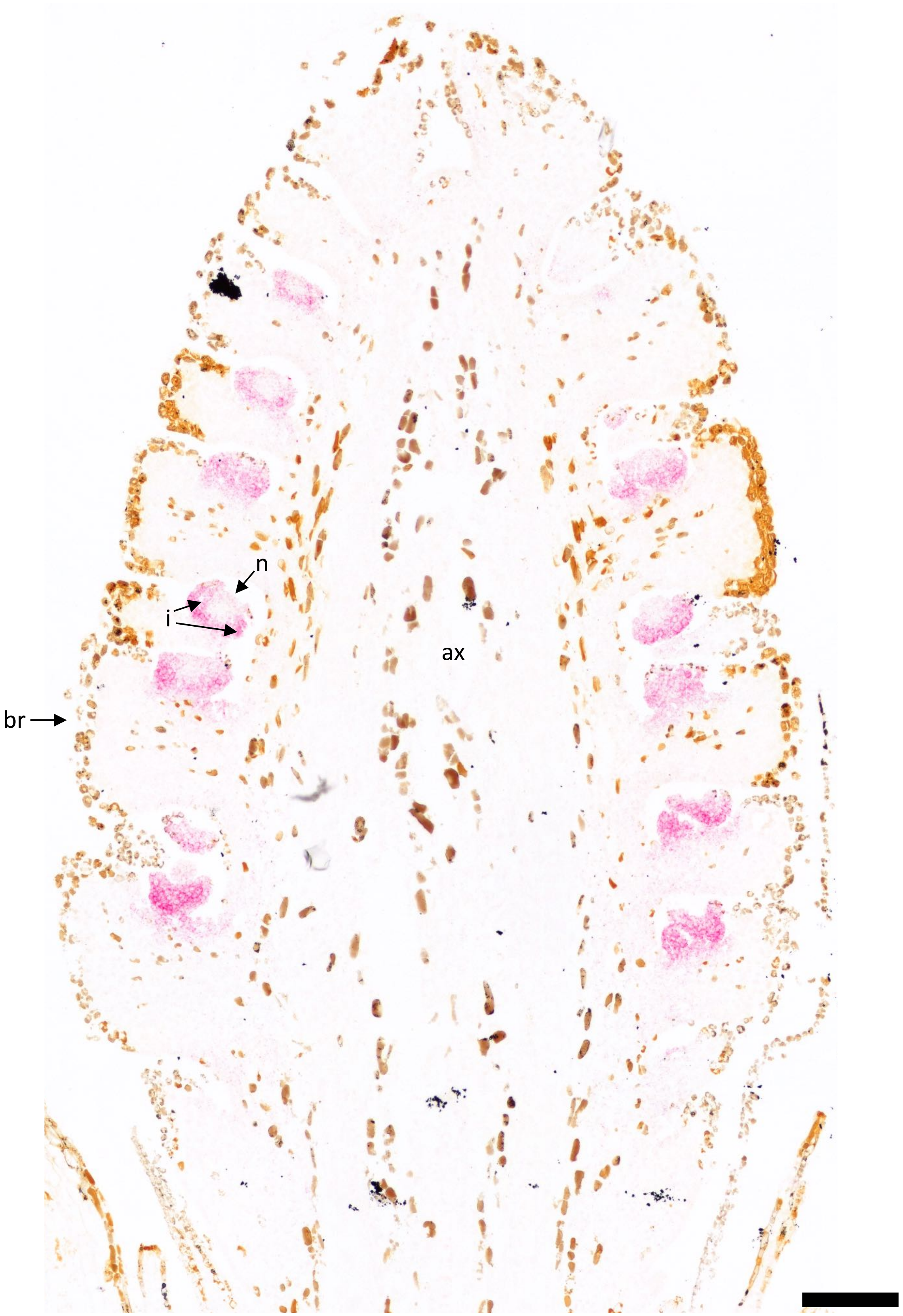


**Fig. S16.** *AG1* expression in a *Metasequoia glyptostroboides* seed cone at a developmental stage earlier than that shown in Fig. 2. At this stage, the nucellus and integument are primordial, and no ovuliferous scale is apparent. br = bract; n = nucellus; i = integument; ax = cone axis. Scale bar = 200 μm.

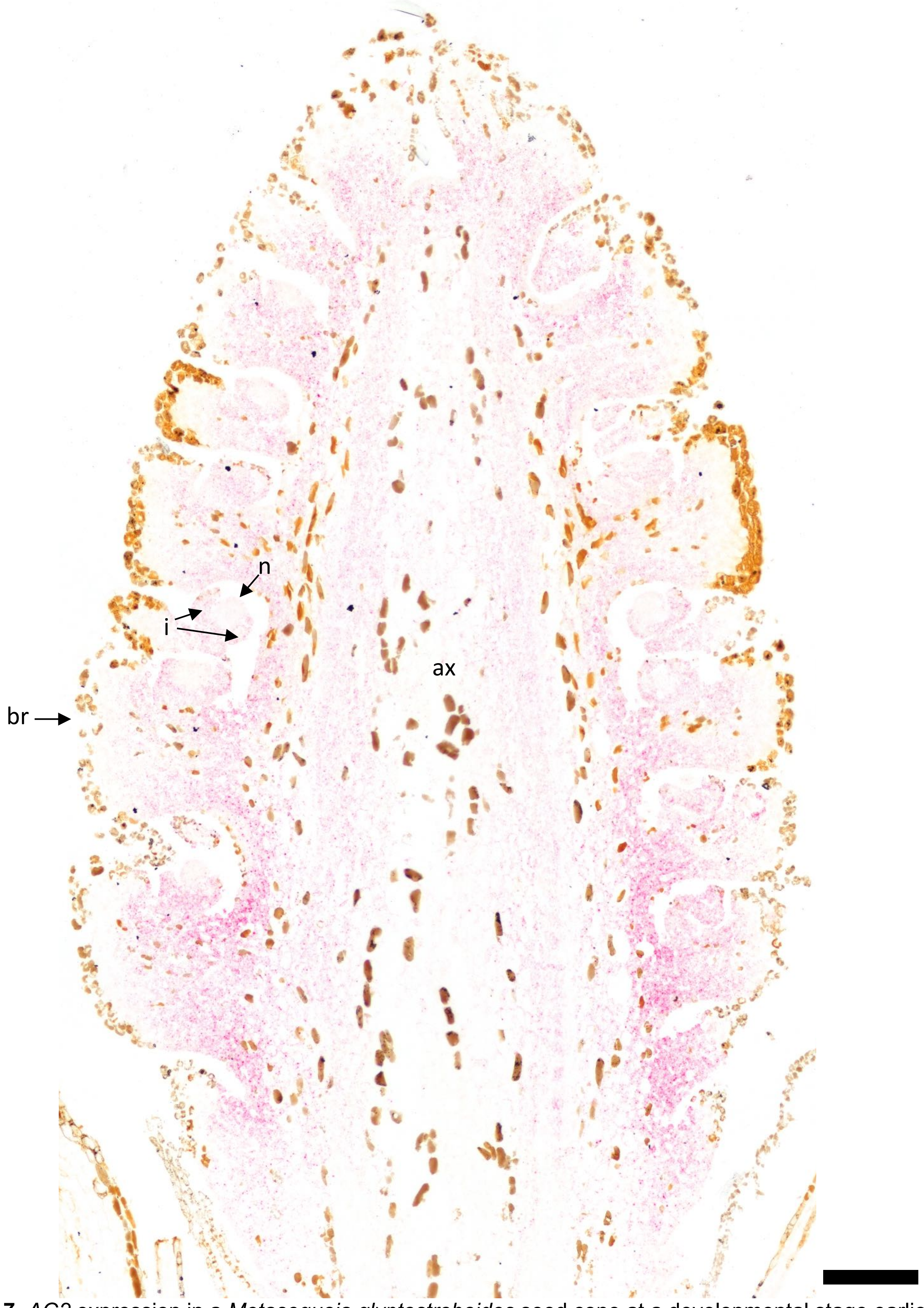


**Fig. S17.** *AG2* expression in a *Metasequoia glyptostroboides* seed cone at a developmental stage earlier than that shown in Fig. 2. At this stage, the nucellus and integument are primordial, and no ovuliferous scale is apparent. br = bract; n = nucellus; i = integument; ax = cone axis. Scale bar = 200 μm.

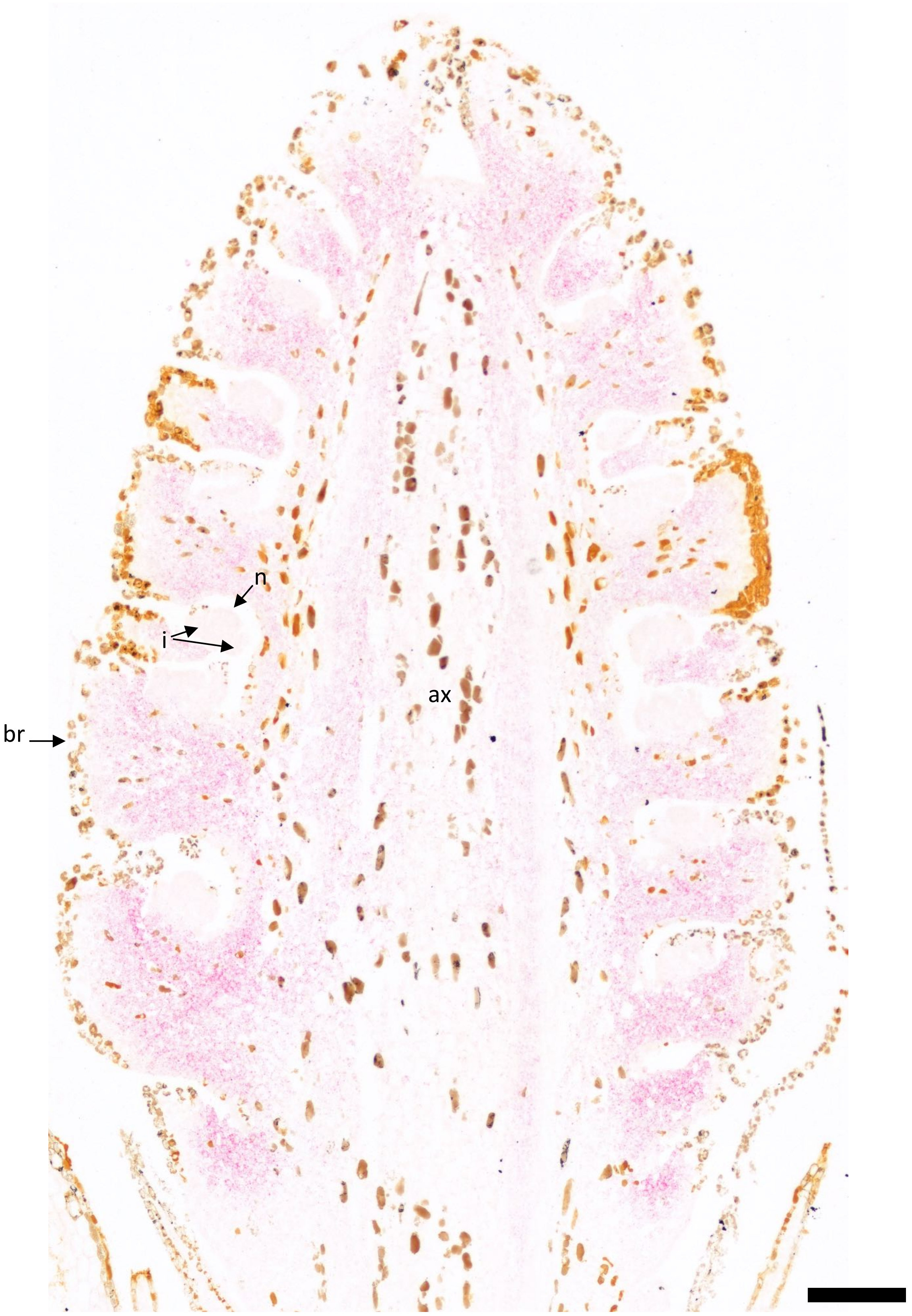


**Fig. S18.** *AG3* expression in a *Metasequoia glyptostroboides* seed cone at a developmental stage earlier than that shown in Fig. 2. At this stage, the nucellus and integument are primordial, and no ovuliferous scale is apparent. br = bract; n = nucellus; i = integument; ax = cone axis. Scale bar = 200 μm.

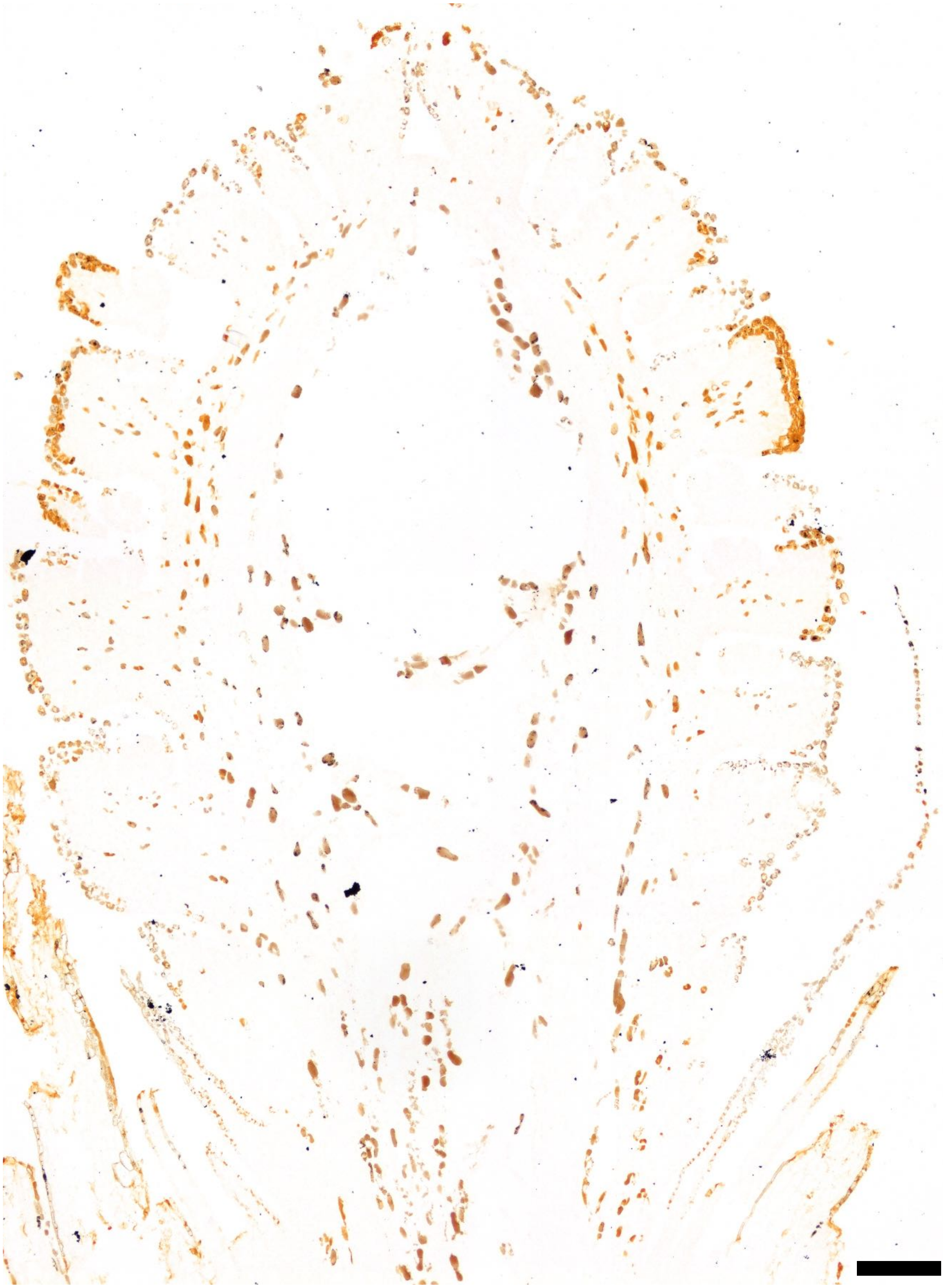

**Fig. S19.** Negative control for *Metasequoia glyptostroboides* seed cone sections at a developmental stage earlier than that shown in Fig. 2. At this stage, the ovules are developing, specifically the integument is starting to grow around the nucellus, and no ovuliferous scale is apparent. Scale bar = 200 μm.

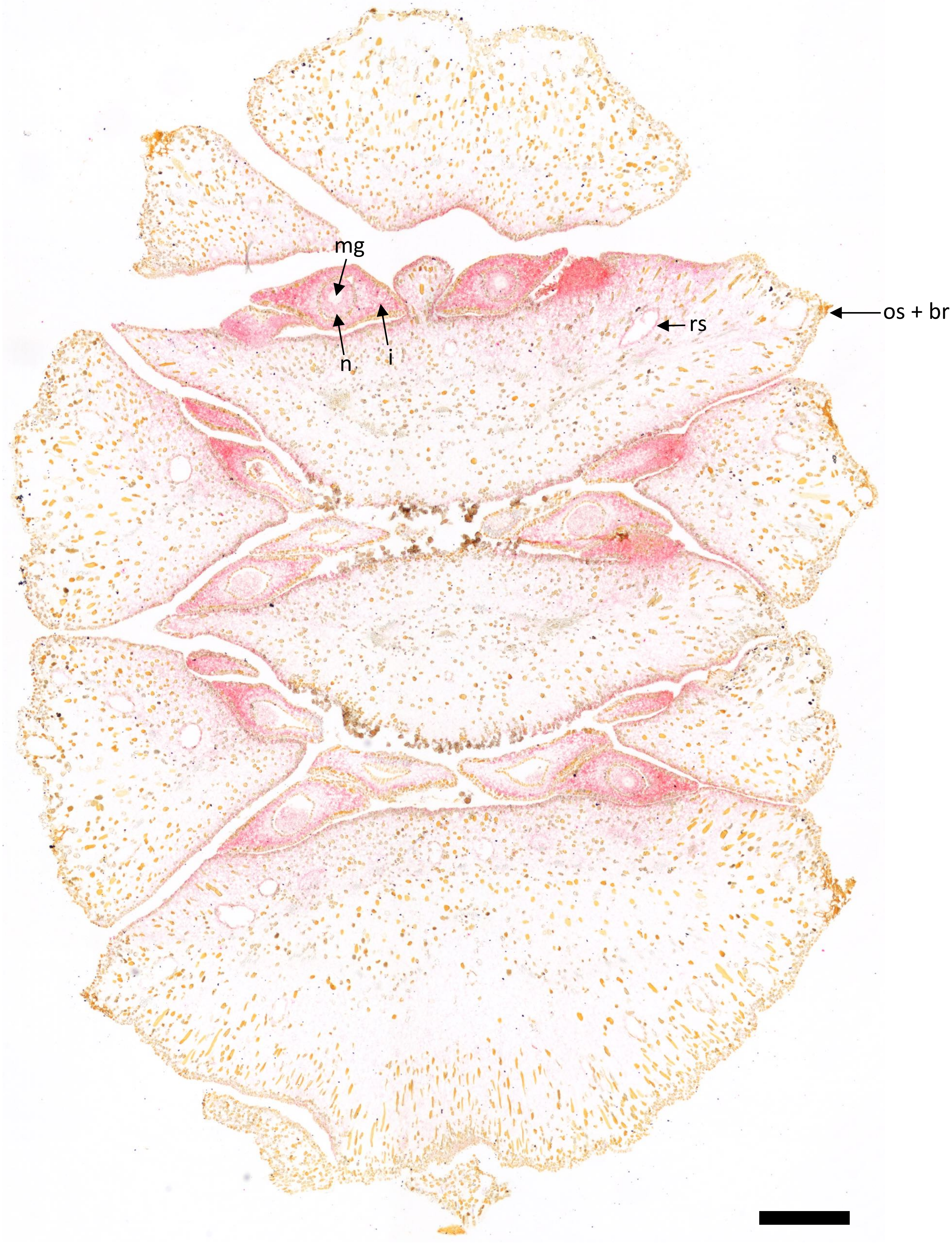


**Fig. S20.** *AG1* expression in a tangential section of a more mature seed cone (transverse through the bracts, ovuliferous scales, and ovules) of *Metasequoia glyptostroboides*. br = bract; os = ovuliferous scale; n = nucellus; i = integument; mg = megagametophyte; rs = resin canal. Scale bar = 500 µm.

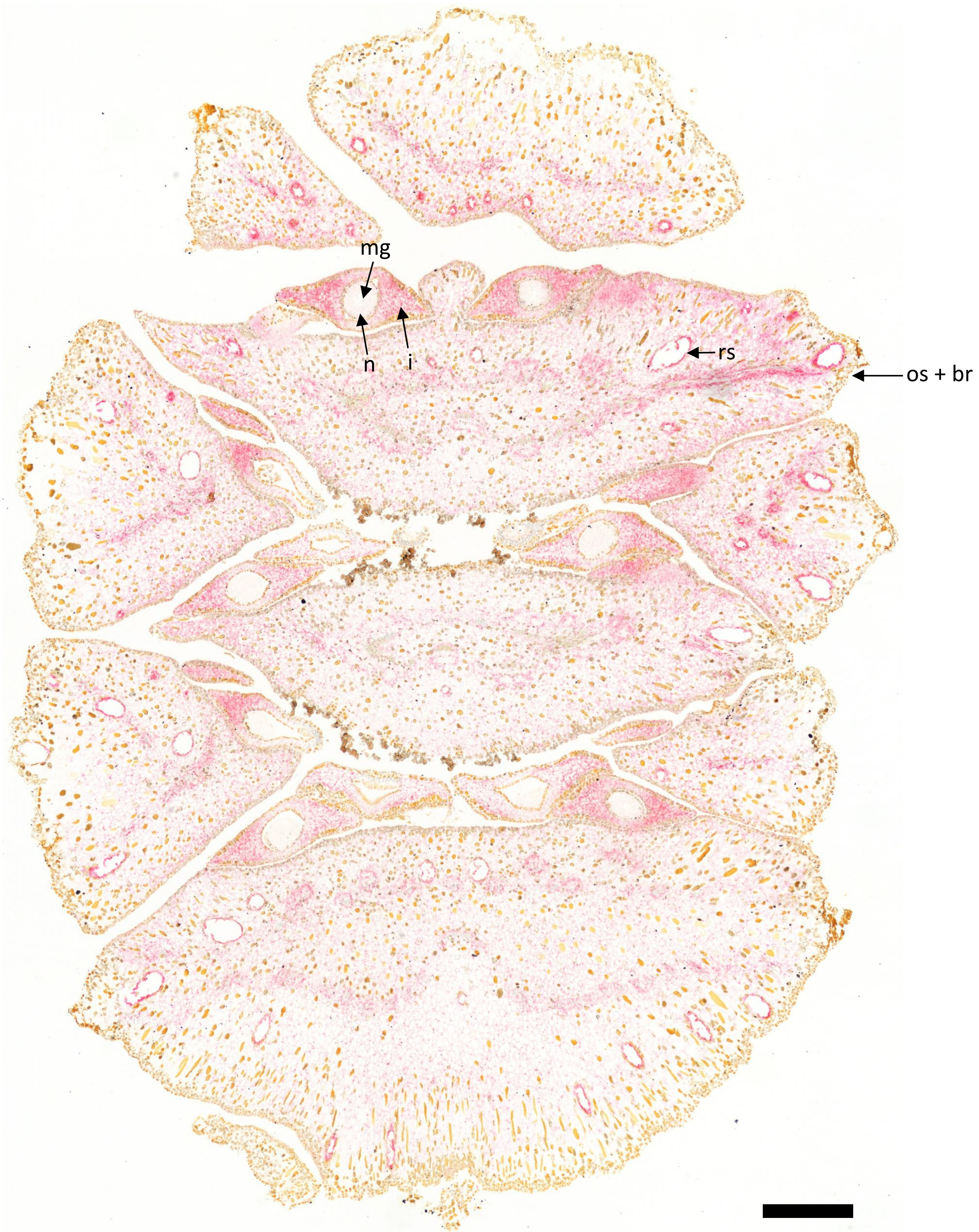


**Fig. S21.** *AG2* expression in a tangential section of a more mature seed cone (transverse through the bracts, ovuliferous scales, and ovules) of *Metasequoia glyptostroboides*. br = bract; os = ovuliferous scale; n = nucellus; i = integument; mg = megagametophyte; rs = resin canal. Scale bar = 500 µm.

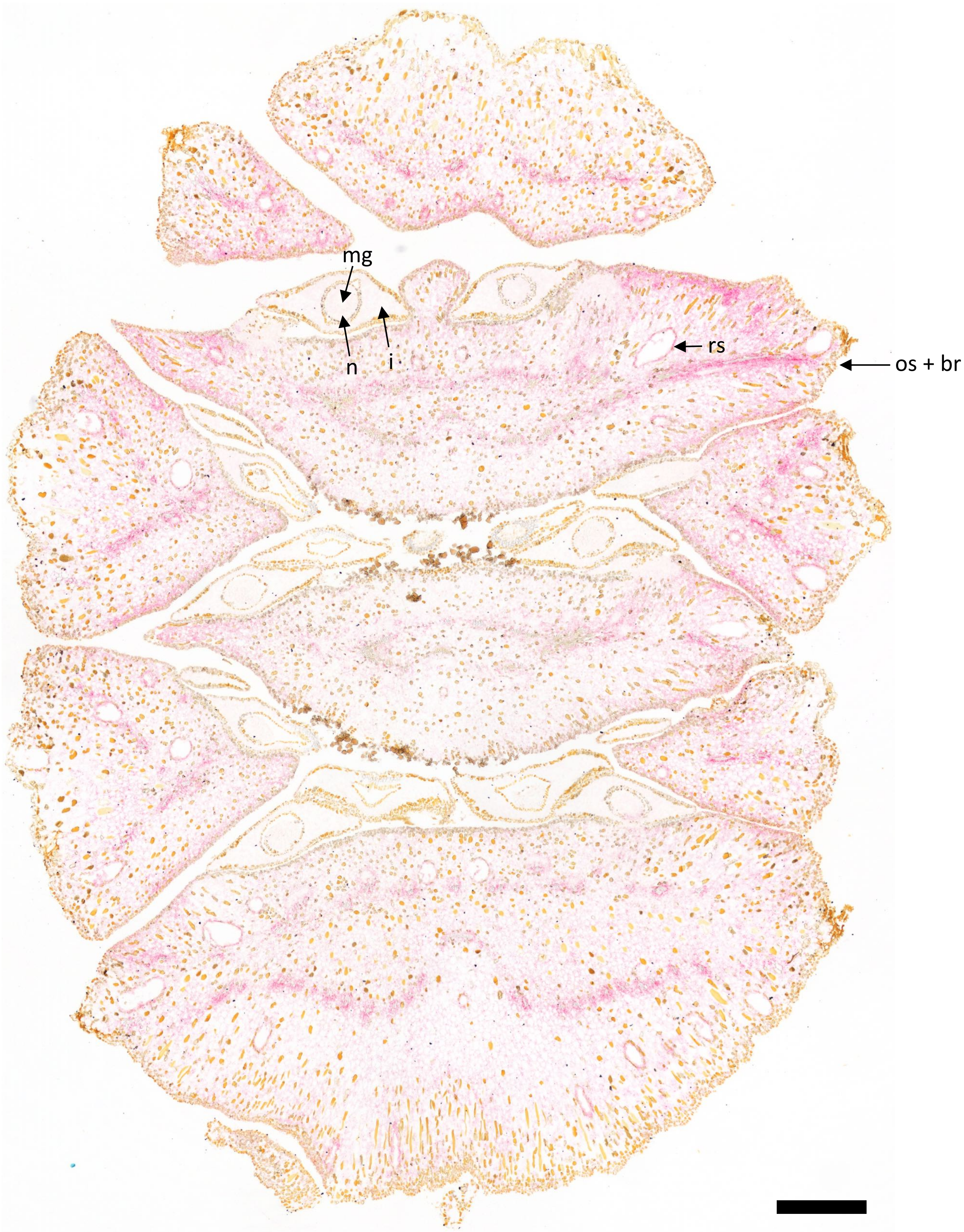


**Fig. S22.** *AG3* expression in a tangential section of a more mature seed cone (transverse through the bracts, ovuliferous scales, and ovules) of *Metasequoia glyptostroboides*. br = bract; os = ovuliferous scale; n = nucellus; i = integument; mg = megagametophyte; rs = resin canal. Scale bar = 500 µm.

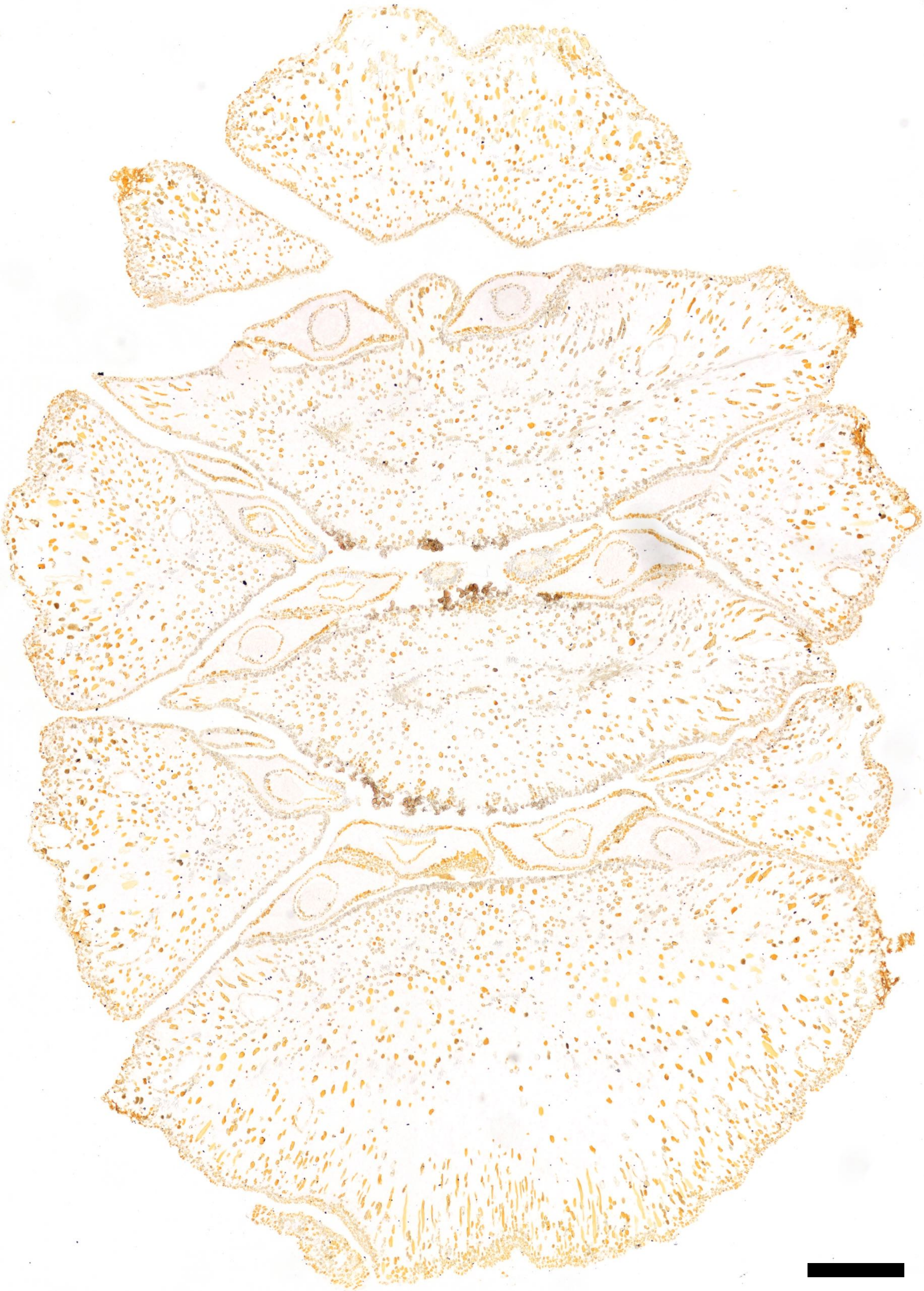

**Fig. S23.** Negative control for tangential sections of a more mature seed cone (transverse through the bracts, ovuliferous scales, and ovules) of *Metasequoia glyptostroboides*. Scale bar = 500 µm.

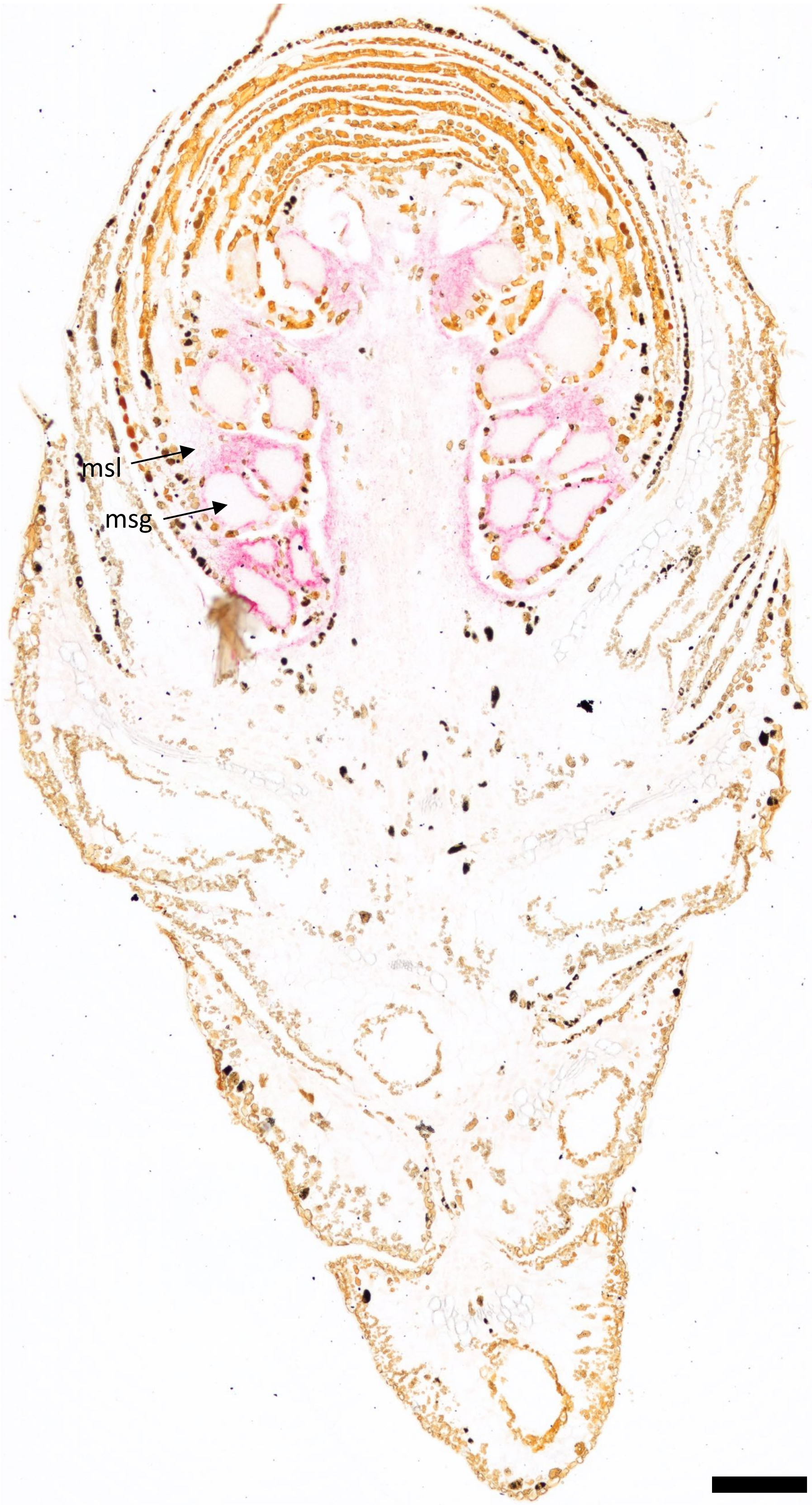


**Fig. S24.** *AG1* expression in *Taxodium distichum* pollen cone. msl = microsporophyll; msg = microsporangium. Scale bar = 200 μm.

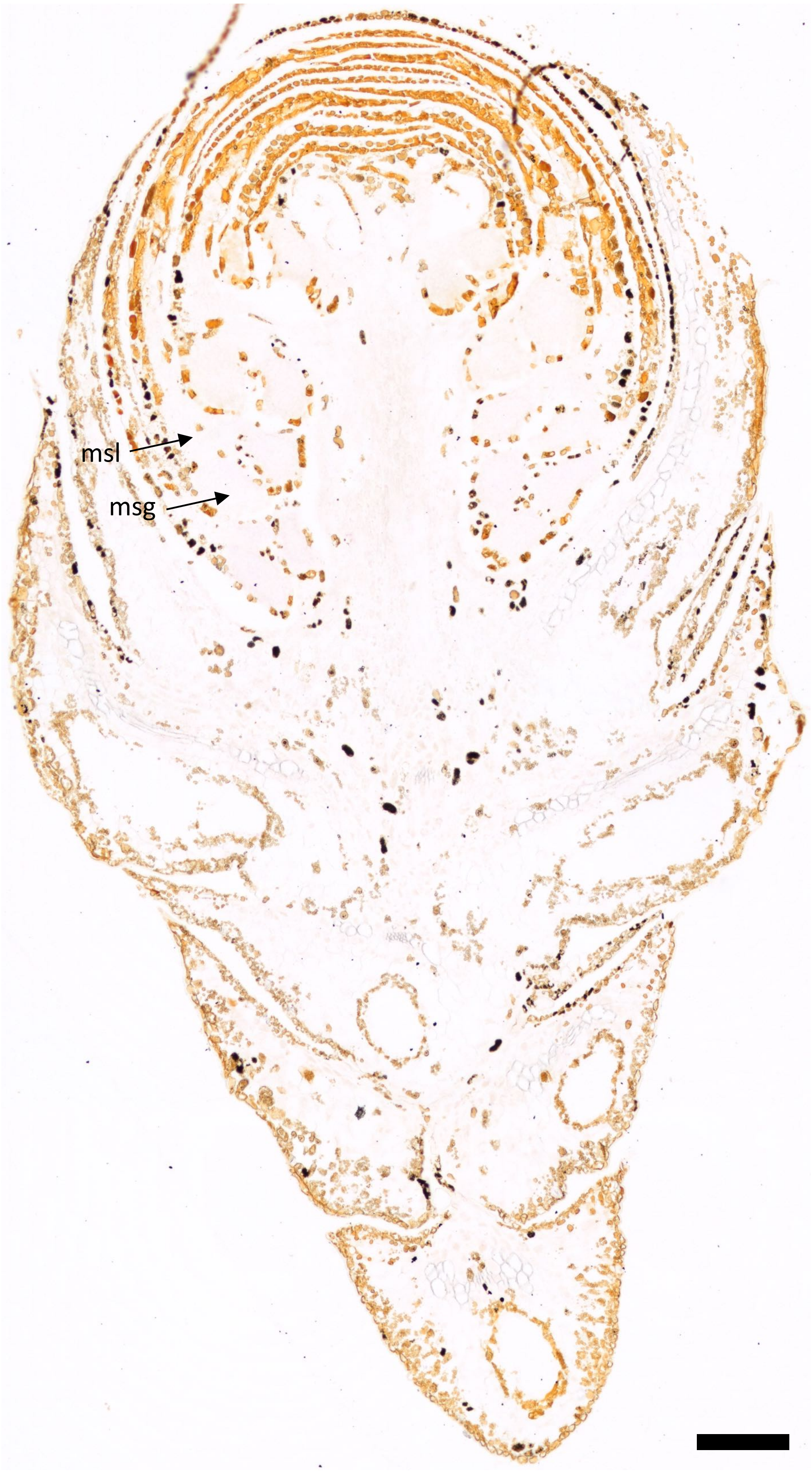


**Fig. S25.** *AG2* expression in *Taxodium distichum* pollen cone. msl = microsporophyll; msg = microsporangium. Scale bar = 200 µm.

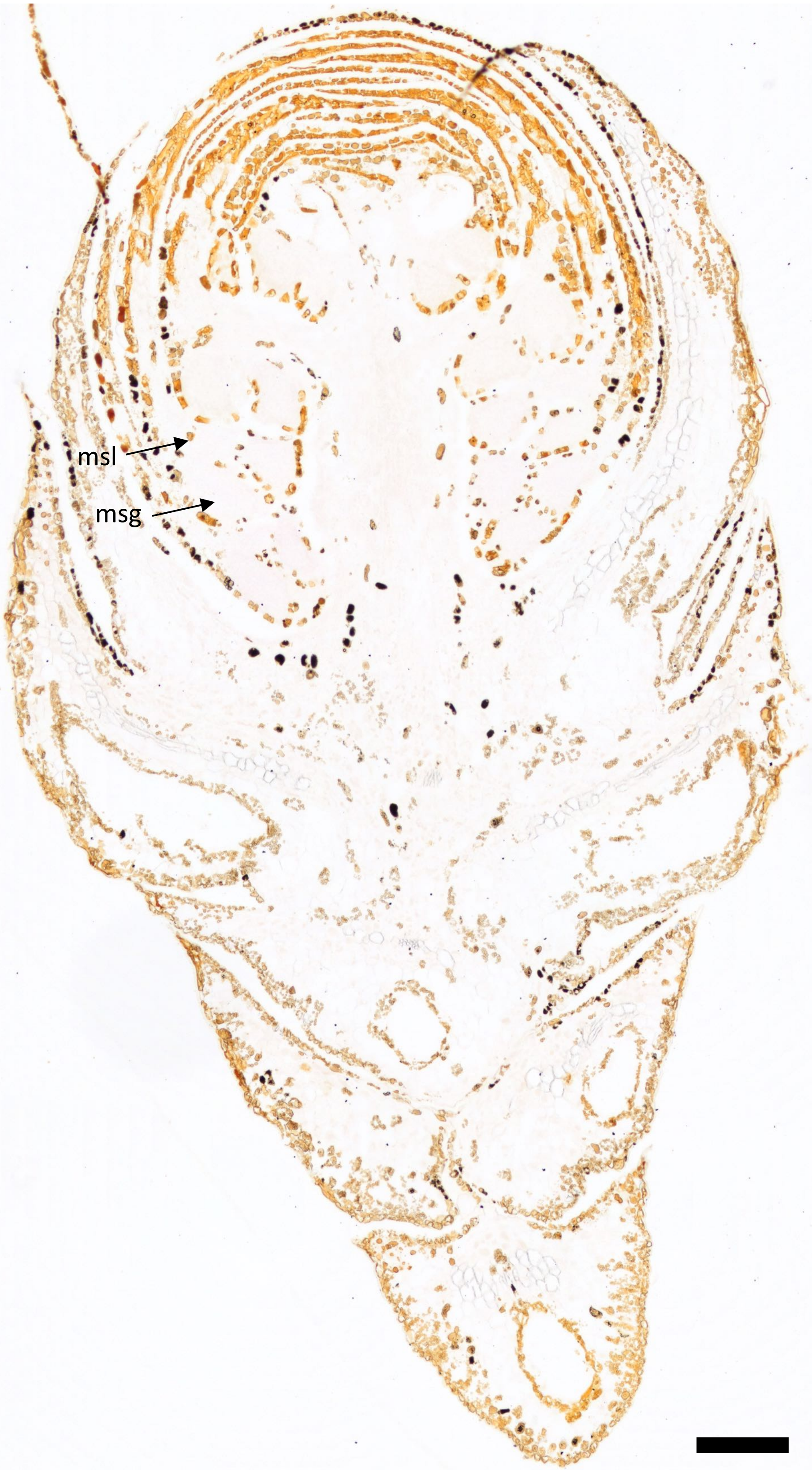


**Fig. S26.** *AG3* expression in *Taxodium distichum* pollen cone. msl = microsporophyll; msg = microsporangium. Scale bar = 200 μm.

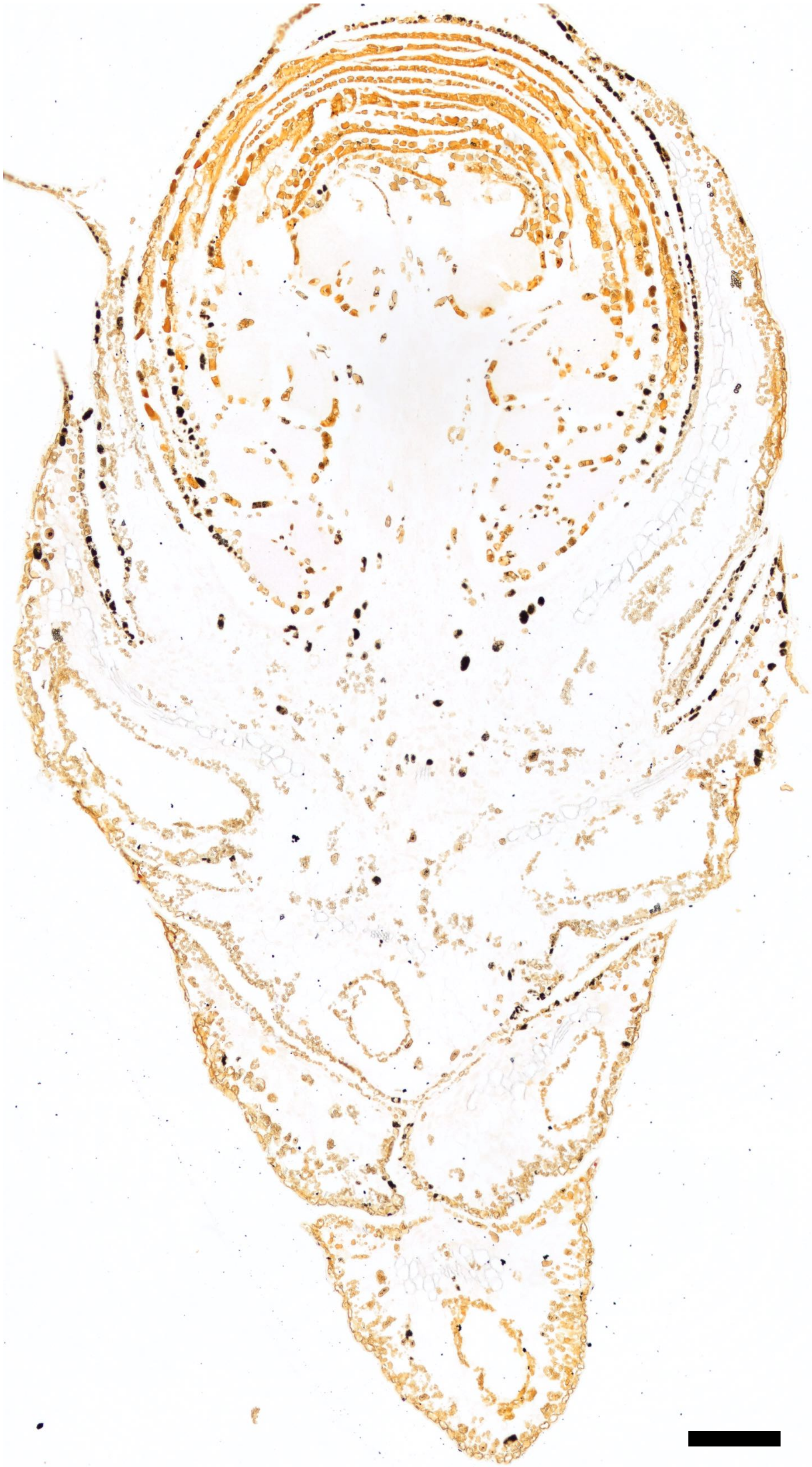

**Fig. S27.** Negative control for *Taxodium distichum* pollen cone. Scale bar = 200 μm.

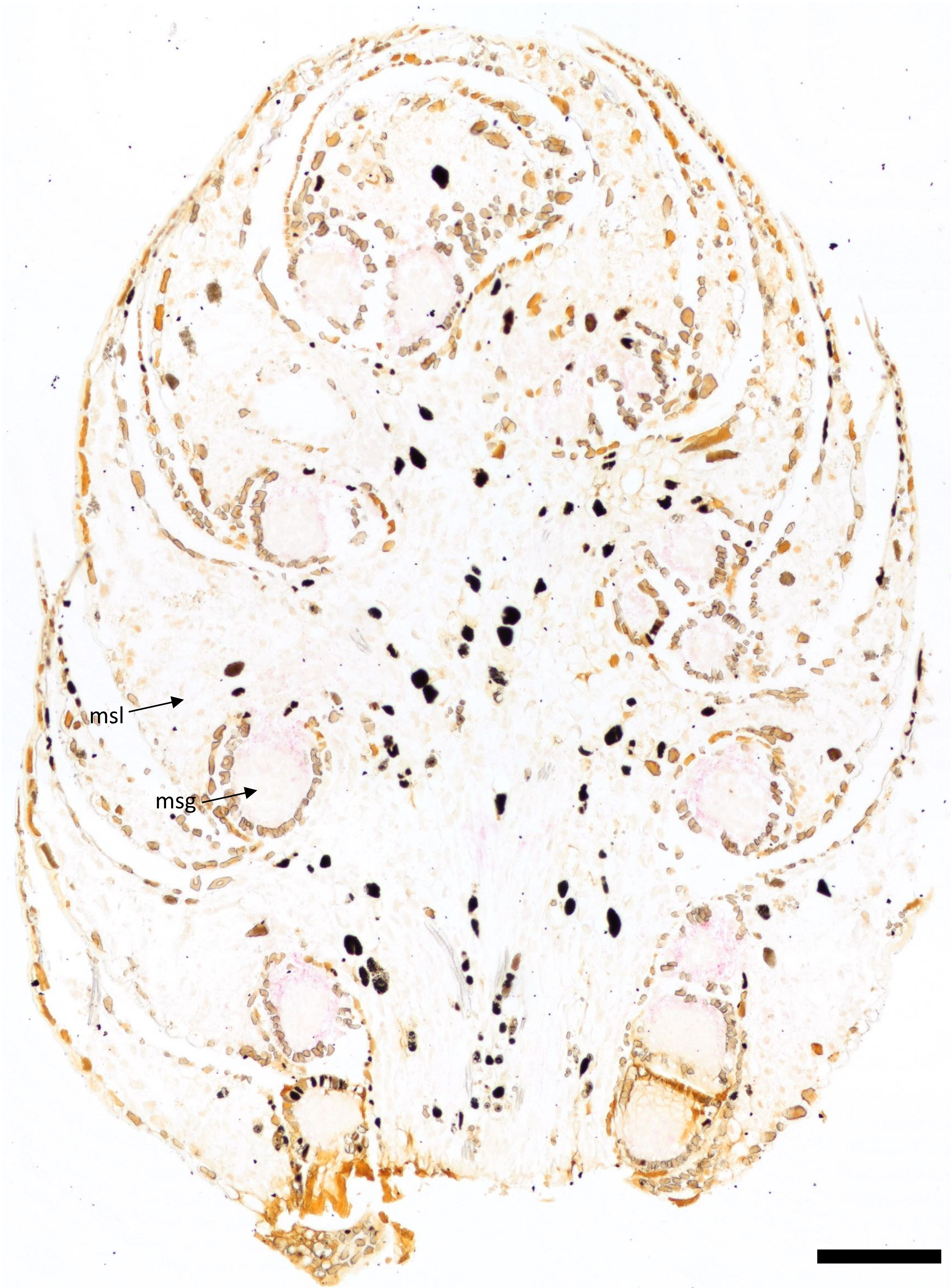


**Fig. S28.** *AG1* expression in *Juniperus virginiana* pollen cone. msl = microsporophyll; msg = microsporangium. Scale bar = 200 µm.

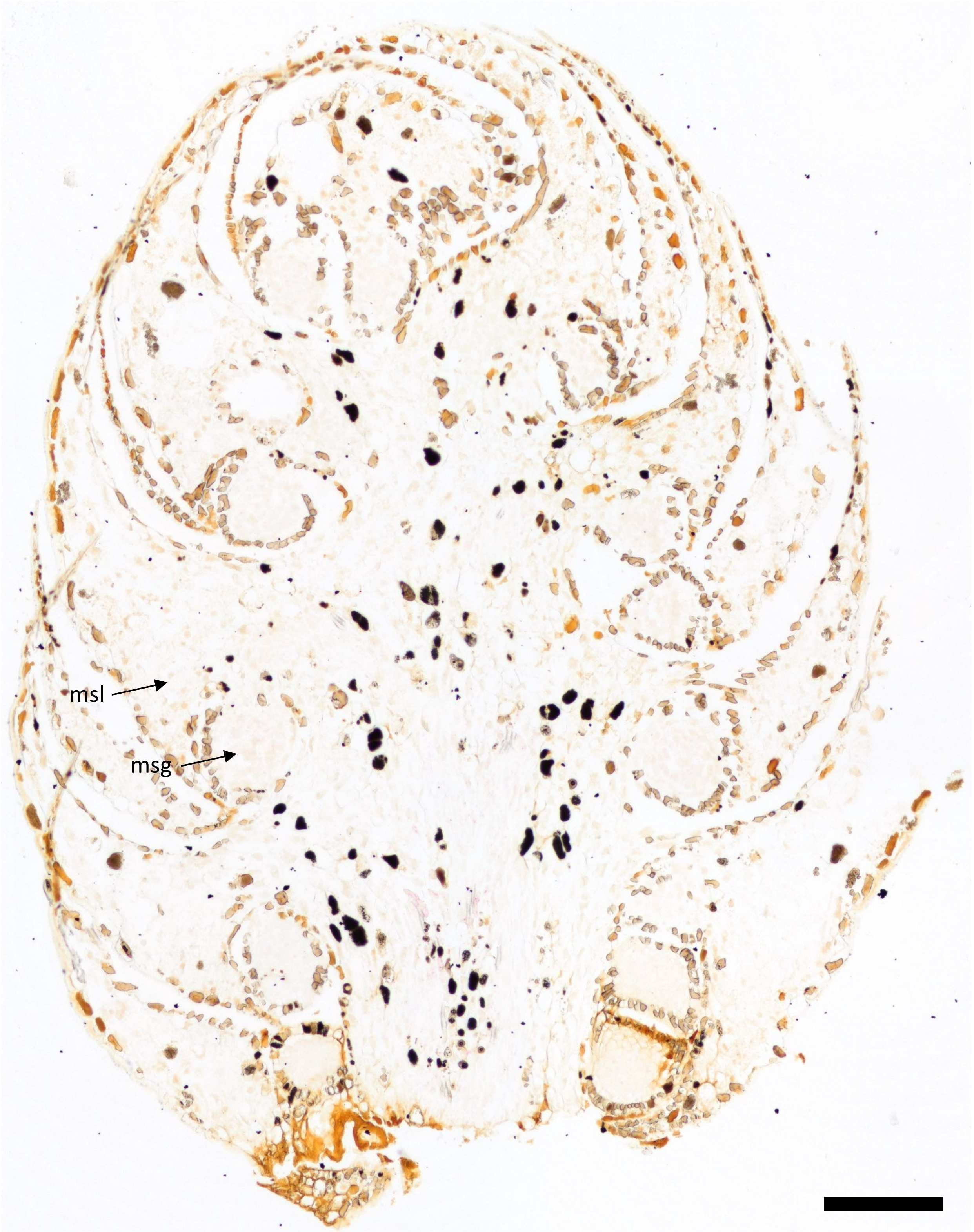


**Fig. S29.** *AG2* expression in *Juniperus virginiana* pollen cone. Note that some expression was detected in vasculature at the base of the cone. No signal was detected in the microsporangia or microsporophylls, however. msl = microsporophyll; msg = microsporangium. Scale bar = 200 µm.

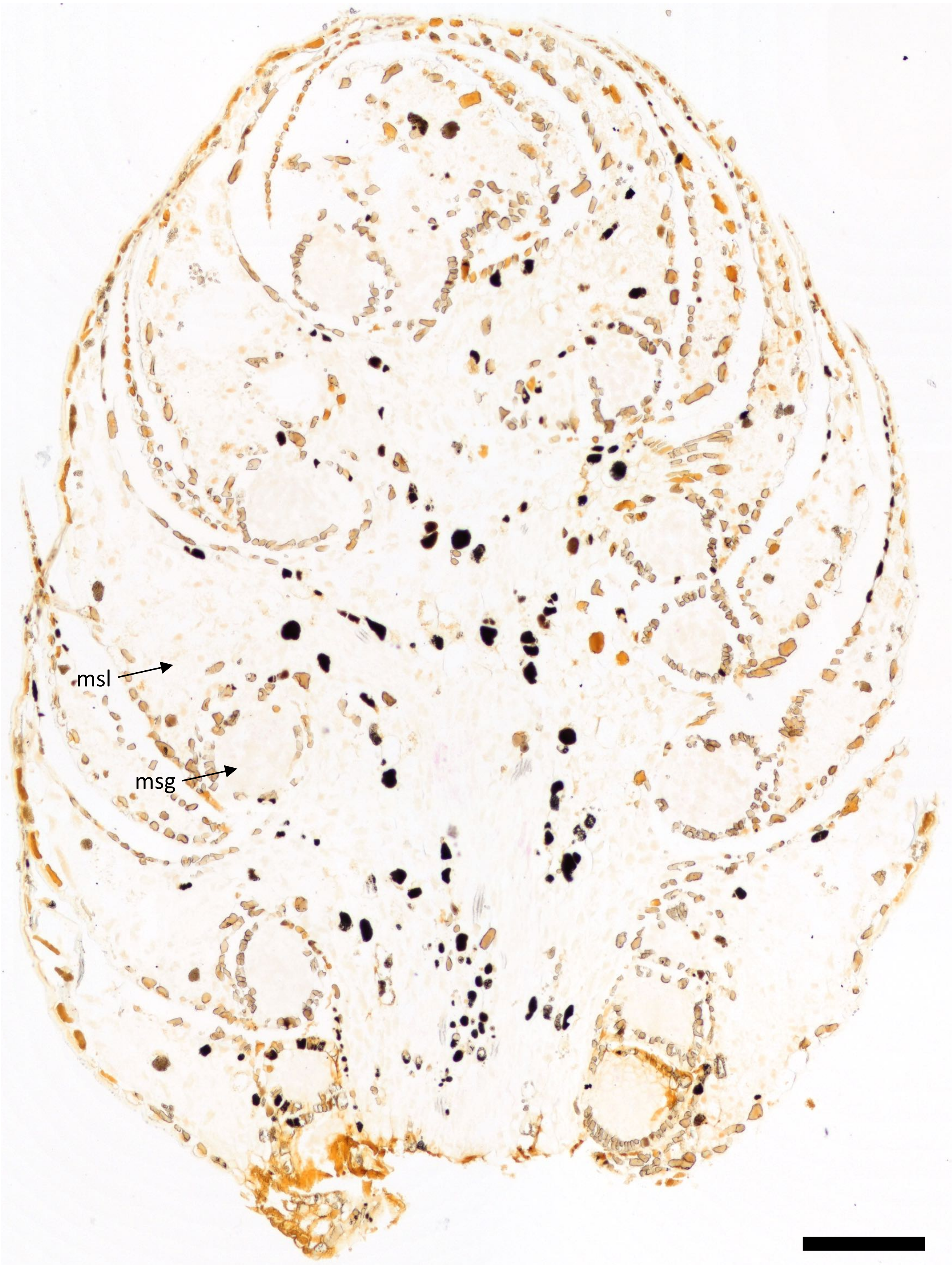


**Fig. S30.** *AG3* expression in *Juniperus virginiana* pollen cone. Note that some expression was detected in vasculature at the base of the cone. No signal was detected in the microsporangia or microsporophylls, however. msl = microsporophyll; msg = microsporangium. Scale bar = 200 µm.

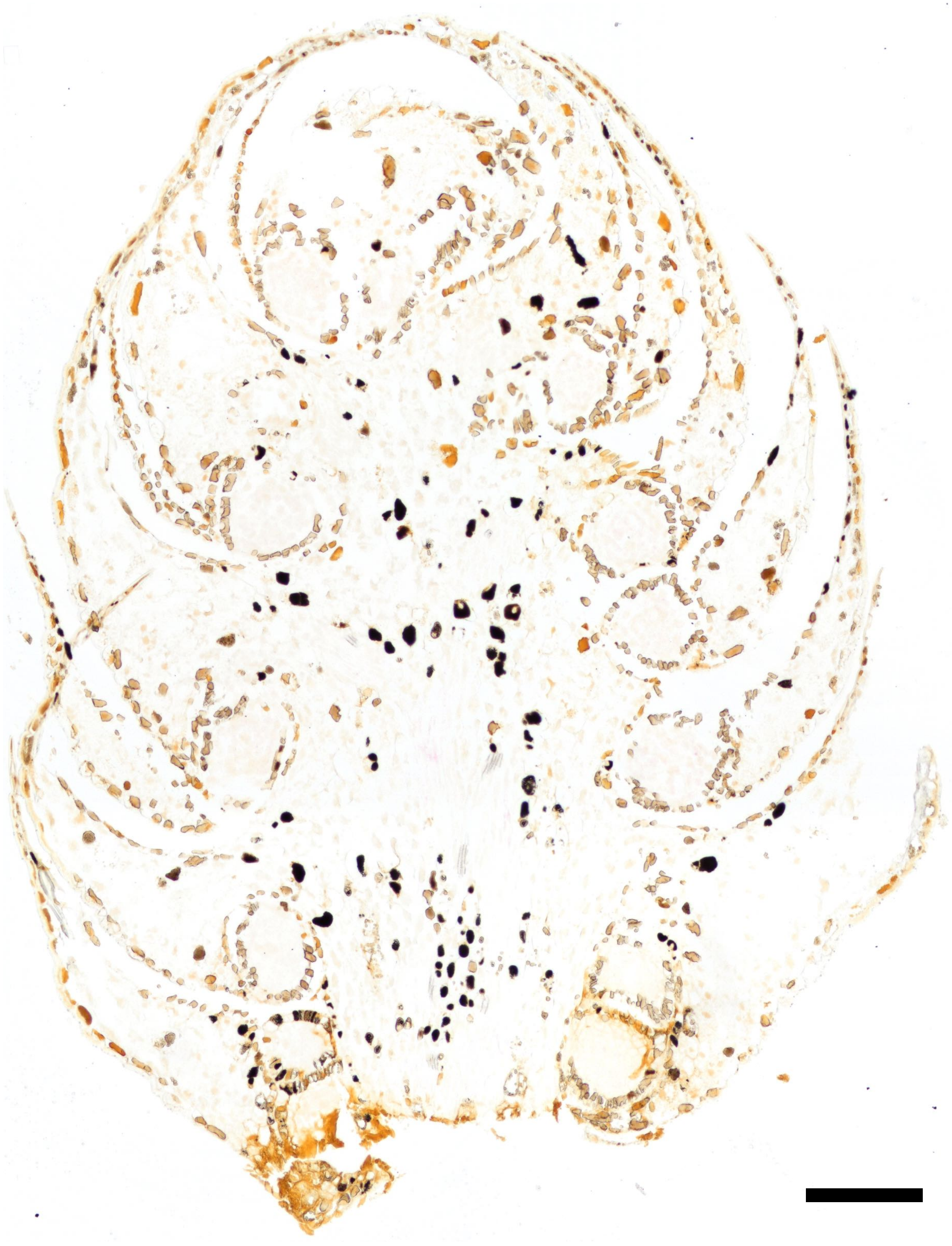

**Fig. S31.** Negative control for *Juniperus virginiana* pollen cone. Scale bar = 200 µm.

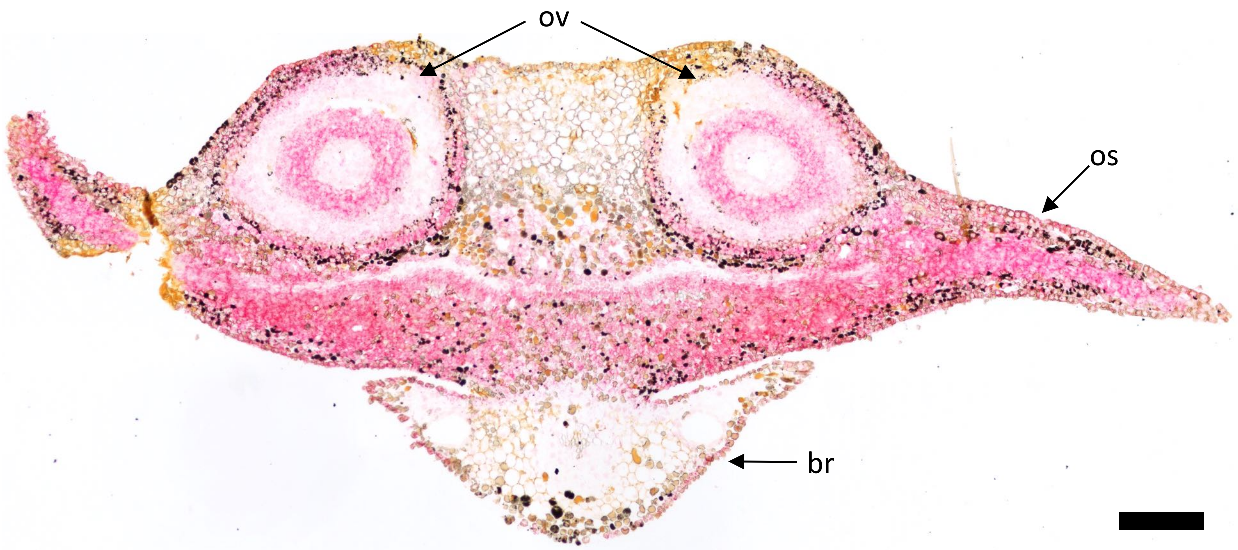


**Fig. S32.** *DAL2* expression in transverse section through bract (br), ovuliferous scale (os), and ovules (ov) of *Picea abies* (Pinaceae). Scale bar = 200 μm.

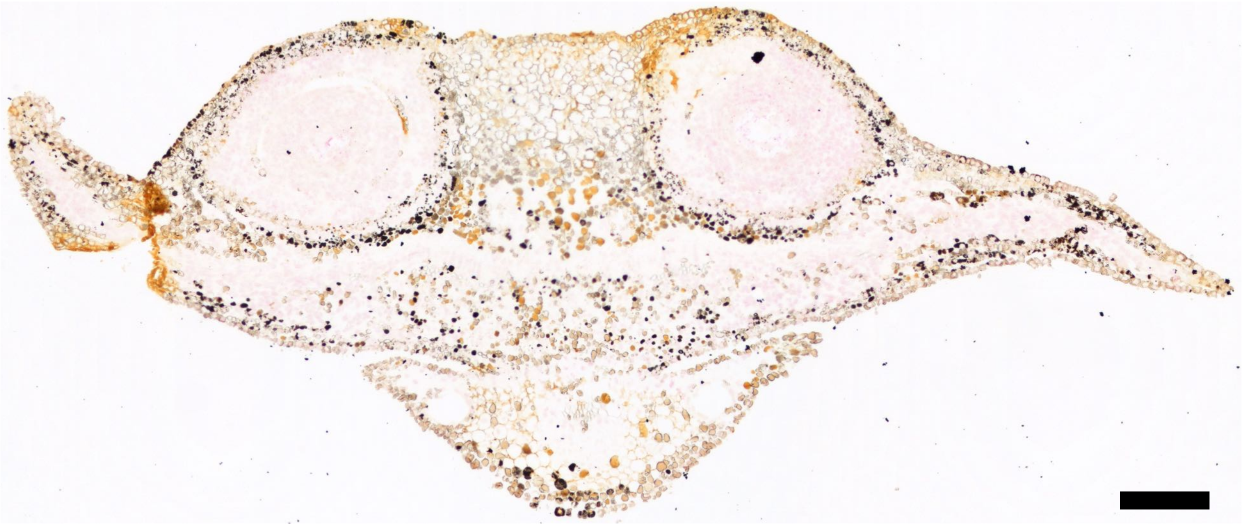

**Fig. S33.** Negative control for transverse section through bract, ovuliferous scale, and ovules of *Picea abies*. Scale bar = 200 µm.

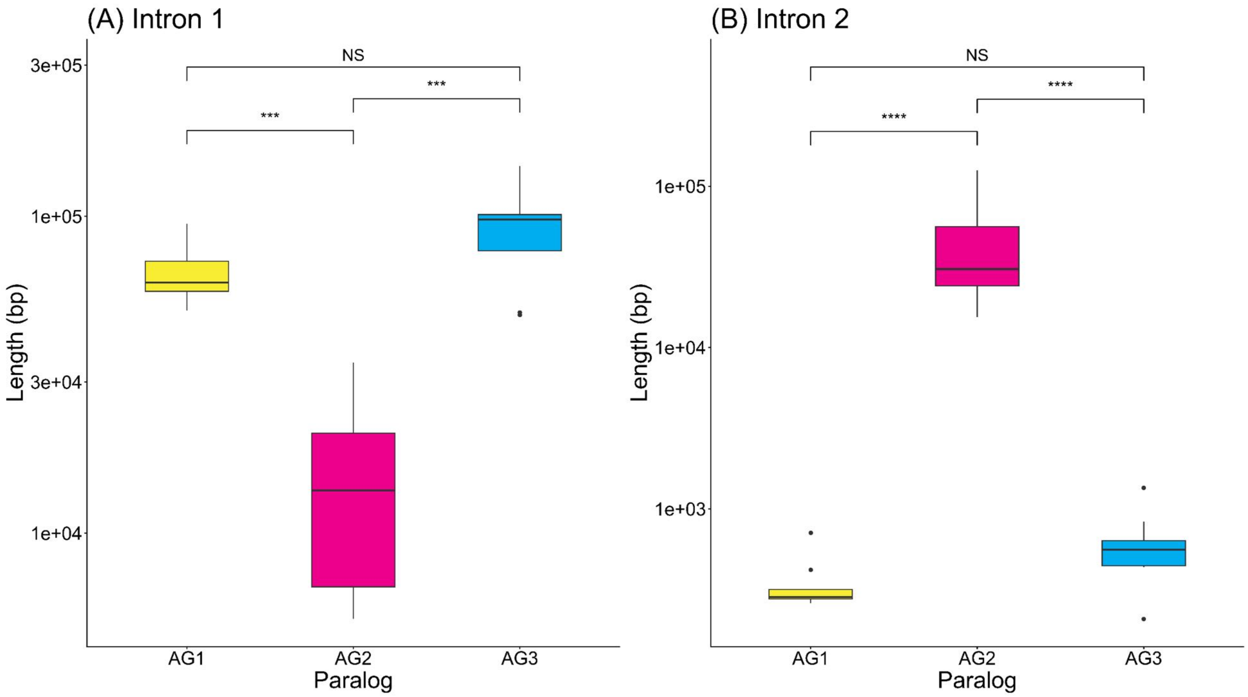


**Fig. S34.** Comparison of lengths for **(A)** first (Welch's ANOVA on log10-transformed data: F = 21.459, *P* = 0.0001119) and **(B)** second (Welch's ANOVA on log10-transformed data: F = 132.27; *P* = 2.717 x $10^{-9}$) introns across *AG* paralogs in 8 Cupressaceae species (n = 8 for each paralog). For Games-Howell post-hoc pairwise tests: *** = 0.0001 < *P* < 0.001; **** = 0 < *P* < 0.0001; NS = not significant (*P* > 0.05). Figure was made in R (19) with the ggplot2 (v. 4.0.3) (20), ggsignif (v. 0.6.4) (21), and gridExtra (v. 2.3.1) (22) packages.

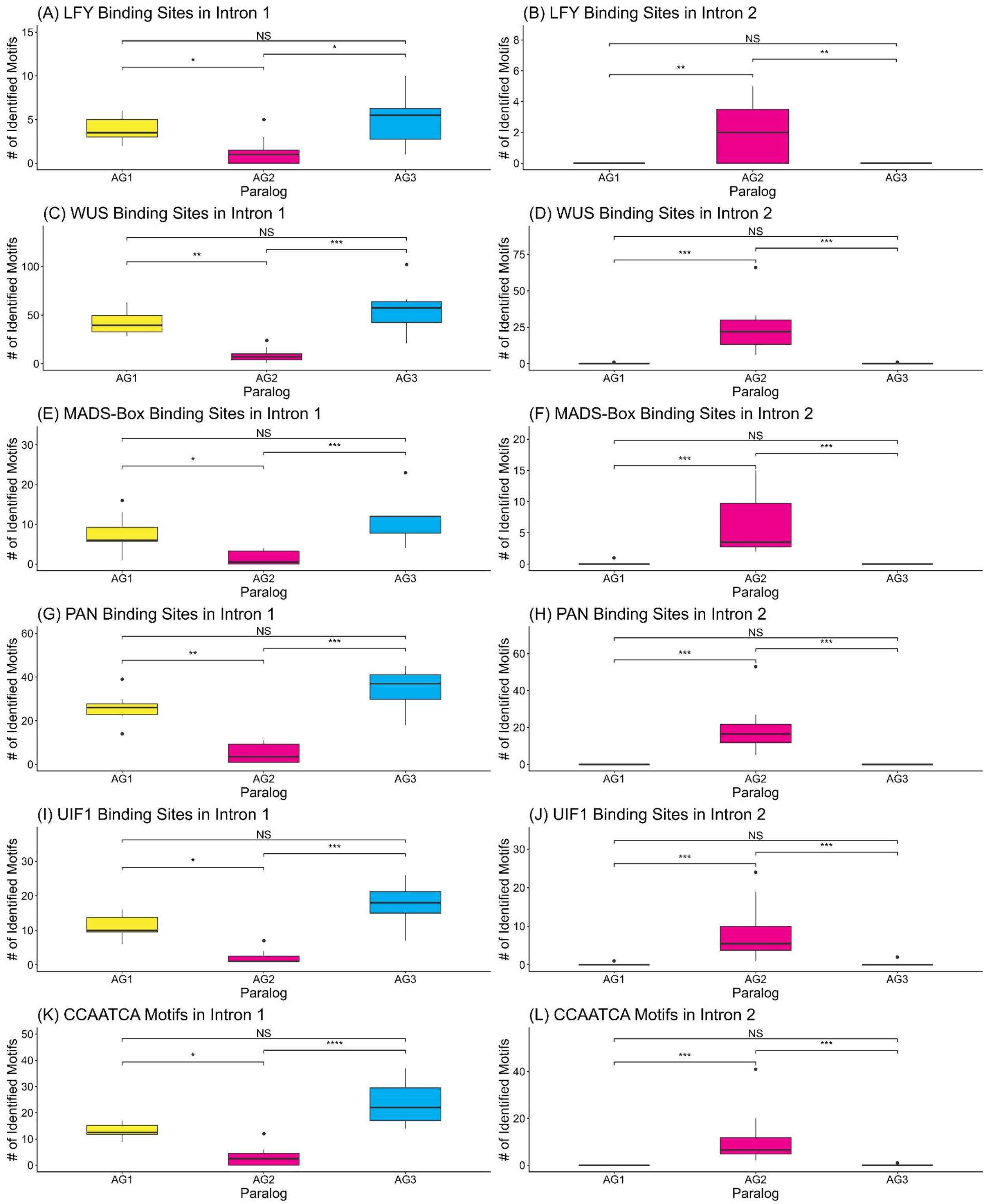


**Fig. S35.** Comparison of number of identified transcription factor binding sites within the introns across *AG* paralogs for 8 Cupressaceae species (n = 8 for each paralog). **(A-B)** Number of LEAFY-binding sites in **(A)** first (Kruskal-Wallis: $\chi^2 = 9.1972$, $P = 0.01007$) and **(B)** second (Kruskal-Wallis: $\chi^2 = 11.917$, $P = 0.002584$) introns. **(C-D)** Number of WUSCHEL-binding sites in **(C)** first (Kruskal-Wallis: $\chi^2 = 15.482$, $P = 0.0004346$) and **(D)** second (Kruskal-Wallis: $\chi^2 = 19.169$, $P = 6.88 \times 10^{-5}$) introns. **(E-F)** Number of binding sites for MADS-box proteins in **(E)** first (Kruskal-Wallis: $\chi^2 = 14.275$, $P = 0.0007948$) and **(F)** second

(Kruskal-Wallis: $\chi^2 = 20.433$, $P = 3.657 \times 10^{-5}$) introns. **(G-H)** Number of PERIANTHIA-binding sites in **(G)** first (Kruskal-Wallis: $\chi^2 = 16.727$, $P = 0.0002332$) and **(H)** second (Kruskal-Wallis: $\chi^2 = 21.834$, $P = 1.814 \times 10^{-5}$) introns. **(I-J)** Number of binding sites for ULTRAPETALA1 INTERACTING FACTOR 1 in **(I)** first (Kruskal-Wallis: $\chi^2 = 17.122$, $P = 0.0001914$) and **(J)** second (Kruskal-Wallis: $\chi^2 = 18.294$, $P = 0.0001065$) introns. **(K-L)** Number of CCAATCA motifs in **(K)** first (Kruskal-Wallis: $\chi^2 = 17.805$, $P = 0.0001361$) and **(L)** second (Kruskal-Wallis: $\chi^2 = 20.409$, $P = 3.7 \times 10^{-5}$) introns. For Dunn's post-hoc pairwise tests: * = $0.01 < P < 0.05$; ** = $0.001 < P < 0.01$; *** = $0.0001 < P < 0.001$; **** = $0 < P < 0.0001$; NS = not significant ($P > 0.05$). Figure was made in R (19) with the ggplot2 (20), ggsignif (21), and gridExtra (22) packages.

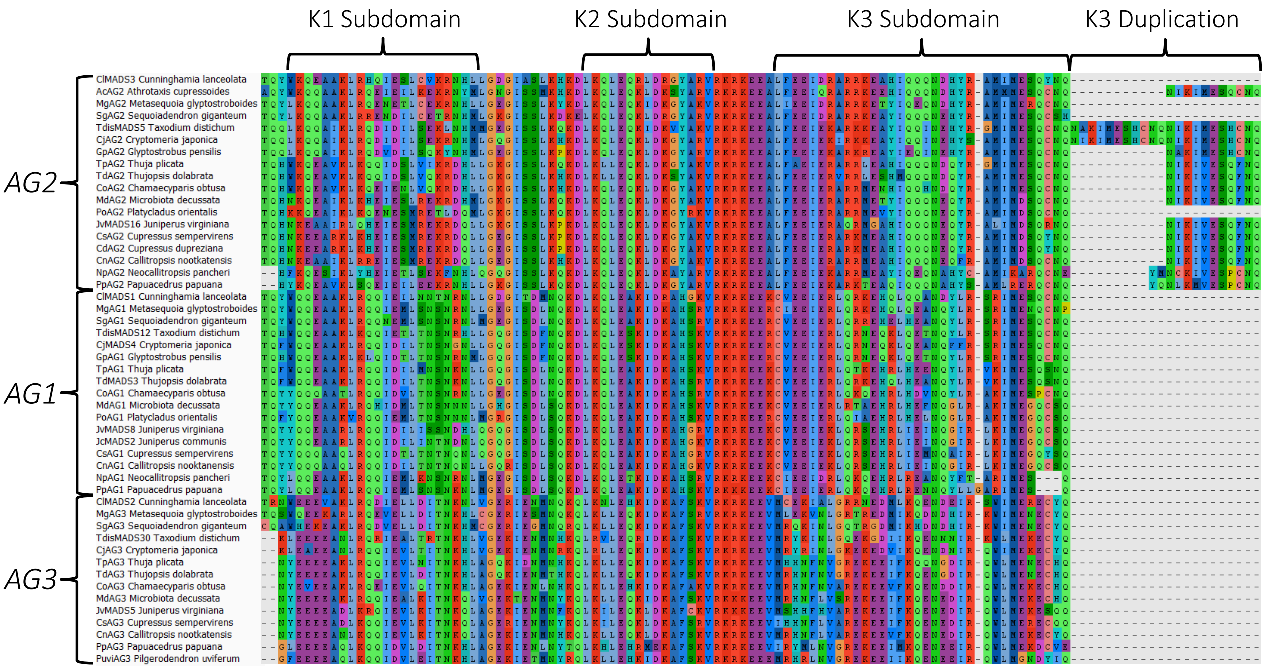


**Fig. S36.** Amino acid alignment of K domain of Cupressaceae *AG* paralogs showing duplication of K3 subdomain in *AG2* genes.

**Tables**

**Table S1.** Genomes and transcriptomes used in this study. * indicate species included in intron length and transcription factor binding site analyses.

| Species | Accession | Literature Cited |
|---|---|---|
| *Larix kaempferi* | GenBank Assembly: GCA_027924585.1 | (23) |
| *Pinus albicaulis* | GenBank Assembly: GCA_034641835.3 | (24) |
| *Pinus tabuliformis* | GenBank Assembly: GCA_031772625.1 | (25) |
| *Pinus taeda* | GenBank Assembly: GCA_000404065.3 | (26) |
| *Pseudotsuga menziesii* | GenBank Assembly: GCA_001517045.1 | (27) |
| *Nageia nagi* | GenBank Assembly: GCA_050084985.1 | (28) |
| *Wollemia nobilis* | GenBank Assembly: GCA_028583015.1 | (29) |
| *Cunninghamia lanceolata** | GenBank Assembly: GCA_027924645.1 | (23) |
| *Sequoiadendron giganteum** | GenBank Assembly: GCA_007115665.2 | (30) |
| *Metasequoia glyptostroboides** | NGDC Accession: GWHCBJF00000000 | (31) |
| *Taxodium distichum** | NGDC Accession: GWHFQIU00000000.1 | (32) |
| *Cryptomeria japonica** | GenBank Assembly: GCA_030272615.1 | (33) |
| *Thuja plicata** | GenBank Assembly: GCA_018584345.1 | (34) |
| *Chamaecyparis obtuse** | GenBank Assembly: GCA_027943335.1 | (23) |
| *Cupressus sempervirens** | GenBank Assembly: GCA_028749045.1 | (35) |
| *Torreya grandis* | GenBank Assembly: GCA_037957365.1 | (36) |
| *Taxus chinensis* | GenBank Assembly: GCA_019776745.2 | (37) |
| *Taxus wallichiana* | GenBank Assembly: GCA_018340775.1 | (38) |
| *Cycas panzhihuaensis* | GenBank Assembly: GCA_023213395.1 | (39) |
| *Welwitschia mirabilis* | CNGB Assembly ID: CNA0022760 | (40) |
| One Thousand Plant Transcriptomes (OneKP) | | (41) |
| *Agathis macrophylla* | GenBank: GKZN00000000.1 | (17) |
| *Callitropsis nootkatensis* | GenBank: GKZO00000000.1 | (17) |
| *Cephalotaxus sinensis* | GenBank: GKZP00000000.1 | (17) |
| *Sciadopitys verticillata* | GenBank: GKZY00000000.1 | (17) |
| *Taxus baccata* | GenBank: GLAA00000000.1 | (17) |
| *Tsuga canadensis* | GenBank: GLAB00000000.1 | (17) |

**Table S2.** Assembly statistics for *Taxodium distichum* and *Juniperus virginiana* transcriptomes.

| | | *Taxodium distichum* | *Juniperus virginiana* |
|---|---|---|---|
| **E90N50** | | 2222 | 1993 |
| **% of Reads Aligned** | First Replicate | 95.3% | 93.7% |
| | Second Replicate | 95.9% | 94.3% |
| | Third Replicate | 95.7% | 94.0% |
| **BUSCO** | Complete and single-copy | 1448 (90.3%) | 1460 (91.1%) |
| | Complete and duplicated | 28 (1.7%) | 35 (2.2%) |
| | Fragmented | 13 (0.8%) | 15 (0.9%) |
| | Missing | 114 (7.2%) | 93 (5.8%) |

**Table S3.** Genes and their accessions included in the gene tree.

| Gene Name | Other Name | Species | GenBank Accession Number / Transcript ID / Location in Genome |
|---|---|---|---|
| *DAL1* | | *Picea abies* | X80902.1 |
| *DAL2* | | *Picea abies* | X79280.1 |
| *DAL3* | | *Picea abies* | X79281.1 |
| *DAL4* | | *Picea abies* | KC347013.1 |
| *DAL5* | | *Picea abies* | KC347011.1 |
| *DAL6* | | *Picea abies* | KC347010.1 |
| *DAL9* | | *Picea abies* | KC347014.1 |
| *DAL10* | | *Picea abies* | AF064080.1 |
| *DAL11* | | *Picea abies* | AF158539.1 |
| *DAL12* | | *Picea abies* | AF158542.1 |
| *DAL13* | | *Picea abies* | AY269278.1 |
| *DAL14* | | *Picea abies* | KC347012.1 |
| *DAL18*[g] | | *Picea abies* | KC516777.1 |
| *DAL19* | | *Picea abies* | KC347015.1 |
| *DAL20*[g] | | *Picea abies* | KC305492.1 |
| *DAL21* | | *Picea abies* | KC347016.1 |
| *DAL22* | | *Picea abies* | KC347017.1 |
| *DAL23* | | *Picea abies* | KC347018.1 |
| *PrDGL* | | *Pinus radiata* | AF120097.1 |
| *PrMADS* | | *Pinus radiata* | AF023615.1 |
| *PrMADS2* | | *Pinus radiata* | U42400.1 |
| *PrMADS3* | | *Pinus radiata* | U76726.1 |
| *PrMADS4* | | *Pinus radiata* | U90345.1 |
| *PrMADS5* | | *Pinus radiata* | U90346.1 |
| *PrMADS6* | | *Pinus radiata* | U90347.1 |
| *PrMADS7* | | *Pinus radiata* | U90348.1 |
| *PrMADS8* | | *Pinus radiata* | U90349.1 |
| *PrMADS9* | | *Pinus radiata* | U90344.1 |
| *AG* | *TbAG3* | *Taxus baccata* | JF519754.1 / GLAA01017259.1 |
| *AGL6* | | *Taxus baccata* | JF519755.1 |
| *TgAG* | | *Taxus globosa* | HM596863.1 |
| *PodAG* | | *Podocarpus matudae* | HM596865.1 / GKZW01002102.1 |
| *JcMADS2* | | *Juniperus communis* | HM177455.1 |
| *TdMADS3* | | *Thujopsis dolabrata* | HM177454.1 |
| *CjMADS4* | *CjAG1* | *Cryptomeria japonica* | HM177453.1 / NC_081407.1 Chromosome 3: 703399963-703463969 (+ strand) |
| *CjAGL6* | | *Cryptomeria japonica* | AB359029.2 |
| *CjB1* | | *Cryptomeria japonica* | AB359036.1 |
| *CjB2* | | *Cryptomeria japonica* | AB359033.1 |
| *CjB4* | | *Cryptomeria japonica* | AB359034.1 |
| *CjB5* | | *Cryptomeria japonica* | AB359035.1 |
| *CjB6* | | *Cryptomeria japonica* | AB359032.1 |
| *CjSOC1* | | *Cryptomeria japonica* | AB359030.1 |
| *TM8-l1* | | *Cryptomeria japonica* | AB359027.1 |
| *TM8-l2* | | *Cryptomeria japonica* | AB359028.1 |
| *GBM5* | | *Ginkgo biloba* | AY114304.1 |
| *GbMADS1* | | *Ginkgo biloba* | AB029463.1 |
| *GbMADS3* | | *Ginkgo biloba* | AB029465.1 |

| | | | |
|---|---|---|---|
| *GbMADS4* | | *Ginkgo biloba* | AB029466.1 |
| *GbMADS5* | | *Ginkgo biloba* | AB029467.1 |
| *GbMADS6* | | *Ginkgo biloba* | AB029468.1 |
| *GbMADS7* | | *Ginkgo biloba* | AB029469.1 |
| *GbMADS8* | | *Ginkgo biloba* | AB029470.1 |
| *GbMADS9* | | *Ginkgo biloba* | AB029471.1 |
| *GbMADS10* | | *Ginkgo biloba* | AB029472.1 |
| *GbMADS11* | | *Ginkgo biloba* | AB029473.1 |
| *GGM1* | | *Gnetum gnemon* | AJ132207.2 |
| *GGM2* | | *Gnetum gnemon* | AJ132208.1 |
| *GGM3* | | *Gnetum gnemon* | AJ132209.1 |
| *GGM4* | | *Gnetum gnemon* | AJ132210.1 |
| *GGM5* | | *Gnetum gnemon* | AJ132211.1 |
| *GGM6* | | *Gnetum gnemon* | AJ132212.2 |
| *GGM7* | | *Gnetum gnemon* | AJ132213.1 |
| *GGM8* | | *Gnetum gnemon* | AJ132214.1 |
| *GGM9* | | *Gnetum gnemon* | AJ132215.1 |
| *GGM10* | | *Gnetum gnemon* | AJ132216.2 |
| *GGM11* | | *Gnetum gnemon* | AJ132217.2 |
| *GGM12* | | *Gnetum gnemon* | AJ132218.1 |
| *GGM13* | | *Gnetum gnemon* | AJ132219.1 |
| *GGM14* | | *Gnetum gnemon* | AJ251554.1 |
| *GGM15* | | *Gnetum gnemon* | AJ251555.1 |
| *GGM16* | | *Gnetum gnemon* | AJ251556.1 |
| *GGM17* | | *Gnetum gnemon* | AJ251557.1 |
| *GGM18* | | *Gnetum gnemon* | AJ251559.1 |
| *GGM19* | | *Gnetum gnemon* | AJ251558.1 |
| *GpMADS4* | | *Gnetum parvifolium* | AB022666.1 |
| *WelAP3/PI-1* | | *Welwitschia mirabilis* | KF145184.1 |
| *WelAP3/PI-2* | | *Welwitschia mirabilis* | KF145185.1 |
| *WelAG* | | *Welwitschia mirabilis* | KF145186.1 / Chromosome 4: 51935650-51962367 |
| *WelBsister* | | *Welwitschia mirabilis* | KF145187.1 |
| *WelAGL6* | | *Welwitschia mirabilis* | KF145188.1 |
| *CyAG* | | *Cycas edentata* | AF492455.1 |
| *StaAG* | | *Stangeria eriopus* | HM596864.1 |
| *MacAG* | | *Macrozamia spiralis* | HM596862.1 |
| *CerAG* | | *Ceratozamia mexicana* | HM596861.1 |
| *Il.fl.AG* | | *Illicium floridianum* | AY936229.1 |
| *Il.fl.PI* | | *Illicium floridianum* | AY936224.1 |
| *Il.fl.AP3.1* | | *Illicium floridianum* | AY936225.1 |
| *Il.fl.AP3.2* | | *Illicium floridianum* | AY936226.1 |
| *Il.fl.AP3.3* | | *Illicium floridianum* | AY936227.1 |
| *Am.tr.AG* | | *Amborella trichopoda* | AY936231.1 |
| *Am.tr.AGL2* | | *Amborella trichopoda* | AY936232.1 |
| *Am.tr.AGL6* | | *Amborella trichopoda* | AY936234.1 |
| *Am.tr.PI* | | *Amborella trichopoda* | AY337760.1 |
| *Am.tr.AP3-1* | | *Amborella trichopoda* | AY337743.1 |
| *Am.tr.AP3-2* | | *Amborella trichopoda* | AY337744.1 |
| *Nu.ad.AP1* | | *Nuphar advena* | AY936223.1 |
| *Nu.ad.AG* | | *Nuphar advena* | AY936230.1 |
| *Nuad.AGL6* | | *Nuphar advena* | GU048649.1 |
| *Ma.gr.AG* | | *Magnolia grandiflora* | AY936228.1 |
| *Ma.gr.AGL6* | | *Magnolia grandiflora* | AY936233.1 |

| | | | |
|---|---|---|---|
| *Ma.gr.AP1* | | *Magnolia grandiflora* | AY821777.1 |
| *Ma.gr.PI* | | *Magnolia grandiflora* | AY821779.1 |
| *Ma.gr.AGL9* | | *Magnolia grandiflora* | AY821782.1 |
| *Ma.gr.AGL2* | | *Magnolia grandiflora* | AY821781.1 |
| *SOC1* | | *Magnolia virginiana* | GQ489241.1 |
| *Eu.be.AP1* | | *Eupomatia bennettii* | AY821776.1 |
| *Eu.be.AP3-1* | | *Eupomatia bennettii* | AY337750.1 |
| *Eu.be.AP3-2* | | *Eupomatia bennettii* | AY337751.1 |
| *Eu.be.PI* | | *Eupomatia bennettii* | AY337740.1 |
| *Eu.be.AGL9* | | *Eupomatia bennettii* | AY821780.1 |
| *LtAGL9* | | *Liriodendron tulipifera* | AY850182.1 |
| *MfAGL6A* | | *Michelia figo* | AY306157.1 |
| *MfAGL6B* | | *Michelia figo* | AY306158.1 |
| *MfFL* | | *Michelia figo* | AY306159.1 |
| *MfPI-1* | | *Michelia figo* | AF052863.1 |
| *MfAP3* | | *Michelia figo* | AF052877.1 |
| *CcAP3* | | *Cabomba caroliniana* | AB158353.1 |
| *CcPI* | | *Cabomba caroliniana* | AB158354.1 |
| *Caca.AGL6-1* | | *Cabomba caroliniana* | GU048631.1 |
| *Caca.AP1* | | *Cabomba caroliniana* | GU048634.1 |
| *Caca.AG* | | *Cabomba caroliniana* | GU048626.1 |
| *Caca.Bs1* | | *Cabomba caroliniana* | GU048636.1 |
| *Caca.SEP1-1* | | *Cabomba caroliniana* | GU048628.1 |
| *Nenu.AG* | | *Nelumbo nucifera* | GU048639.1 |
| *Nenu.AGL6* | | *Nelumbo nucifera* | GU048640.1 |
| *Nenu.SEP1* | | *Nelumbo nucifera* | GU048641.1 |
| *Nenu.SEP3* | | *Nelumbo nucifera* | GU048642.1 |
| *Nenu.AP1* | | *Nelumbo nucifera* | GU048643.1 |
| *Nenu.AP3-1* | | *Nelumbo nucifera* | GU048644.1 |
| *Nenu.AP3-2* | | *Nelumbo nucifera* | GU048645.1 |
| *Nenu.PI* | | *Nelumbo nucifera* | GU048646.1 |
| *OsMADS3* | | *Oryza sativa* | NM_001424572.1 |
| *OsMADS13* | | *Oryza sativa* | NM_001404655.1 |
| *OsMADS26* | | *Oryza sativa* | NM_001403334.1 |
| *OsMADS15* | | *Oryza sativa* | NM_001402986.1 |
| *OsMADS8* | | *Oryza sativa* | NM_001422804.1 |
| *OsMADS6* | | *Oryza sativa* | NM_001401921.1 |
| *OsMADS58* | | *Oryza sativa* | NM_001420259.1 |
| *OsMADS21* | | *Oryza sativa* | NM_001405446.1 |
| *ZZM2* | *MADS2* | *Zea mays* | NM_001111456.2 |
| *ZAG1* | | *Zea mays* | NM_001111851.2 |
| *ZAG2* | | *Zea mays* | NM_001305994.1 |
| *JOINTLESS* | | *Solanum lycopersicum* | NM_001319841.1 |
| *TM6* | *TDR6* | *Solanum lycopersicum* | NM_001324380.1 |
| *TM8* | *TDR8* | *Solanum lycopersicum* | NM_001247176.2 |
| *TAG1* | | *Solanum lycopersicum* | NM_001308296.2 |
| *TAGL1* | | *Solanum lycopersicum* | NM_001313930.1 |
| *TAGL11* | | *Solanum lycopersicum* | NM_001247265.2 |
| *pMADS3* | | *Petunia* x *hybrida* | X72912.1 |
| *FBP6* | | *Petunia* x *hybrida* | X68675.1 |
| *FBP7* | | *Petunia* x *hybrida* | X81651.1 |
| *FBP11* | | *Petunia* x *hybrida* | X81852.1 |
| *SQUA* | | *Antirrhinum majus* | X63701.1 |

| | | | |
|---|---|---|---|
| *DEF* | | *Antirrhinum majus* | X52023.1 |
| *GLO* | | *Antirrhinum majus* | X68831.1 |
| *PLE* | | *Antirrhinum majus* | AB516404.1 |
| *FAR* | | *Antirrhinum majus* | AB516405.1 |
| *SEP3* | *AGL9* | *Arabidopsis thaliana* | NM_102272.4 |
| *CAL* | *AGL10, CAL1* | *Arabidopsis thaliana* | NM_102395.3 |
| *AP1* | *AGL7* | *Arabidopsis thaliana* | NM_105581.3 |
| *AGL12* | *XAL1* | *Arabidopsis thaliana* | NM_105825.3 |
| *MAF1* | *FLM, AGL27* | *Arabidopsis thaliana* | NM_001334764.1 |
| *SEP4* | *AGL3* | *Arabidopsis thaliana* | NM_126418.2 |
| *ANR1* | *AGL44* | *Arabidopsis thaliana* | NM_001335416.1 |
| *SVP* | *AGL22, FAQ1* | *Arabidopsis thaliana* | NM_127820.4 |
| *AGL17* | | *Arabidopsis thaliana* | NM_127828.3 |
| *SHP1* | *AGL1* | *Arabidopsis thaliana* | NM_001339937.1 |
| *SHP2* | *AGL5* | *Arabidopsis thaliana* | NM_180046.2 |
| *AGL6* | *RSB1* | *Arabidopsis thaliana* | NM_130127.2 |
| *SOC1* | *AGL20* | *Arabidopsis thaliana* | NM_130128.4 |
| *SEP2* | *AGL4* | *Arabidopsis thaliana* | NM_111098.4 |
| *AP3* | | *Arabidopsis thaliana* | NM_115294.6 |
| *PI* | | *Arabidopsis thaliana* | NM_122031.4 |
| *AGL16* | | *Arabidopsis thaliana* | NM_115583.6 |
| *AGL18* | | *Arabidopsis thaliana* | NM_115599.4 |
| *AGL13* | | *Arabidopsis thaliana* | NM_115976.2 |
| *STK* | *AGL11* | *Arabidopsis thaliana* | NM_001340645.1 |
| *AGL14* | *XAL2* | *Arabidopsis thaliana* | NM_001340739.1 |
| *AG* | | *Arabidopsis thaliana* | NM_118013.2 |
| *AGL19* | *GL19* | *Arabidopsis thaliana* | NM_001341568.1 |
| *AGL24* | | *Arabidopsis thaliana* | NM_118587.6 |
| *AGL21* | | *Arabidopsis thaliana* | NM_119955.3 |
| *FLC* | *FLF, AGL25* | *Arabidopsis thaliana* | NM_121052.3 |
| *AGL15* | | *Arabidopsis thaliana* | NM_001343293.1 |
| *SEP1* | *AGL2* | *Arabidopsis thaliana* | NM_001125758.2 |
| *TT16* | *ABS, AGL32* | *Arabidopsis thaliana* | NM_001343797.1 |
| *AGL72* | | *Arabidopsis thaliana* | NM_124565.2 |
| *AGL71* | | *Arabidopsis thaliana* | NM_001203588.1 |
| *FUL* | *AGL8* | *Arabidopsis thaliana* | NM_125484.4 |
| *AGL42* | *FYF* | *Arabidopsis thaliana* | NM_180916.1 |
| *AGL31* | *MAF2* | *Arabidopsis thaliana* | NM_001126026.2 |
| *MAF3* | *FCL3, AGL70* | *Arabidopsis thaliana* | NM_125905.4 |
| *MAF4* | | *Arabidopsis thaliana* | NM_001203688.1 |
| *MAF5* | *AGL68* | *Arabidopsis thaliana* | NM_001085330.2 |
| *DIA*[b] | *AGL61* | *Arabidopsis thaliana* | NM_179727.5 |
| *AGL62*[b] | | *Arabidopsis thaliana* | NM_125437.5 |
| *AGL23*[b] | | *Arabidopsis thaliana* | NM_105210.2 |
| *ClMADS1* | *ClAG1* | *Cunninghamia lanceolata* | MT103468.1 / BSBN01001244.1<br>CLA1p1244: 599083-660841 (+<br>strand) |
| *ClMADS2* | *ClAG3* | *Cunninghamia lanceolata* | MT103469.1 / BSBN01000580.1<br>CLA1p0580: 5974258-6024870 (+<br>strand) |
| *ClMADS3* | *ClAG2* | *Cunninghamia lanceolata* | MT103470.1 / BSBN01000580.1<br>CLA1p0580: 1492892-1647222 (+<br>strand) |
| *ClMADS4*[g] | | *Cunninghamia lanceolata* | MT103471.1 |

| | | | |
|---|---|---|---|
| *ClMADS5*[g] | | *Cunninghamia lanceolata* | MT103472.1 |
| *ClMADS6*[g] | | *Cunninghamia lanceolata* | MT103473.1 |
| *ClMADS7* | | *Cunninghamia lanceolata* | MT103474.1 |
| *ClMADS8* | | *Cunninghamia lanceolata* | MT103475.1 |
| *ClMADS9* | | *Cunninghamia lanceolata* | MT103476.1 |
| *ClMADS10* | | *Cunninghamia lanceolata* | MT103477.1 |
| *ClMADS11* | | *Cunninghamia lanceolata* | MT103478.1 |
| *ClMADS12* | | *Cunninghamia lanceolata* | MT103479.1 |
| *ClMADS13* | | *Cunninghamia lanceolata* | MT103480.1 |
| *ClMADS14* | | *Cunninghamia lanceolata* | MT103481.1 |
| *ClMADS15* | | *Cunninghamia lanceolata* | MT103482.1 |
| *ClMADS16* | | *Cunninghamia lanceolata* | MT103483.1 |
| *ClMADS17* | | *Cunninghamia lanceolata* | MT103484.1 |
| *ClMADS18* | | *Cunninghamia lanceolata* | MT103485.1 |
| *ClMADS19* | | *Cunninghamia lanceolata* | MT103486.1 |
| *ClMADS20* | | *Cunninghamia lanceolata* | MT103487.1 |
| *ClMADS21* | | *Cunninghamia lanceolata* | MT103488.1 |
| *ClMADS22* | | *Cunninghamia lanceolata* | MT103489.1 |
| *ClMADS23* | | *Cunninghamia lanceolata* | MT103490.1 |
| *ClMADS24* | | *Cunninghamia lanceolata* | MT103491.1 |
| *ClMADS25* | | *Cunninghamia lanceolata* | MT103492.1 |
| *ClMADS26* | | *Cunninghamia lanceolata* | MT103493.1 |
| *ClMADS27* | | *Cunninghamia lanceolata* | MT103494.1 |
| *ClMADS28* | | *Cunninghamia lanceolata* | MT103495.1 |
| *ClMADS29* | | *Cunninghamia lanceolata* | MT103496.1 |
| *ClMADS30* | | *Cunninghamia lanceolata* | MT103497.1 |
| *ClMADS31* | | *Cunninghamia lanceolata* | MT103498.1 |
| *ClMADS32* | | *Cunninghamia lanceolata* | MT103499.1 |
| *ClMADS33* | | *Cunninghamia lanceolata* | MT103500.1 |
| *TdisMADS1* | | *Taxodium distichum* | TRINITY_DN264_c0_g1_i6 |
| *TdisMADS2* | | *Taxodium distichum* | TRINITY_DN264_c0_g2_i1 |
| *TdisMADS3* | | *Taxodium distichum* | TRINITY_DN264_c0_g3_i1 |
| *TdisMADS4* | | *Taxodium distichum* | TRINITY_DN362_c0_g1_i4 |
| *TdisMADS5* | *TdisAG2* | *Taxodium distichum* | TRINITY_DN404_c0_g1_i2 / GWHFQIU00000011.1 Chromosome 11: 620515498-620553378 (- strand) |
| *TdisMADS6* | | *Taxodium distichum* | TRINITY_DN607_c0_g1_i8 |
| *TdisMADS7* | | *Taxodium distichum* | TRINITY_DN607_c2_g1_i8 |
| *TdisMADS8* | | *Taxodium distichum* | TRINITY_DN607_c2_g4_i1 |
| *TdisMADS9* | | *Taxodium distichum* | TRINITY_DN3983_c0_g1_i14 |
| *TdisMADS10* | | *Taxodium distichum* | TRINITY_DN3983_c0_g2_i7 |
| *TdisMADS11* | | *Taxodium distichum* | TRINITY_DN4775_c0_g1_i3 |
| *TdisMADS12* | *TdisAG1* | *Taxodium distichum* | TRINITY_DN4709_c0_g1_i1 / GWHFQIU00000011.1 Chromosome 11: 622055779-622109664 (+ strand) |
| *TdisMADS13* | | *Taxodium distichum* | TRINITY_DN4981_c0_g1_i15 |
| *TdisMADS14* | | *Taxodium distichum* | TRINITY_DN6906_c0_g1_i1 |
| *TdisMADS15* | | *Taxodium distichum* | TRINITY_DN8513_c1_g1_i1 |
| *TdisMADS16* | | *Taxodium distichum* | TRINITY_DN9486_c0_g1_i3 |
| *TdisMADS17* | | *Taxodium distichum* | TRINITY_DN9689_c0_g1_i2 |
| *TdisMADS18* | | *Taxodium distichum* | TRINITY_DN9703_c0_g1_i4 |
| *TdisMADS19*[a] | | *Taxodium distichum* | TRINITY_DN9703_c0_g2_i1 |

| | | | |
|---|---|---|---|
| *TdisMADS20* | | *Taxodium distichum* | TRINITY_DN10616_c0_g1_i6 |
| *TdisMADS21* | | *Taxodium distichum* | TRINITY_DN11462_c0_g1_i4 |
| *TdisMADS22* | | *Taxodium distichum* | TRINITY_DN13482_c0_g1_i2 |
| *TdisMADS23* | | *Taxodium distichum* | TRINITY_DN13847_c0_g1_i2 |
| *TdisMADS24* | | *Taxodium distichum* | TRINITY_DN14149_c0_g1_i1 |
| *TdisMADS25* | | *Taxodium distichum* | TRINITY_DN14149_c0_g3_i1 |
| *TdisMADS26* | | *Taxodium distichum* | TRINITY_DN16145_c0_g1_i2 |
| *TdisMADS27* | | *Taxodium distichum* | TRINITY_DN16145_c0_g2_i1 |
| *TdisMADS28* | | *Taxodium distichum* | TRINITY_DN18665_c0_g1_i6 |
| *TdisMADS29* | | *Taxodium distichum* | TRINITY_DN18665_c0_g2_i5 |
| *TdisMADS30* | *TdisAG3* | *Taxodium distichum* | TRINITY_DN18980_c0_g2_i19 / GWHFQIU00000011.1 Chromosome 11: 617716484-617815093 (- strand) |
| *TdisMADS31*[g] | | *Taxodium distichum* | TRINITY_DN76969_c0_g1_i1 |
| *TdisMADS32* | | *Taxodium distichum* | TRINITY_DN94077_c0_g1_i1 |
| *TdisMADS33* | | *Taxodium distichum* | TRINITY_DN99330_c0_g1_i1 |
| *JvMADS1* | | *Juniperus virginiana* | TRINITY_DN917_c0_g1_i16 |
| *JvMADS2* | | *Juniperus virginiana* | TRINITY_DN2515_c0_g1_i1 |
| *JvMADS3* | | *Juniperus virginiana* | TRINITY_DN2515_c0_g2_i1 |
| *JvMADS4* | | *Juniperus virginiana* | TRINITY_DN2925_c0_g1_i6 |
| *JvMADS5* | *JvAG3* | *Juniperus virginiana* | TRINITY_DN3832_c0_g1_i9 |
| *JvMADS7* | | *Juniperus virginiana* | TRINITY_DN4183_c0_g1_i1 |
| *JvMADS8* | *JvAG1* | *Juniperus virginiana* | TRINITY_DN5759_c0_g1_i1 |
| *JvMADS9* | | *Juniperus virginiana* | TRINITY_DN6096_c0_g1_i1 |
| *JvMADS10* | | *Juniperus virginiana* | TRINITY_DN6121_c0_g1_i5 |
| *JvMADS11* | | *Juniperus virginiana* | TRINITY_DN8280_c0_g1_i21 |
| *JvMADS12* | | *Juniperus virginiana* | TRINITY_DN8704_c0_g1_i3 |
| *JvMADS14* | | *Juniperus virginiana* | TRINITY_DN10464_c0_g1_i5 |
| *JvMADS15* | | *Juniperus virginiana* | TRINITY_DN10648_c1_g1_i3 |
| *JvMADS16* | *JvAG2* | *Juniperus virginiana* | TRINITY_DN10663_c0_g1_i7 |
| *JvMADS18* | | *Juniperus virginiana* | TRINITY_DN14462_c0_g1_i2 |
| *JvMADS19* | | *Juniperus virginiana* | TRINITY_DN20328_c0_g1_i6 |
| *JvMADS20* | | *Juniperus virginiana* | TRINITY_DN22848_c0_g1_i1 |
| *JvMADS21* | | *Juniperus virginiana* | TRINITY_DN34423_c0_g1_i2 |
| *JvMADS22* | | *Juniperus virginiana* | TRINITY_DN35208_c0_g1_i1 |
| *JvMADS23* | | *Juniperus virginiana* | TRINITY_DN43046_c0_g1_i2 |
| *JvMADS24* | | *Juniperus virginiana* | TRINITY_DN98229_c0_g1_i1 |
| *JvMADS25* | | *Juniperus virginiana* | TRINITY_DN132968_c0_g1_i2 |
| *JvMADS26* | | *Juniperus virginiana* | TRINITY_DN134090_c0_g1_i1 |
| *MgAG1* | | *Metasequoia glyptostroboides* | GWHCBJF00000002 Chromosome 3: 306307916-306404125 (- strand) |
| *MgAG2* | | *Metasequoia glyptostroboides* | GWHCBJF00000002 Chromosome 3: 308588635-308668959 (+ strand) |
| *MgAG3* | | *Metasequoia glyptostroboides* | GWHCBJF00000002 Chromosome 3: 311650618-311796166 (+ strand) |
| *CjAG2* | | *Cryptomeria japonica* | XM_058001897.2 / NC_081407.1 Chromosome 3: 701461423-701485693 (- strand) |
| *CjAG3* | | *Cryptomeria japonica* | XM_058001865.2 / NC_081407.1 Chromosome 3: 699084187-699192766 (- strand) |

| | | | |
|---|---|---|---|
| *SgAG1* | | *Sequoiadendron giganteum* | CM021102.1 Chromosome 3: 306473424-306537103 (- strand) |
| *SgAG2* | | *Sequoiadendron giganteum* | CM021102.1 Chromosome 3: 309520027-309565885 (+ strand) |
| *SgAG3* | | *Sequoiadendron giganteum* | CM021102.1 Chromosome 3: 312155930-312257328 (+ strand) |
| *CsAG1* | | *Cupressus sempervirens* | CM053785.1 Chromosome 3: 749716187-749796384 (+ strand) |
| *CsAG2* | | *Cupressus sempervirens* | CM053785.1 Chromosome 3: 746854424-746924622 (- strand) |
| *CsAG3* | | *Cupressus sempervirens* | CM053785.1 Chromosome 3: 742960508-743061590 (- strand) |
| *CoAG1* | | *Chamaecyparis obtusa* | BSBK01003392.1 COB1p03392: 727913-779915 (- strand) |
| *CoAG2* | | *Chamaecyparis obtusa* | BSBK01002658.1 COB1p02658: 46429-76029 (+ strand) |
| *CoAG3* | | *Chamaecyparis obtusa* | BSBK01005049.1 COB1p05049: 425493-476902 (+ strand) |
| *TpAG1* | | *Thuja plicata* | MU225471.1 scaffold29379416: 839521-911146 (- strand) |
| *TpAG2* | | *Thuja plicata* | MU227659.1 scaffold29381604: 318394-401305 (+ strand) |
| *TpAG3* | | *Thuja plicata* | MU225163.1 scaffold29379108: 289493-381711 (- strand) |
| *NnAG* | | *Nageia nagi* | JBITNC010002277.1 SC2282: 135900067-135941681 (+ strand) |
| *WnAG* | | *Wollemia nobilis* | JAQPPZ010017076.1 scaffold_24681: 2324241-2381651 (- strand) |
| *TgAG1* | | *Torreya grandis* | JARHVA010005221.1 contig_5216: 893329-975986 (+ strand) |
| *TgAG2* | | *Torreya grandis* | JARHVA010001371.1 contig_1370: 3242024-3250206 (- strand) |
| *TgAG3* | | *Torreya grandis* | JARHVA010010244.1 contig_10238: 14618-72506 (+ strand) |
| *TcAG1* | | *Taxus chinensis* | CM034146.1 Chromosome 1: 645483252-645630336 (- strand) |
| *TcAG2A* | | *Taxus chinensis* | CM034146.1 Chromosome 1: 650341172-650390395 (+ strand) |
| *TcAG2B* | | *Taxus chinensis* | CM034146.1 Chromosome 1: 651182616-651210450 (+ strand) |
| *TcAG3* | | *Taxus chinensis* | CM034146.1 Chromosome 1: 653970957-654123526 (- strand) |
| *TwAG1* | | *Taxus wallichiana* | CM031239.1 Chromosome 7: 273761501-273911911 (- strand) |
| *TwAG2A*[c] | | *Taxus wallichiana* | CM031239.1 Chromosome 7: 278925998-278975502 (+ strand) |
| *TwAG2B*[c] | | *Taxus wallichiana* | CM031239.1 Chromosome 7: 279785031-279812823 (+ strand) |
| *TwAG3* | | *Taxus wallichiana* | CM031239.1 Chromosome 7: 282787577-282925610 (- strand) |
| *LkAG* | | *Larix kaempferi* | BSBM01000120.1 LKA1p00120: 6275478-6389590 (- strand) |

| | | | |
|---|---|---|---|
| *PalbAG* | | *Pinus albicaulis* | CM076580.1: 943550943-943660046 (+ strand) |
| *PtabAG* | | *Pinus tabuliformis* | CM062811.1 Chromosome 3: 310297447-310433553 (- strand) |
| *PtaeAG* | *PtDAL2* | *Pinus taeda* | APFE031103740.1 scaffold205186: 31718-175332 (+ strand) |
| *PmAG* | | *Pseudotsuga menziesii* | LPNX011162981.1 jcf7190000027815: 199176-323348 (- strand) |
| *CpAG* | | *Cycas panzhihuaensis* | CM041469.1 Chromosome 5: 401673882-401727852 (+ strand) |
| *AcAG2* | | *Athrotaxis cupressoides* | OneKP scaffold-XIRK-2009770: 239-943 (- strand) |
| *AsAG1* | | *Austrotaxus spicata* | OneKP scaffold-BTTS-2011270: 360-1031 (+ strand) |
| *CdAG2* | | *Cupressus dupreziana* | OneKP scaffold-QNGJ-2070995: 82-780 (- strand) |
| *ChAG1* | | *Cephalotaxus harringtonia* | OneKP scaffold-WYAJ-2089989: 465-1136 (+ strand) |
| *ChAG3*[d] | | *Cephalotaxus harringtonia* | OneKP scaffold-WYAJ-2008272: 141-819 (+ strand) |
| *DcAG* | | *Dacrycarpus compactus* | OneKP scaffold-FMWZ-2046394: 1-324 (- strand) |
| *GpAG1* | | *Glyptostrobus pensilis* | OneKP scaffold-OXGJ-2010405: 88-762 (+ strand) |
| *GpAG2* | | *Glyptostrobus pensilis* | OneKP scaffold-OXGJ-2044691: 268-975 (- strand) |
| *MdAG1* | | *Microbiota decussata* | OneKP scaffold-XQSG-2051433: 2-565 (+ strand) |
| *MdAG2* | | *Microbiota decussata* | OneKP scaffold-XQSG-2052351: 227-913 (- strand) |
| *MdAG3* | | *Microbiota decussata* | OneKP scaffold-XQSG-2053285: 146-805 (+ strand) |
| *MtAG* | | *Microcachrys tetragona* | OneKP scaffold-MHGD-2071655: 272-937 (- strand) |
| *NpAG1* | | *Neocallitropsis pancheri* | OneKP scaffold-JDQB-2008203: 3-548 (+ strand) |
| *NpAG2* | | *Neocallitropsis pancheri* | OneKP scaffold-JDQB-2008192: 370-1071 (- strand) |
| *PcAG1*[d] | | *Pseudotaxus chienii* | OneKP scaffold-YLPM-2012018: 34-745 (- strand) |
| *PoAG1* | | *Platycladus orientalis* | OneKP scaffold-BUWV-2007345: 10-681 (- strand) |
| *PoAG2*[e] | | *Platycladus orientalis* | OneKP scaffold-BUWV-2001427: 1-494 (- strand) |
| *PpAG1* | | *Papuacedrus papuana* | OneKP scaffold-OVIJ-2083454: 317-985 (- strand) |
| *PpAG2* | | *Papuacedrus papuana* | OneKP scaffold-OVIJ-2082882: 268-969 (+ strand) |
| *PpAG3* | | *Papuacedrus papuana* | OneKP scaffold-OVIJ-2005716: 400-1071 (- strand) |
| *PustAG* | | *Parasitaxus usta* | OneKP scaffold-JZVE-2036946: 224-906 (+ strand) |
| *PuviAG3* | | *Pilgerodendron uviferum* | OneKP scaffold-ETCJ-2056171: 3-422 (+ strand) |

| | | | |
|---|---|---|---|
| *SaAG* | | *Sundacarpus amarus* | OneKP scaffold-KLGF-2089070: 1-363 (- strand) |
| *TdAG2* | | *Thujopsis dolabrata* | OneKP scaffold-NKIN-2008827: 342-1040 (- strand) |
| *TdAG3* | | *Thujopsis dolabrata* | OneKP scaffold-NKIN-2048040: 161-820 (+ strand) |
| *TnAG1*[d] | | *Torreya nucifera* | OneKP scaffold-HQOM-2014418: 367-1030 (+ strand) |
| *TnAG3* | | *Torreya nucifera* | OneKP scaffold-HQOM-2006240: 58-666 (+ strand) |
| *TtAG3* | | *Torreya taxifolia* | OneKP scaffold-EFMS-2077547: 3-608 (- strand) |
| *AmAG* | | *Agathis macrophylla* | GKZN01020124.1 |
| *AmAGL*[g] | | *Agathis macrophylla* | GKZN01016303.1 |
| *CnAG1* | | *Callitropsis nooktanensis* | GKZO01019107.1 |
| *CnAG2* | | *Callitropsis nooktanensis* | GKZO01023595.1 |
| *CnAG3* | | *Callitropsis nooktanensis* | GKZO01011510.1 |
| *CepsinAG2* | | *Cephalotaxus sinensis* | GKZP01008021.1 |
| *CepsinAG3* | | *Cephalotaxus sinensis* | GKZP01008022.1 |
| *SvAG* | | *Sciadopitys verticillata* | GKZY01002684.1 |
| *TbAG1* | | *Taxus baccata* | GLAA01028133.1 |
| *TbAG2B*[f] | | *Taxus baccata* | GLAA01013640.1 |
| *TbAGL*[g] | | *Taxus baccata* | GLAA01002276.1 |
| *TsucanAG* | | *Tsuga canadensis* | GLAB01018135.1 |

[a] *TdisMADS19* appears to be a chimeric sequence since it has two putative M domains. For the alignment, M domain toward the C terminus was used.

[b] The *Arabidopsis thaliana* Type I MADS-box genes (*DIA*, *AGL62*, and *AGL23*) with sequences downstream of the M domain removed to serve as the outgroup.

[c] *TwAG2A* and *TwAG2B* contain indels that cause frameshifts in their coding sequences. We are uncertain whether these are real or sequencing/assembly errors given the sequence conservation in the rest of the coding sequence. Since these genes are only incorporated for phylogenetic analysis, these frameshifts were adjusted based on sequences of *TcAG2A* and *TcAG2B* prior to alignment.

[d] *ChAG3* has an insertion (N) at position 598 in the CDS that causes a frameshift. *PcAG1* has an insertion (N) at position 201 in the CDS (after reverse complementation) that causes a frameshift. *TnAG1* has an insertion (N) at position 569 in the CDS that causes a frameshift. These insertions were deleted prior to alignment.

[e] *PoAG2* has a deletion that causes a frameshift. An N was inserted at position 130 in the CDS (after reverse complementation) to accommodate the frameshift.

[f] First 84 nucleotides of *TbAG2B* were deleted due to lack of similarity to M domain of other sequences.

[g] Gene is a member of the long-branch *AG*-like clade.

**Table S4.** Information for RNAscope *in situ* hybridization probes.

| | |
|---|---|
| **Probe for *TdisAG1*** | |
| Submitted database(s) | *Taxodium distichum* transcriptome assembly (this study) |
| Probe name (per ACDBio) | Tdis-MADS12 |
| # of ZZ pairs | 10 |
| Target region of CDS | Nucleotides 6-671 |
| Cross detections | Isoforms of *TdisAG1* (labeled TRINITY_DN4709_c0_g1 in transcriptome assembly) |
| Notes | The *T. distichum* transcriptome was sequenced, assembled, and submitted to ACDBio prior to the publication of the genome (32). The RNAscope Probe Design Team confirmed that the probes only detect transcripts produced by the *TdisAG1* locus in the genome. |
| **Probe for *TdisAG2*** | |
| Submitted database(s) | *Taxodium distichum* transcriptome assembly (this study) and genome (32) |
| Probe name (per ACDBio) | Tdis-MADS5 |
| # of ZZ pairs | 10 |
| Target region of CDS | Nucleotides 16-729 |
| Cross detections | None |
| Notes | The *T. distichum* transcriptome was sequenced, assembled, and submitted to ACDBio prior to the publication of the genome (32). The RNAscope Probe Design Team confirmed that the probes only detect transcripts produced by the *TdisAG2* locus in the genome. |
| **Probe for *TdisAG3*** | |
| Submitted database(s) | *Taxodium distichum* transcriptome assembly (this study) and genome (32) |
| Probe name (per ACDBio) | Tdis-MADS30 |
| # of ZZ pairs | 11 |
| Target region of CDS | Nucleotides 49-673 |
| Cross detections | Isoforms of *TdisAG3* (labeled TRINITY_DN18980_c0_g2 in transcriptome assembly) |
| Notes | The *T. distichum* transcriptome was sequenced, assembled, and submitted to ACDBio prior to the publication of the genome (32). The RNAscope Probe Design Team confirmed that the probes only detect transcripts produced by the *TdisAG3* locus in the genome. |
| **Probe for *JvAG1*** | |
| Submitted database(s) | *Juniperus virginiana* transcriptome assembly (this study) |
| Probe name (per ACDBio) | Jv-MADS8 |
| # of ZZ pairs | 10 |
| Target region of CDS | Nucleotides 20-662 |
| Cross detections | None |
| **Probe for *JvAG2*** | |
| Submitted database(s) | *Juniperus virginiana* transcriptome assembly (this study) |
| Probe name (per ACDBio) | Jv-MADS16 |
| # of ZZ pairs | 9 |
| Target region of CDS | Nucleotides 2-692 |
| Cross detections | Isoforms of *JvAG2* (labeled TRINITY_DN10663_c0_g1 in transcriptome assembly) |
| Notes | The ACD Probe Design Team warned us that the probe is short (<10ZZ pairs), so we should ensure that optimal assay conditions are maintained. |
| **Probe for *JvAG3*** | |
| Submitted database(s) | *Juniperus virginiana* transcriptome assembly (this study) |
| Probe name (per ACDBio) | Jv-MADS5 |

| | |
|---|---|
| # of ZZ pairs | 13 |
| Target region of CDS | 4-656 |
| Cross detections | Isoforms of *JvAG3* (labeled TRINITY_DN3832_c0_g1 in transcriptome assembly) and TRINITY_DN155514_c0_g1_i1 |
| Notes | TRINITY_DN155514_c0_g1_i1 and *JvMADS5* BLAST to the same region in the genome of *Cupressus sempervirens* (35) (*Cupressus* and *Juniperus* are sister genera). Thus, TRINITY_DN155514_c0_g1_i1 is likely an isoform of *JvAG3* that did not get assembled properly. |
| **Probe for *MgAG1*** | |
| Submitted database(s) | *Metasequoia glyptostroboides* genome (31) |
| Probe name (per ACDBio) | Mgl-MgAG1 |
| # of ZZ pairs | 12 |
| Target region of CDS | 20-675 |
| Cross detections | None |
| **Probe for *MgAG2*** | |
| Submitted database(s) | *Metasequoia glyptostroboides* genome (31) |
| Probe name (per ACDBio) | Mgl-MgAG2 |
| # of ZZ pairs | 12 |
| Target region of CDS | 3-593 |
| Cross detections | None |
| **Probe for *MgAG3*** | |
| Submitted database(s) | *Metasequoia glyptostroboides* genome (31) |
| Probe name (per ACDBio) | Mgl-MgAG3 |
| # of ZZ pairs | 8 |
| Target region of CDS | 221-663 |
| Cross detections | None |
| Notes | The ACD Probe Design Team warned us that the probe is short (<10ZZ pairs), so we should ensure that optimal assay conditions are maintained. |
| **Probe for *DAL2*** | |
| Submitted database(s) | *Picea abies* transcriptome assembly (11, 12) |
| Probe name (per ACDBio) | Pabi-DAL2 |
| # of ZZ pairs | 20 |
| Target region of transcript | 74-1433 |
| Cross detections | Isoforms of *DAL2* (labeled TRINITY_DN10906_c0_g1 in transcriptome assembly) |

**Table S5.** Results for positive selection from the MEME program in HyPhy. Results from aBSREL not shown since no branches were significant.

| Codon | Domain | α (dS) | β (dN) distribution (rates:weights) | LRT | Adjusted p-value | # of Branches |
|---|---|---|---|---|---|---|
| 50 | M | 2.256 | 0.00/23785.87:0.45/0.55 | 9.062 | 0.0047 | 1 |
| 117 | K | 1.726 | 0.00/10.49:0.01/0.99 | 4.746 | 0.0430 | 1 |
| 118 | K | 1.565 | 0.69/126.03:0.44/0.56 | 10.106 | 0.0028 | 0 |
| 153 | K (K2) | 3.120 | 0.00/5368.94:0.62/0.38 | 13.479 | 0.0005 | 1 |
| 156 | K | 0.000 | 0.00/8.51:0.01/0.99 | 10.309 | 0.0025 | 0 |
| 167 | K (K3) | 0.919 | 0.00/7.20:0.01/0.99 | 4.741 | 0.0431 | 2 |
| 208 | C | 1.153 | 0.00/6.59:0.01/0.99 | 4.661 | 0.0449 | 1 |

**Dataset S1 (separate file).** Length of introns and number of identified transcription factor binding sites for each *AG* paralog of 8 Cupressaceae species.
